# Structural and functional classification of G-quadruplex families within the human genome

**DOI:** 10.1101/2023.02.09.527851

**Authors:** Aryan Neupane, Julia H. Chariker, Eric C. Rouchka

## Abstract

G quadruplexes are short secondary DNA structures located throughout genomic DNA and transcribed RNA. though G4 structures have been shown to form *in vivo*, no current search tools are known to exist to examine these structures based on previously identified G quadruplexes, much less filter them based on similar sequence, structure, and thermodynamic properties. We present a framework for clustering G quadruplex sequences into families using the *CD-HIT, MeShClust and DNACLUST* methods along with a combination of *Starcode* and *BLAST*. Utilizing this framework to filter and annotate clusters, 95 families of G quadruplex sequences were identified within the human genome. Profiles for each family were created using hidden Markov models to allow for identification of additional family members and generate homology probability scores. The thermodynamic folding energy properties, functional annotation of genes associated with the sequences, scores from different prediction algorithms and transcription factor binding and motif to the G4 region for the sequences within a family were used to annotate and compare the diversity within and across clusters. The resulting set of G quadruplex families can be used to further understand how different regions of the genome are regulated by factors targeting specific structures common to members of a specific cluster.

## INTRODUCTION

G-quadruplexes are stranded secondary structures of nucleic acids rich in guanine containing four runs of at least three guanines. These runs are separated by short loops, typically 2-7 nucleotides in length, which can potentially fold into an intramolecular G-quadruplex structure. The tetrad guanine structure is stacked on top of each other and held together by mixed loops of DNA forming Hoogsteen base pairing giving a four-stranded structure with nucleobases on the inside and a sugar phosphate backbone on the outside (Figures 1A and 1B). Metal ions (typically K+ or Na+) sitting internally to the Hoogsteen bases stabilize the base pairing. Stacking occurs through the O6 atoms of guanines facing the center creating a tubular space able to function as an ion channel. The presence of a metal cation in this channel allows for interaction with the eight O6 atoms of the guanine quartet.

**Figure 1.**
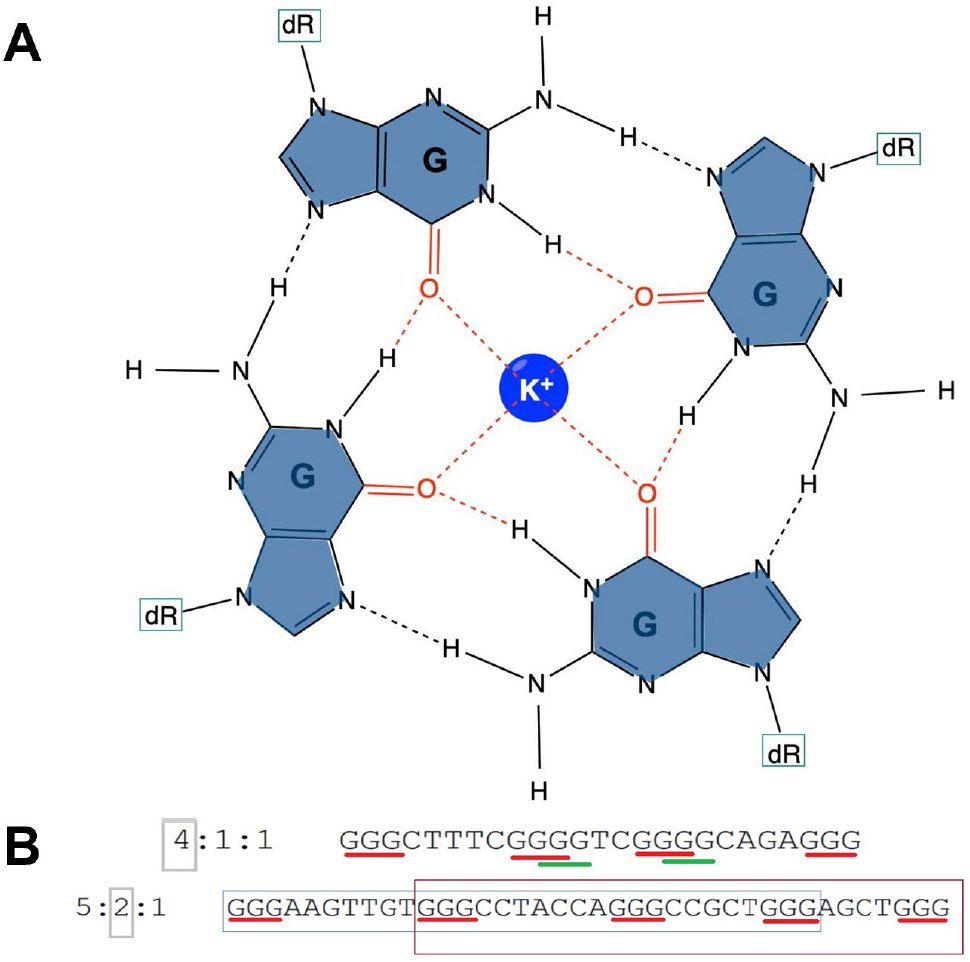
G quadruplex structure. (A) G-tetrad structure forming G quadruplexes. Hydrogen bonds between the guanine from different tetrads form a planar ring (B) Sequence of G4 with multiple guanine tetrads. Here, 4:1:1 and 5:2:1 refers to the result from Quadparser separated out as the number of tetrads: Total G4 sequences: Non overlapping G4 sequences.

### Roles of G-quadruplexes

Over the past three decades, guanine rich quadruplex sequences have been implicated as key structural regulators of gene expression, cellular differentiation and transcription factors and their cell line and tissue specificity (1). Similarly, elevated levels of G quadruplexes have been identified across cancer tissues including breast (2), stomach (3), liver (4), as well as neurodegenerative diseases (5). Computational analyses of G quadruplex patterns have identified the prevalence of G quadruplexes in oncogenic promoters, introns, splice sites, intergenic and telomeric ends. Initially the secondary structures were thought to act as a physical obstacle to RNA polymerase for transcription as identified through G4 specific antibodies (6,7) and chemical probing (8,9). Further evidence suggests the varied tissue specific functionality of these structures are affected by the cross talk of additional transcription factors (10), proteins and physiological conditions. Additionally, G4 structures have a role in genomic instability and are associated with higher rates of double strand breakage in nucleosome depleted regions of highly expressed cancer genes. High- and low-density bands of G4 across both chromosomal strands have been observed showcasing a role of G quadruplex in pairing of homologous chromosomes during meiosis (11). Further, recent evidence shows that G4 formation is highest during DNA replication at the S phase and lowest during G2 and M phase, consistent with phases of transcription, replication, and chromatin accessibility (12).

### Characteristics of G4s

Sequence characteristics such as sequence length (13), base composition (14,15) and loop length (16–20) are important parameters for defining the secondary structure and stability of G quadruplexes. Molecular dynamics show that telomeric G4 repeats (TTAGGG) in the presence of a K+ cation form a structure with three single nucleotide loops in a parallel fashion. Increasing the loop length by a single base causes the sequences to adopt a mixture of parallel and antiparallel folded structures (21). The conformation and stability of G quadruplexes has been used to study to effect of transcription factor binding and altered mRNA expression of several genes. Examples include nucleolin (22) and Ewing’s Sarcoma proteins (23) which preferentially bind to structures with longer loop length. Computationally, G4s are defined by the pattern G_*x*_N_1-7_G_*x*_N_1-7_G_*x*_N_1-7_G_*x*_ where x ≥3 (length of G-guanine repeats) separated by loops of any base composition with a length 1-7 bases. This pattern is the basis for regular expression-based tools such as Quadparser (24) and QGRSmapper (25). With experimental data, it is known that different intermolecular structure, long loops, and non-canonical structures with G tracts containing two guanines exist (26–29). Methods such as G4screener (30), PQSfinder (31) and G4Catchall (32) allows the search of G quadruplexes for variable quartet and larger loop sequences. G4Hunter (33) provides a score for guanine skewness which is based on predefined values, with a score based on the number of consecutive Gs. G4RNAscreener (30) uses a machine learning algorithm trained with experimental RNA sequences from the G4RNA database (34) and incorporates a threshold using metrics from tools such as G4Hunter score (33), cG/cCscore, and G4 Neural Network score for G4 prediction. RNAfold (35) has an option to predict the thermodynamic parameters for G quadruplex formation. DSSR (36) and ElTetrado (37) use the tertiary structure of each G quadruplex for annotating and classifying different base pairs and tetrad structures. 3D-NuS (38) allows visualization of 3D DNA structures including duplex, triplex, and quadruplexes. 3D NuS visualizes the G quadruplex structure and its strand orientation, loops and G quartets based on the energy minimization of G4 structures using experimental data.

G4 structure was found to be evolutionarily conserved in seven yeast species (39). While G quadruplex regions are significantly enriched in regulatory regions of eukaryotes, short loops of G4 are conserved in different species. Protozoa and fungus have limited diversity of G4 while an increase in diversity has been observed across invertebrates and vertebrates (40). However, the evolutionary mechanism for this structure or the relationship of these structures at an evolutionary scale is not known.

Sequencing read fragments utilizing a customized approach that introduces stabilizing and destabilizing conditions (K+, Li+, PDS) allows for high throughput sequencing of G4 locations (41,42) with a method known as G4 seq. Versions of this method have been used to identify 1,420,841 G quadruplexes in 12 species. Using a similar method, 161 and 168 G4 sites were identified in the genomes of *Pseudomonas* (43) and *Escherichia* (44), respectively.

Over 100,000 G4 sequences have been mapped *in vivo* to the human genome. Several proteins such as FUS, TAF15, TARDBP, PCBP1 have been determined to be enriched at G4 loci using artificial G4 binding (45). SP2, a transcription factor (TF) encoded by a subfamily of the Sp/XKLF family, is a sequence specific TF that has a strong association with G quadruplex affinity. SP2 binds to the CCAAT motif independent of the zinc finger domain necessary for binding to GC rich motifs (46). It was shown *in vitro* that the SP1 TF was able to bind to a DNA sequence lacking the consensus motif and was able to form G quadruplex sequences (47). Luciferase expression studies show sequences of G4s in the KIT promoter mutated through site directed mutagenesis were able to create a modulation (on/off) system for KIT expression through SP1 binding (48). Additionally, G quadruplex structures can bind to G quadruplex sites in other promoter locations (49) mediating cis (50) and trans (51) acting regulation of transcriptional and translational processes respectively, implying that G quadruplex sequence and structural diversity is a key factor for biological functions.

### G4 families

Previously, a small family of G quadruplexes labelled Pu27 was identified based on sequence homology (49). The parent G quadruplex is a 27 nucleotide (nt) G4 formed in the nuclease hypersensitive element (NHE) region of the c-MYC promoter associated with different forms of cancer, and predominantly involved in the regulation of expression of c-MYC gene (52). c-MYC is an oncogene that regulates genes in cell cycle and molecular metabolisms. Rezzoug et al identified seventeen potential G quadruplex forming sequences homologous to the Pu27 G4 which has been shown to selectively bind to the NHE region of c-MYC promoter (49). In addition, G4 regions regulating VEGF genes have been shown to have an additional G-tract to act as a spare tire for formation of G quadruplex sequence upon oxidative damage upon the guanine tracts (53). Similar sequences have been identified for c-MYC, KRAS (54), BCL2 (55), HIF-1α and RET genes. This highlights the presence of sequence specific G quadruplexes able to form, bind and regulate gene expression. Further, over the past decade numerous G quadruplex stabilizing and destabilizing ligands have been identified that recognize and interact selectively to these G4 sequences. Different classes of these heteroaromatic polycyclic, macrocyclic and aromatic compounds have been designed to target the diversity of G4 structure. The subtle differences in grooves, loop composition and loop length allow for structure variability in these sequences. DNA aptamers that can form G4 are used for binding nucleolin (56). More than 50 transcription factors with overlapping binding sites to G4 region have been identified (1,57). Folding, misfolding and unfolding of G4 structures have been implicated in different biological processes (58,59).

### Detection of G4 families

The prediction of G quadruplexes across genomes can be useful to identify the location of similarly structured G quadruplexes, which can in turn be used to develop profiles of independent families based on conservation of a variety of factors. We present a framework to predict G quadruplex sequences and identify similar sequences using trained profile hidden Markov models (HMMs) (60). We identify pG4 sequences across human genome, cluster these sequences using sequence clustering tools, CD-hit (61), MeShClust (62) and DNACLUST (63) as well as starcode (64) and BLAST (65). These approaches utilize average weighted clustering to identify the quartet and loop patterns, and further train HMM models using these clusters for creation of families. Despite the short length of G quadruplex sequences, position dependent insertion and deletion within loops offers insight into the loop characteristics.

## MATERIAL AND METHODS

### Dataset preparation

Since there are no current families or experimental similarities of G4 structure, we start with putative G4s and apply sequence-based methods for clustering. Later, these clusters are used as initial seeds for identifying G quadruplexes in experimental datasets. Initially, we focused on the G4s identified from Quadparser (24) on the human GRCh38 genome. The following process is followed for all groups of sequences based on the number of GGG tetrads (Figure 2A).

**Figure 2.**
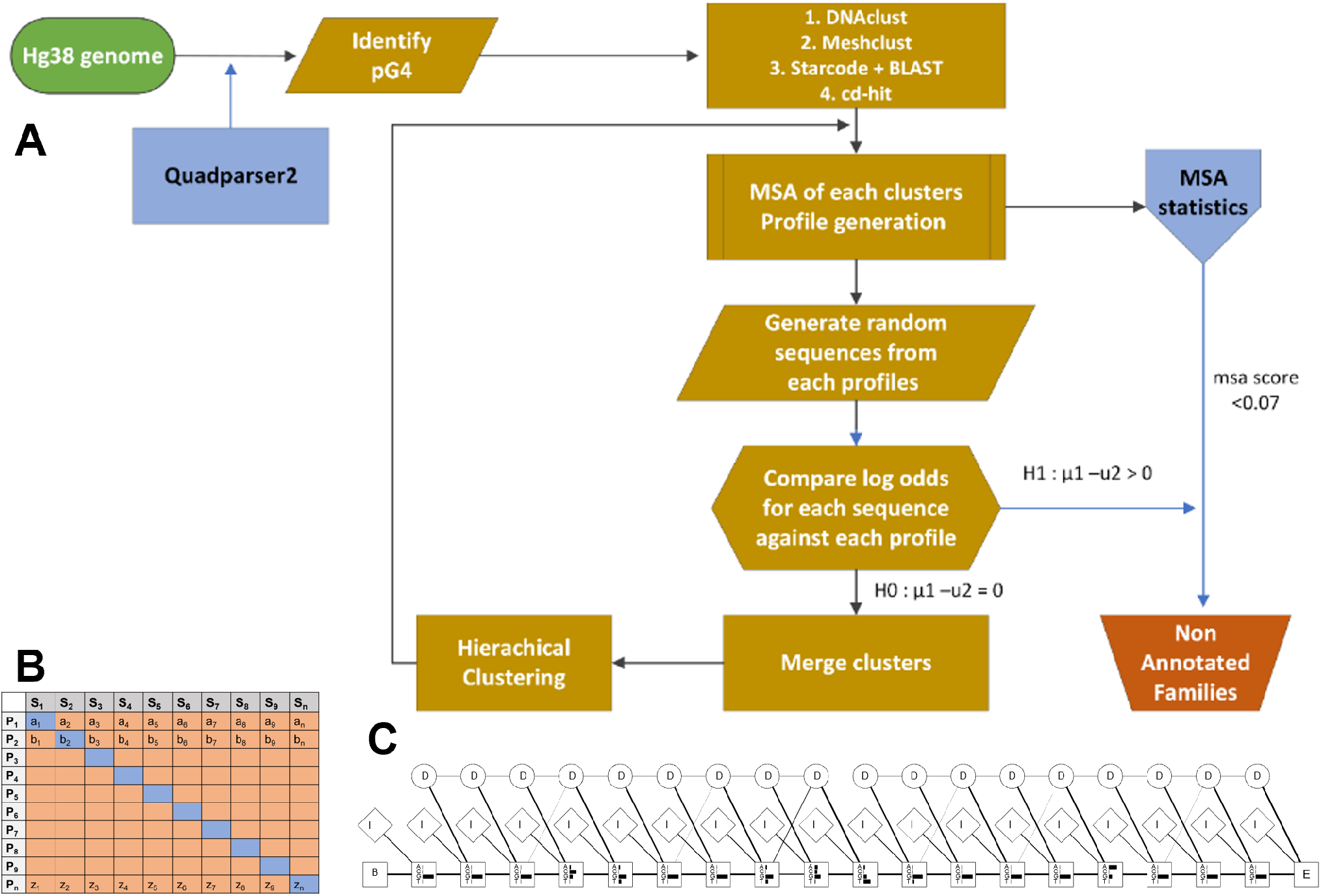
Process for identifying and characterizing G quadruplex families. (A) Workflow diagram for identifying distinct G quadruplex families. (B) Process for identifying appropriate profile for a specific family. In this case, S1,…,Sn represents the list of sequences generated from HMM profile P1,…,Pn respectively. We compare the average log odds for input S1 over profile P1 … Pn and recursively measure for all the profiles. For each row, the diagonal element is compared with non-diagonal values(log-odds) using a Wilcoxon rank sum test with null and alternate hypothesis, H0: T1 –T2 = 0, H1: T1 –T2 > 0. (C) Profile HMM derived from a selected G4 alignment. Match states are represented as rectangles with four residue emission probabilities indicated as black bars, insert states (I) as diamond, and delete states as circle. The start and end states are B (begin) and E (end) respectively. Delete states are silent states with no emission probabilities and weighed lines represent the transition probabilities between states.

1. CD-hit, MeShClust, and DNAclust and a combination of Starcode and BLAST with hierarchical clustering are utilized for the initial clustering of G-quadruplex sequences.
2. Steps (3)-(7) are repeated separately for each clustering method.
3. A multiple sequence alignment (MSA) of each cluster of sequences is carried out in R using the DECIPHER package (66). The StaggerAlignment and AdjustAlignment functions are used to separate regions of alignment and gaps are shifted to improve the alignment.
4. Clusters with fewer than four sequences are filtered out. An MSA score for each cluster is calculated as the average number of gaps in each column of an alignment divided by the length using MStatX (67).
5. Each alignment is trained as a model profile HMM using HMMER 3.0 (68) and the aphid package (69) in R version 3.4.1 independently. The transition and emission probability matrices are estimated based on the plan7 PHMM model based on Durbin (60). An example of a profile HMM stating match, insert and delete state is shown in Figure 2B. There are seven outgoing transitions based on the match, insert and delete states, i.e. I_n_ → I_n_, M_n_→ I_n_, M_n_ → M_n_+1, M_n_ → D_n_+1; D_n_ → M_n_+1, D_n_ → D_n_+1; I_n_ → M_n_+1 where n represents each position of the alignment (except the final position). The observed counts of emissions and state transitions are converted into probabilities.
6. Due to the highly repetitive guanine stretches and the short sequence length of G4 sequences, most of the clusters suffer redundancy (overfitting) in predicting similar sequences. 20 sequences are generated from each profile HMM with an average length of the sequences used to create the profile. The sequences are used as input for all the profiles and the log-odds scores are generated using the forward algorithm.
7. A pairwise Wilcoxon rank sum test is carried out to compare each profile using the log-odds between the profile HMM through which the sequences were generated and all other profiles (Figure 2C). If a profile is diverse (p value < 0.05) against all profiles compared, has a probability of 0.90 for the tested sequences, and has a gap score less than a threshold of 0.10, the profile is saved as a family. For the sequences that are non-significant (p value > 0.05) the sequences are input to the MSA and are merged and/or clustered using agglomerative clustering. Alignments with a gap score of 0.6 after merging are filtered. The process is iterated for a maximum of n=100 times.
8. The group of sequences obtained from all the methods is combined and checked for redundancy using a modification of step (7) utilizing a threshold score of log-odds 6 and Akaike weight of 0.7 for identifying the final families, which are additionally manually checked and filtered.
9. The alignment and profile HMM are manually verified resulting in 95 clusters referred to as families. Experimentally validated G quadruplexes were obtained from processed peaks mapped to hg19 from GEO, accession GSE63874 (41) using bedtools (70) and quadparser2 after conversion to human genome hg38 coordinates by liftover. The models are used as a trained classifier for identifying additional sequences. G4 sequences from experimental G4 seq was tested against the cluster HMMs. The likelihood that a query sequence fits the model of an individual family is calculated using the forward algorithm (71) and the normalized Akaike weights (72,73) is calculated. The maximum Akaike weight of the query given a particular model is selected as the nearest family of the query sequence. The families are manually verified and the variability of sequences in the families is further analyzed based on annotation of the G4, thermodynamic scores (folding energy), G4hunter scores and literature. The steps below highlight the method for the combination of Starcode and BLAST with hierarchical clustering.

a. Levenshtein distance is used to identify the nearest group of sequences which are then filtered based on the length of the sequence and the number of G tetrads. Starcode (64) utilizes a modified Needleman-Wunsch dynamic programming approach known as the poucet
b. algorithm for determining the initial and nearest groups of sequences. Sequences below a fixed Levenshtein score are used to identify the groups and each group is filtered by length of the sequence and loop sequence content. Using specific Levenshtein distance as a constraint through this algorithm, one or two nucleotide mismatches can be identified in short DNA sequences.
c. The remaining sequences from step (1) that are not in any group are passed through BLAST for pairwise all vs all BLAST. −log(E value) is used as the similarity metric.
d. Hierarchical clustering is applied comparing the agglomerative, Ward, complete, and divisive methods of clustering. The number of clusters is calculated based on the sum of the within-cluster inertia. The optimal number of clusters is the maximum difference from two successive clusters between the groups, i.e. max (I_m_/I_m+1_). The mode of the number of clusters was selected as the optimal cluster.
e. Pairwise alignment of sequences of individual clusters obtained from steps (a) and (c) is carried out using the Pairwisealignment function in the Biostrings (74) package. Hierarchical clustering of the sequences is performed based on the pairwise distance. Consensus of Silhouette (75), Frey index, Macclain Index, Cindex, and Dunn index were used for identifying the optimal number of clusters. The metrics are calculated using the NbClust package (76) in R.

## RESULTS

In the preliminary step, a combination of Starcode and BLAST was used with hierarchical clustering to identify 2,717 clusters of G quadruplexes with 29,112 sequences. Using DNACLUST, 587 clusters with 4,664 sequences were identified. A total of 786 clusters with 6,335 sequences were identified with Cdhit with a k-mer of 8. MeShClust with an identity threshold of 90% and k-mer size of 9 was able to identify 508 clusters. Any clusters with fewer than four sequences were discarded. The two largest clusters had 1,720 and 1,410 sequences, respectively. The overall summary of the clustering results is provided in Table 1.

**Table 1.** Cluster summary based on different clustering techniques.

| Method | Number of sequences | No of clusters | No. of sequences in 2 largest clusters | HMM clusters (sequences) | HMM families, 1<sup>st</sup> iteration (sequences) | Final families selected (# of sequences) |
| --- | --- | --- | --- | --- | --- | --- |
| Starcode + BLAST with hierarchical clustering | 29,112 | 2,717 | 419, 323 | 95 (842) |  |  |
| DNAClust | 9,610 (4,664) | 587 | 142, 126 | 31 (1,165) |  |  |
| Cd-hit(kmer 8) | 6,335 | 786 | 182, 115 | 30 (389) |  |  |
| Meshclust | 14,222 | 508 | 1,720 1,410 | 72 (1,843) |  |  |
| **Total** |  |  |  |  | 220 (3,888) | 95 (2,174) |

The resulting HMMs for the identified clusters were utilized to predict additional G quadruplex sequences. In addition, the MSA was used to detect transcription factor site motifs found within each family. The G4 families suffered from redundancy of motifs because of the high percentage of guanine bases. To identify unique motifs, a pipeline was created to merge and re-cluster the families. Overall, the Starcode and BLAST pipeline identified 95 clusters of G-quadruplex genomic DNA sequences. The MeShClust pipeline identified 72 clusters, while DNACLUST And CD-HIT identified 31 and 30, respectively. The final iteration of the clustering and merging sequences across profiles from the various clustering approaches resulted in 98 distinct families.

### G quadruplex families

The resulting 95 families were created from 1,739 distinct individual G4s identified from 2,145 distinct regions of the hg38 human genome. Given the short sequence length and guanine composition, many of the G4 sequences are not unique. One of the largest families identified, Family 23 is comprised of 163 regions with 118 distinct G4s occurring over 122 genes (Supplemental Table 1). Similarly, Family 79 has 130 regions with 99 distinct G4s occurring over 128 genes distributed across all chromosomes (Supplemental Table 2). We identified multiple sequence repeats capable of forming multiple G4 structures with different conformation in Families 46, 62, 88, 89 and 90 based on the available guanines (Supplemental Table 3). Smaller families 2 and 3 have 7 and 6 distinct sequences occurring in proximity to 8 and 7 genes respectively (Supplemental Tables 4 and 5). A summary of the predicted of G4 sequence families is present in Table 2.

**Table 2.**
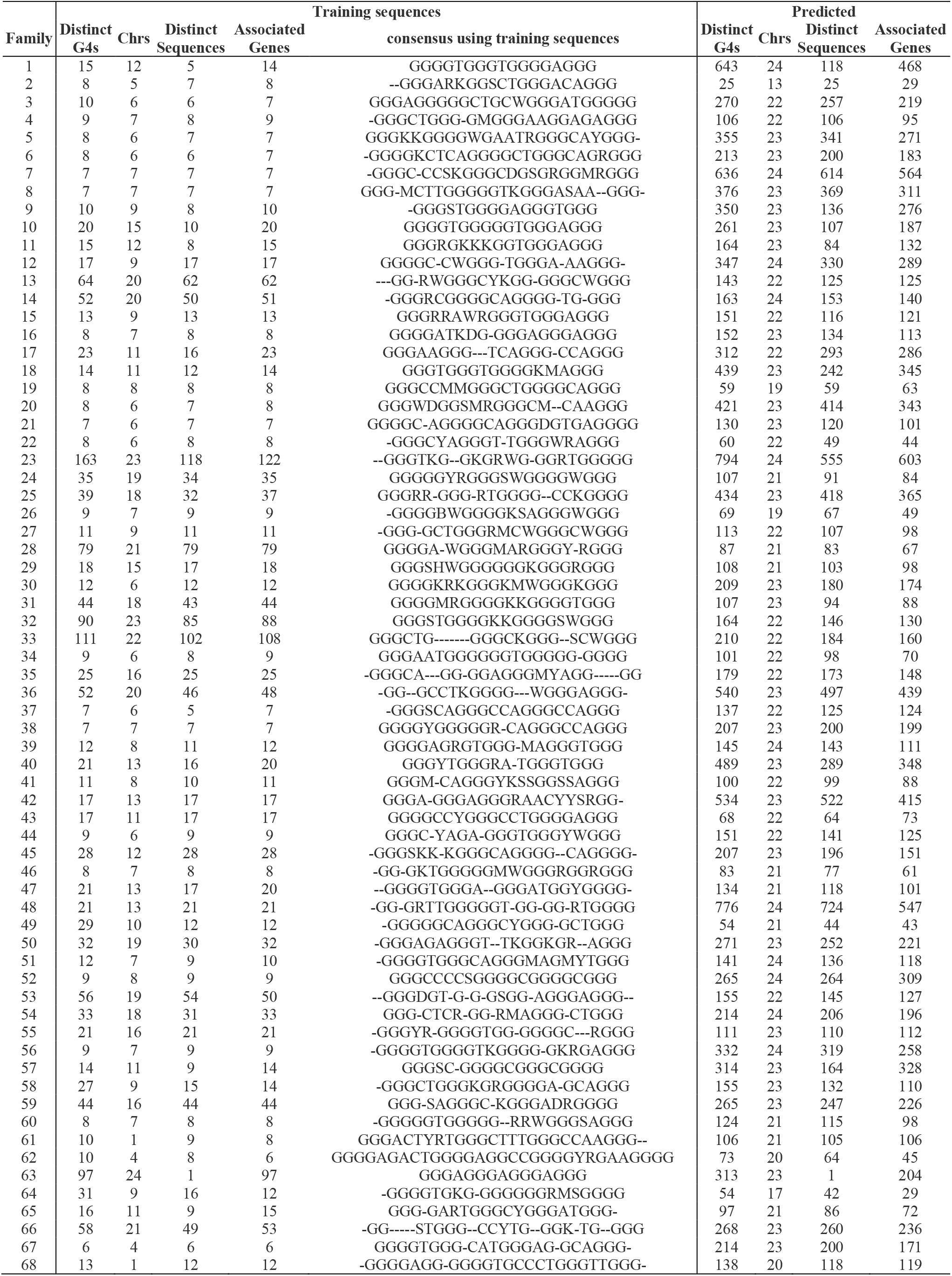
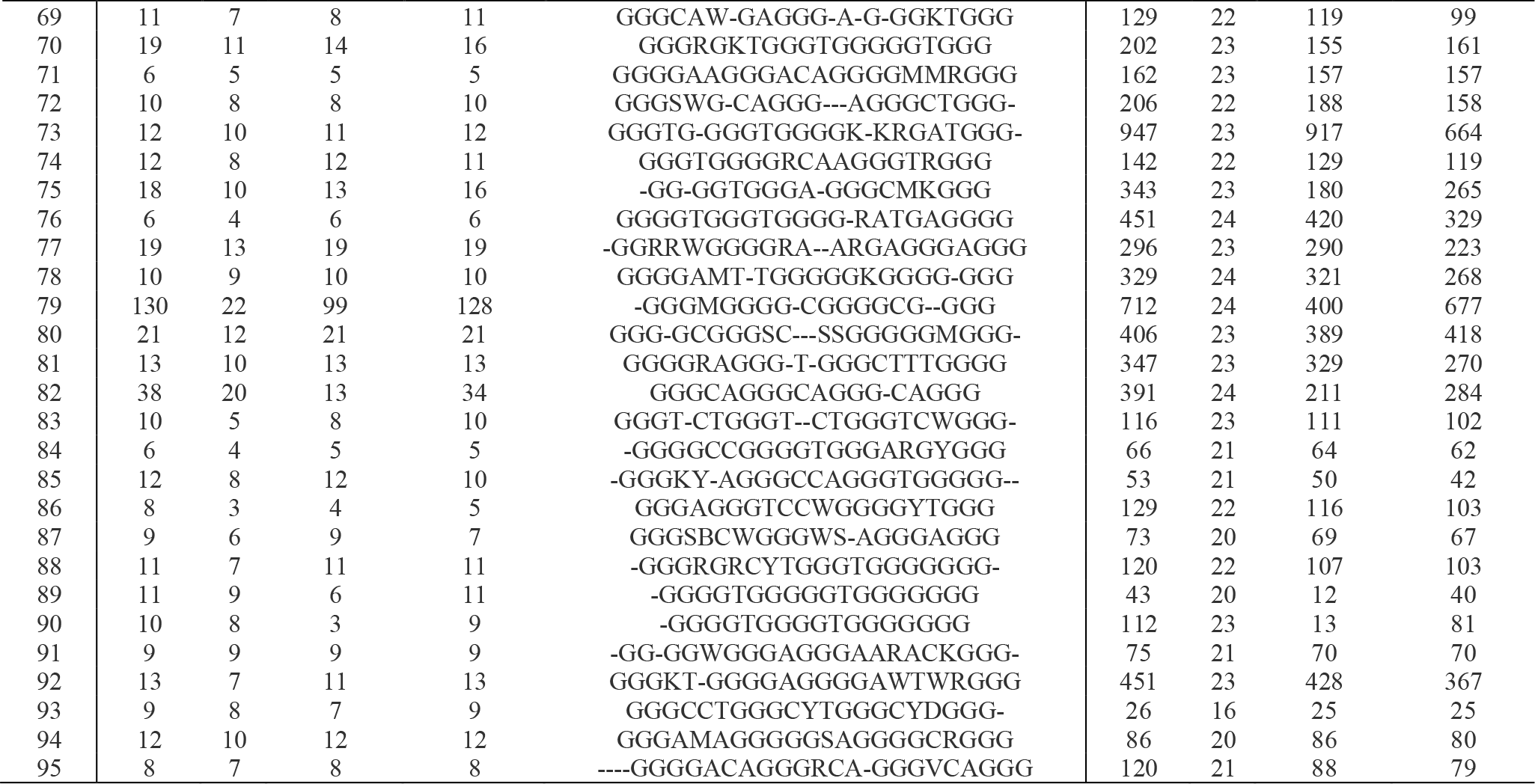
Summary of count of G4 sequences identified using predictive models pHMM across different clusters, genes, and chromosomes.

We analyzed the clusters for their sequence characteristics, functional annotation, and structural structures as presented below. We highlight some of the clusters that have strong biological significance with related biological and molecular processes, including Family 4 (Supplemental Table 6), Family 32 (Supplemental Table 7), Family 75 (Supplemental Table 8), and Family 80 (Supplemental Table 9).

### Categorical enrichment of select families

Family 4 consists of nine sequences distributing over nine genes and seven chromosomes. Figure 3 illustrates the dot bracket notation of the consensus of the family along with thermodynamic characteristics. While this family is relatively small, the associated genes are related, showing an enrichment of terms related to the neural cells (e.g. glia guided migration, synapse assembly, dendritic spine development, and gliogenesis) (Supplemental Figure 1, Supplemental Table 10).

**Figure 3.**
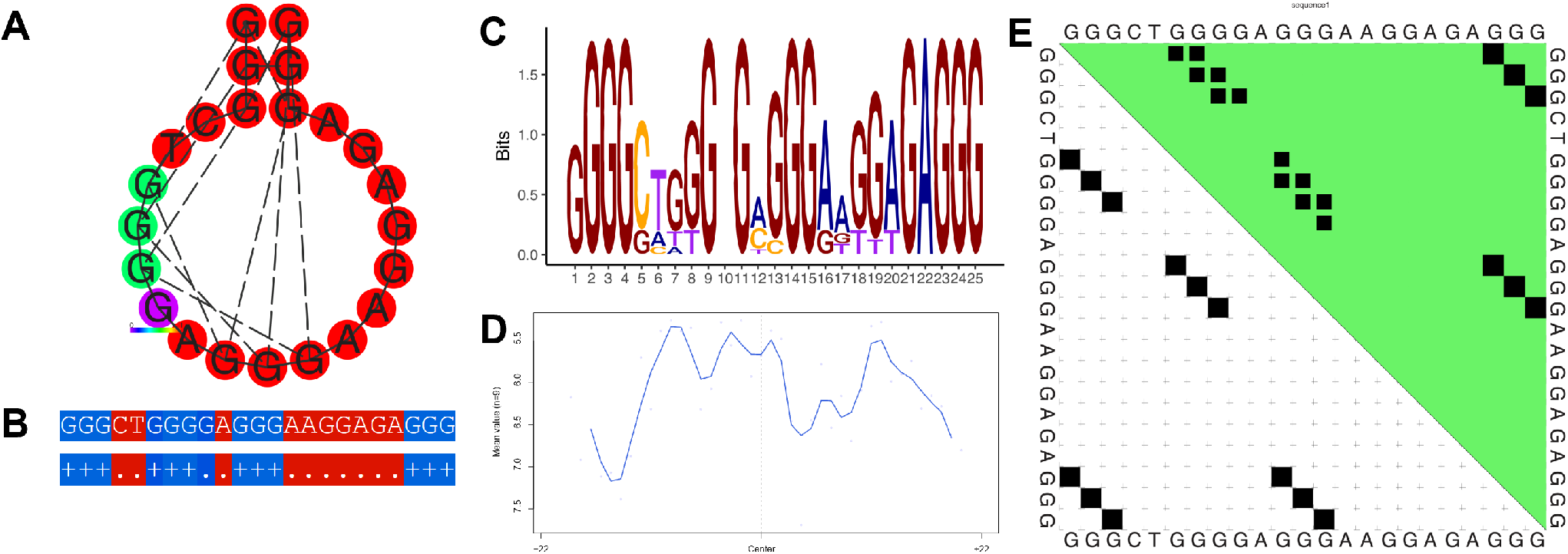
Thermodynamic properties for Family 4. (A) Centroid secondary structure with a minimum free energy of −9.64 kcal/mol using the consensus sequence of the family. (B) Dot-bracket notation showing the secondary structure. (C) Sequence logo representing the per base information content. (D) Electrostatic potential generated from all the sequences of the family using 10 flanking bases on either side of the identified G4. (E) Dot plot showing the substructures with the highest probabilities.

Family 32 contains 90 G4 sequences annotated with 85 genes. The thermodynamic properties are illustrated in Figure 4. The genes associated with Family 32 G4s are enriched for cellular organization (e.g. positive regulation of cell projection organization and positive regulation of cellular component organization), axonal development (e.g. neuron projection guidance, axon guidance), mitochondrial localization (e.g. regulation of protein targeting to mitochondrion and regulation of establishment of protein localization to mitochondrion) and size regulation (e.g. regulation of anatomical structure size and regulation of cell size) (Supplemental Figure 7, Supplemental Table 11).

**Figure 4.**
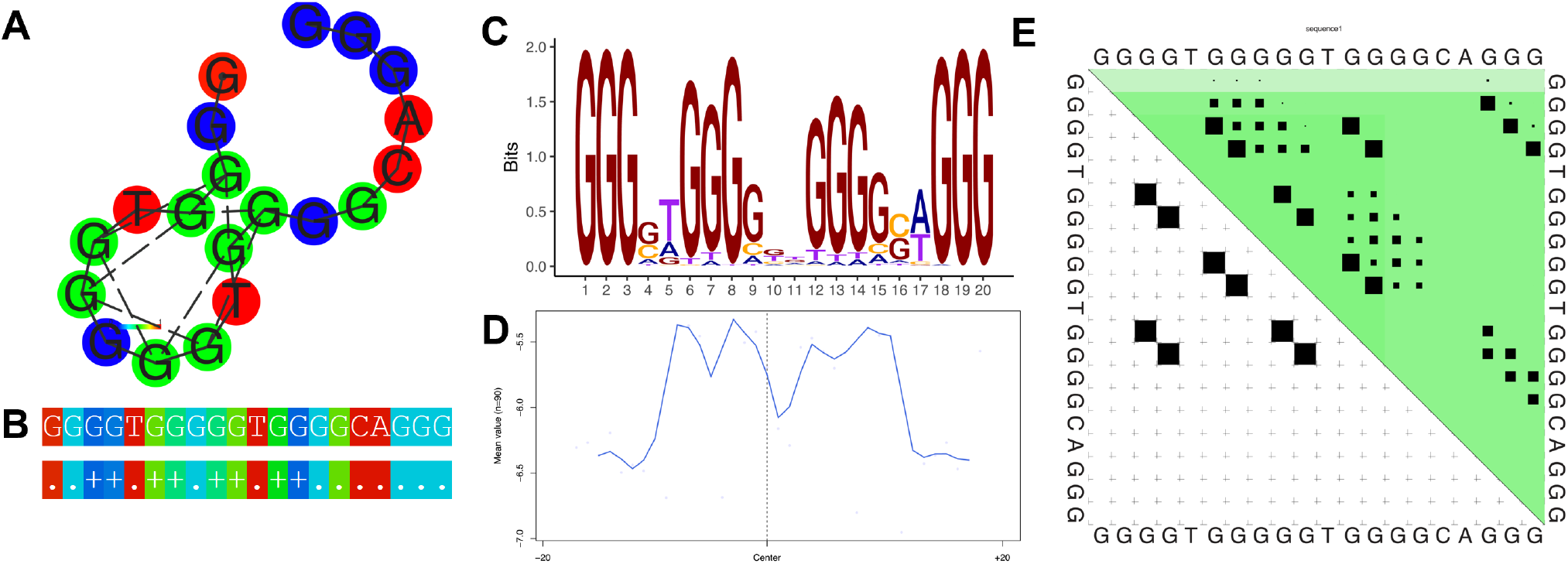
Thermodynamic properties for Family 32. (A) Centroid secondary structure with a minimum free energy of −18.0 kcal/mol using the consensus sequence of the family. (B) Dot-bracket notation showing the secondary structure. (C) Sequence logo representing the per base information content. (D) Electrostatic potential generated from all the sequences of the family using 10 flanking bases on either side of the identified G4. (E) Dot plot showing the substructures with the highest probabilities.

Family 75 is represented by 18 G4 sequences distributed over 10 chromosomes and 16 genes (Figure 5). Enriched GO:BP terms are highly related to immune differentiation and adhesion (e.g. positive regulation of T cell differentiation, positive regulation of lymphocyte differentiation, positive regulation of leukocyte cell-cell adhesion) (Supplemental Figure 3, Supplemental Table 12).

**Figure 5.**
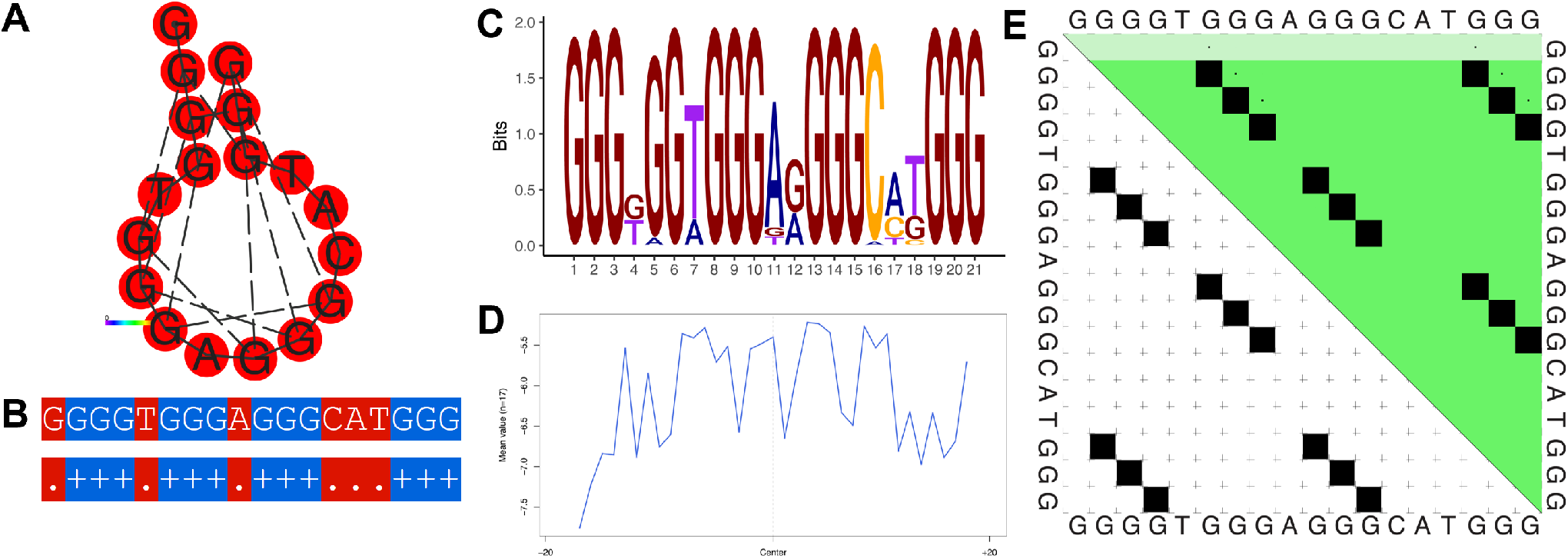
Thermodynamic properties for Family 75. (A) Centroid secondary structure with a minimum free energy of −22.82 kcal/mol using the consensus sequence of the family. (B) Dot-bracket notation showing the secondary structure. (C) Sequence logo representing the per base information content. (D) Electrostatic potential generated from all the sequences of the family using 10 flanking bases on either side of the identified G4. (E) Dot plot showing the substructures with the highest probabilities.

For Family 80, we identified 21 sequences distributed over 12 chromosomes and 21 genes (Figure 6). Genes associated with this family appear to be localized to cellular components, in particular, membranes. Enriched GO:CC categories for the genes include cytoplasmic side of membrane, plasma membrane, cytoplasmic side of plasma membrane, plasma membrane region, cell projection membrane, ficolin-1-rich granule membrane, side of membrane, cell periphery, ruffle membrane, secretory granule membrane, leading edge membrane, actin filament, extrinsic component of cytoplasmic side of plasma membrane, ruffle, membrane, extrinsic component of plasma membrane, intrinsic component of membrane, intrinsic component of endoplasmic reticulum membrane, plasma membrane protein complex, ficolin-1-rich granule, and tertiary granule (Supplemental Figure 4, Supplemental Table 13). A summary of enriched GO terms as determined from GOprofiler for selected families is present in Figure 7.

**Figure 6.**
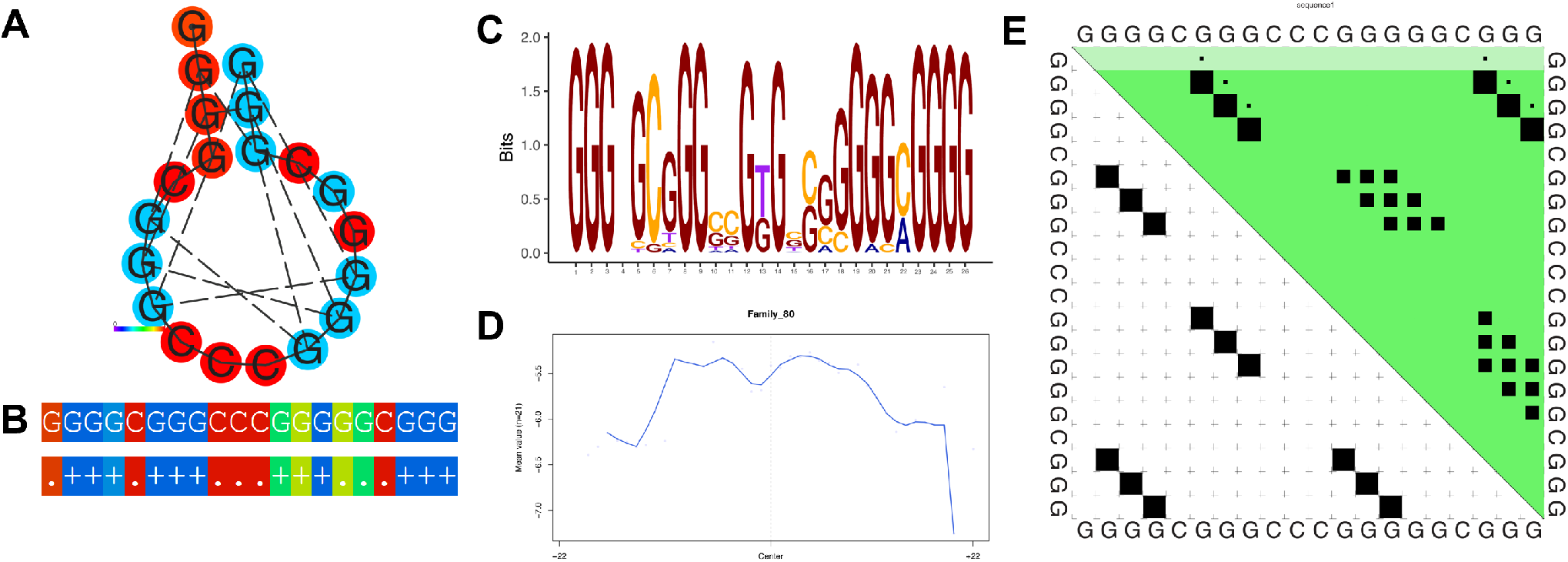
Thermodynamic properties for Family 80. (A) Centroid secondary structure with a minimum free energy of −17.38 kcal/mol using the consensus sequence of the family. (B) Dot-bracket notation showing the secondary structure. (C) Sequence logo representing the per base information content. (D) Electrostatic potential generated from all the sequences of the family using 10 flanking bases on either side of the identified G4. (E) Dot plot showing the substructures with the highest probabilities.

**Figure 7.**
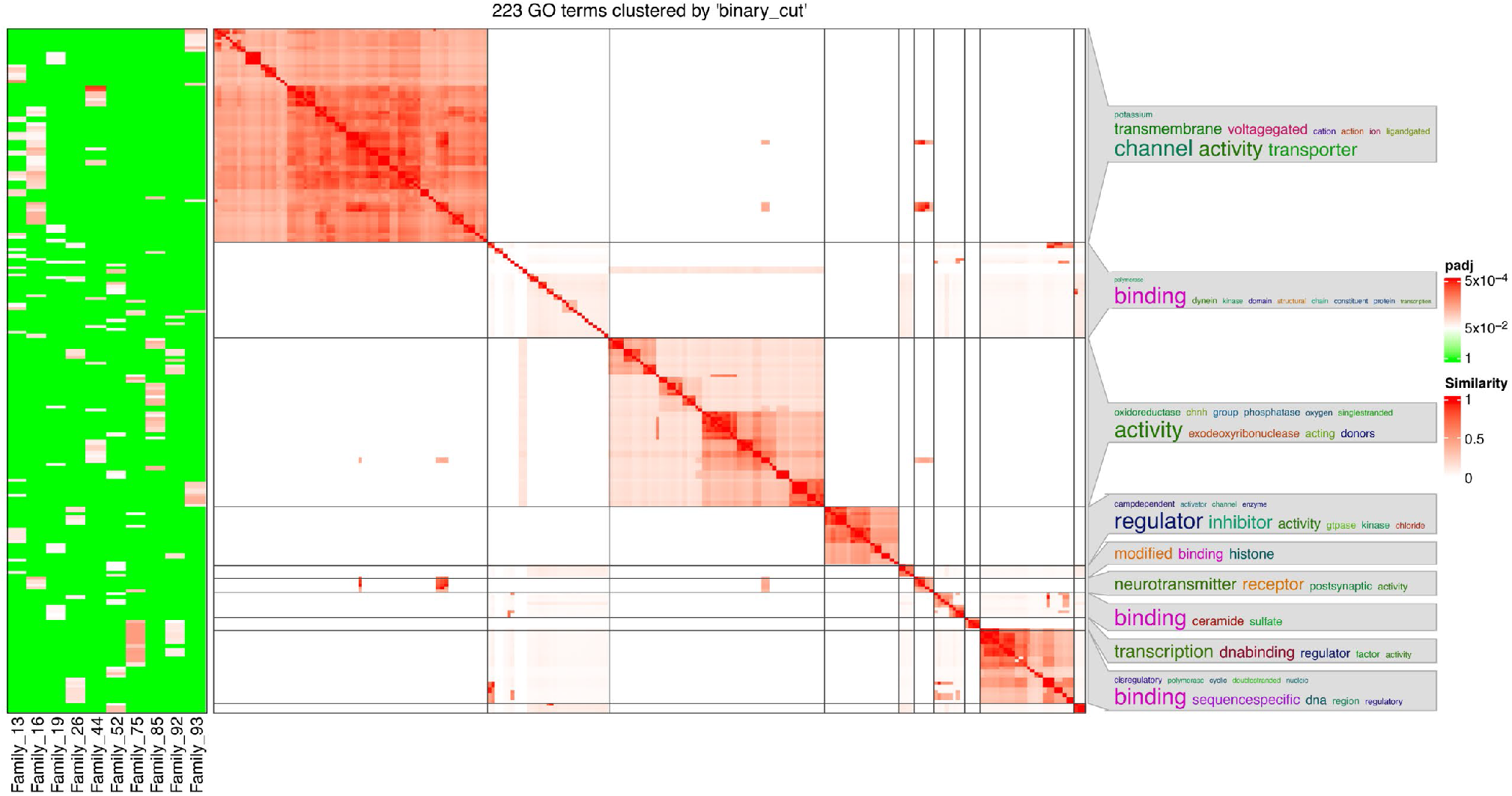
Summary of enriched GO terms for select families as determined by the GOProfiler and simplifyEnrichment R packages.

### Thermodynamic properties of select families

The free energy of the thermodynamic ensemble for the consensus sequence of Family 1 was calculated to be −28.11 kcal/mol. The frequency of the MFE structure was 50.62% with an ensemble diversity of 0, suggesting a strict conformation of tetrads for formation of a G4 structure. The minimum free energy for the family was calculated to be −27.69 kcal/mol. This family consists of six training sequences that have a single length loop with T-T-A loops (represented by 1-1-1 loops).

For Family 11, the free energy of the thermodynamic ensemble was calculated to be −20.22 kcal/mol. The frequency of the MFE structure in the ensemble is 25.23% and the ensemble diversity is 0, suggesting once again a strict conformation of tetrads for G4 formation. Family 63 is identified with the sequence G_3_AG_3_AG_3_AG_3_ and is found across 24 chromosomes and 97 genes distributed across intronic, intergenic and promoter regions. The free energy of the thermodynamic ensemble for family 63 was calculated to be −36.00 kcal/mol while the frequency of the MFE structure in the ensemble is 100% and the ensemble diversity is 0.00.

Figures 3A-D, 4A-D, 5A-D and 6A-D illustrate the thermodynamic properties for Families 4, 32, 75 and 80, respectively. Figures 3A, 4A, 5A and 6A represent the base pairing of each base in the G quadruplex sequence. Figures 3B, 4B, 5B and 6B highlight the centroid secondary structure in dot-bracket notation. A base pairing probability matrix is used to identify added information about the ensemble G4 secondary structure. Applied initially to identify different secondary structures of RNA sequences, dynamic programming provides efficient computation of base pairing probabilities for secondary structure formation. The MFE secondary structure highlighting encoding positional entropy (Figures 3C, 4C, 5C and 6C) is calculated using the consensus sequence of the G4 cluster as predicted by RNAfold. DNA shape features such as the minor groove width and electrostatic potential (Figures 3D, 4D, 5D and 6D) depend upon the charge distribution of nucleotides in a DNA sequence and affect the folding into secondary structure and transcription factor binding in these locations (77). The difference in stacking energies causing the varying hydrogen bonding patterns can be predicted in each dinucleotide step and can be used to infer minor groove width (78). The guanine amino group repeats in G quadruplexes affect charge distributions in the minor and major groove of helical DNA leading to rotation of the tetrads. We use it to annotate the different families of G quadruplex identified here. A dot plot of the structure with MFE is shown in Figures 3E, 4E, 5E, and 6E for each of the selected families.

When DNA is bent around in secondary structures such as helical or G quadruplex structures, the bend is separated based on dinucleotide sequences. Propeller twist is defined as the twist along the axis making two bases “non-coplanar” (79). Previous studies have provided evidence for the flexibility nature of the GG and GC dinucleotides with low propeller twist while AA shows the highest. The flexible nature of such a structure favors G quadruplex sequences. Low propeller twist is related with the ability for the nucleotides to slide on each other and stack in a stable manner. For each cluster, we calculated dinucleotide frequency normalized by individual length of G quadruplex, minimum free energy, minor groove width, propeller twist, helical twist, roll, and electrostatic potential with −10 and +10 region around the identified clusters of G quadruplex using DNAshapeR (80). These features address the shape, thermodynamic stability, and flexibility of rotation of the guanine amino groups, and transcription factor recognition site.

### Classification of experimentally validated G4 sequences

Using the sequences from peaks mapped from a G4 seq experiment (GEO accession GSE63874), and identified using Quadparser2, we identified all possible pG4 sequences with four tetrads and used it as query the model classifier. We classified 18,340 individual G4s identified from 22,226 distinct regions of the hg38 human genome into 95 families. Based on the clustering for experimental sequences, the major families represented are Family 73 (917 unique G4s related to 664 genes), Family 2 (25 unique G4s, 29 genes), and Family 93 (26 unique G4s, 25 genes). Family 63 has a distinct G4 sequence G_3_AG_3_AG_3_ that is repeated throughout the genome, occurring 313 times over 23 chromosomes and 204 genes.

### G4 repeat and loop length characteristics

For genes with repeats of G4 sequences (i.e. more than four tetrads), multiple G4 sequences with a variable loop length are possible (Figure 8). We identify all possible linear combinations of G tetrads for such sequences and classify all combinations of the sequences into families. This provides a way to identify multiple conformations forming G quadruplexes. One example gene with a variable length sequence is BAHCC1, a chromatin regulator known to interact with transcriptional repressors to ensure gene silencing through recognition and bind to PRC2 complex mediated H3K27me3 through chromatin compaction and histone deacetylation (81,82). Within a single G4 region, we identified repeats of 13 different sequences (length of G4 repeat: 314 bases), with each sequence being distinct enough to occur in a separate family. We also find 29 G4 sequences in NRD2, with most of the sequences occurring in Family 17, with one each also occurring in Family 7 and 10.

**Figure 8.**
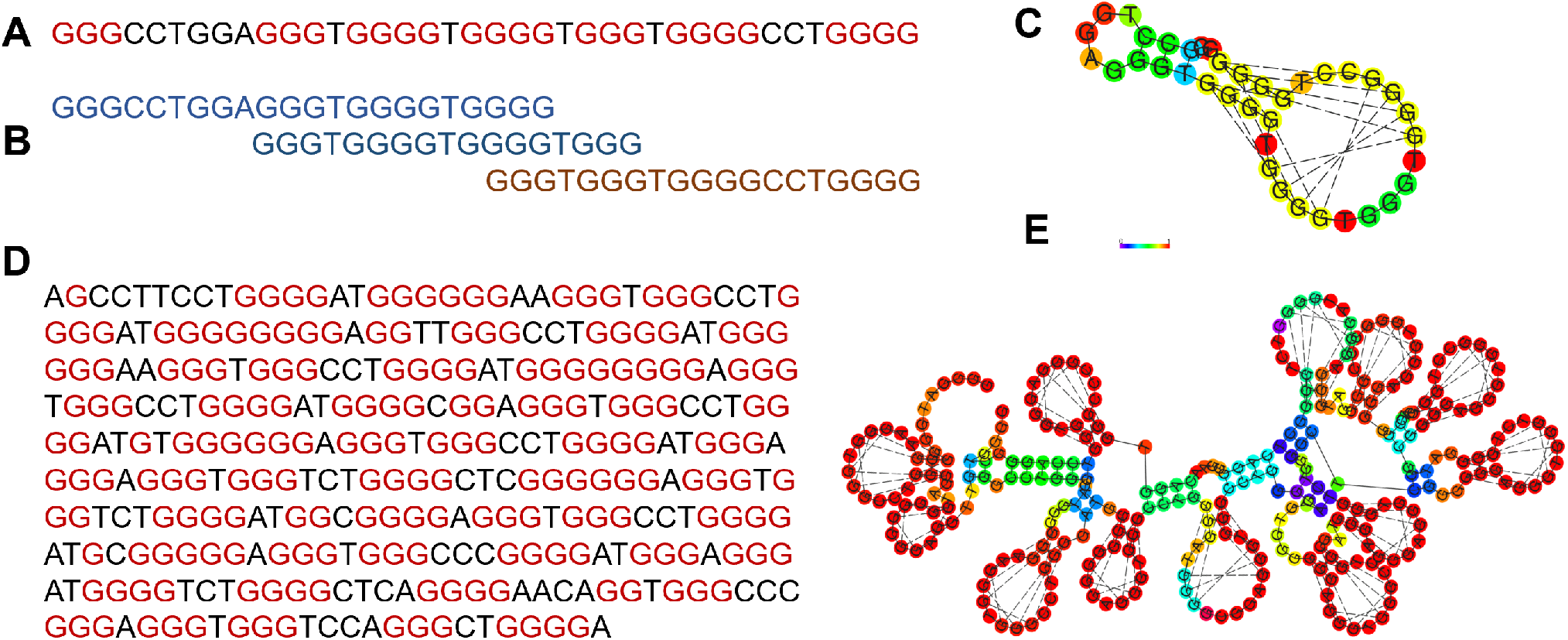
Example sequences with multiple tetrads. (A) G-quadruplex sequence from chr19:43,479,561-43,479,598 overlapping the PHLDB3 gene with guanines labelled in red. (B) Three possible alternate G4 regions for the PHLDB3 region. (C) MFE structure for the PHLDB3 region. (D) G-quadruplex sequence from chr17:81,432,609-81,432,932 overlapping the BAHCC1 gene with guanines labelled in red. (E) MFE structure for the BAHCC1 gene.

We identify similar repeats of five distinct sequences spanning an intronic region in PLOD1, which codes for lysyl hydroxylase and is involved in collagen synthesis. A 45 nucleotide G quadruplex sequence present in the promoter region of tyrosine hydroxylase (TH) can regulate transcription and has been linked with neurological and psychological disorders such as Parkinson’s and schizophrenia (83,84). We found two additional G quadruplex sequences in the opposite strand across promoter and intronic regions of TH which have matches to Family 14 and 37, respectively.

Semaphorins are a group of membrane spanning proteins that bind to Plexin (PLXNA and PLXNB) receptors to regulate axon cue signalling, cytoskeletal development and cell adhesion (85,86). The regulation and signalling of SEMA proteins with the plexin family has been a topic of study, and we identified 39 and 37 distinct G quadruplex forming sequences in the SEMA family and PLXN family respectively, with similar G4 loops present in both genes. The prediction identified multiple G4 sequences present in SEMA6C, SEMA6D, and PLXND1 with the highest match to Family 48 (Table 3). Similarly, SEMA4D, SEMA4B, and PLXNA4 shared sequences occurring in Family 17. These findings suggest that multiple regions can form G quadruplex in these genes, resulting in multiple conformations that might allow for differentiation for methylation in a pattern specific manner.

**Table 3.**
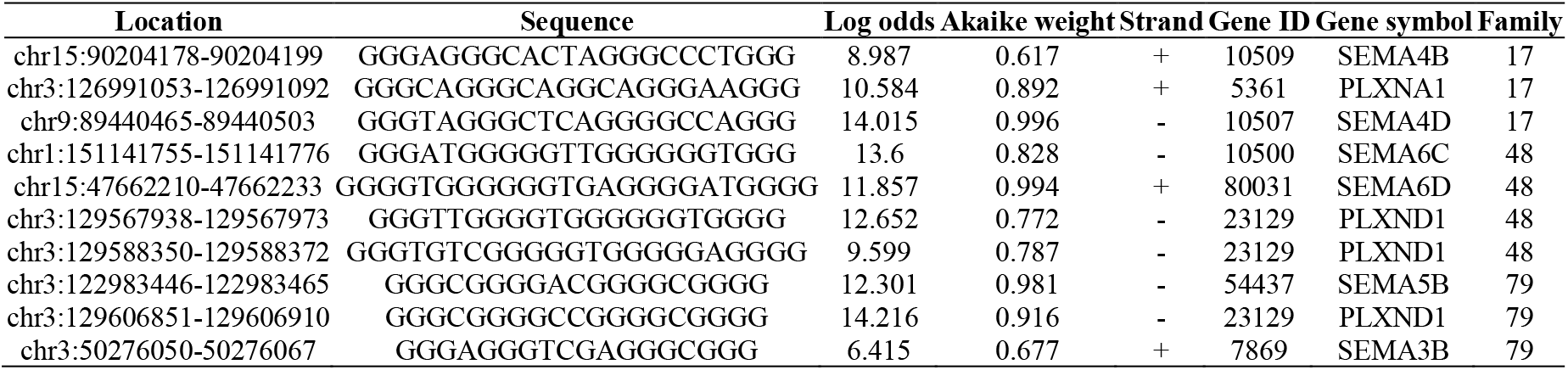
G4 sequences identified in the genic regions associated with the plexin and semaphorin gene families with high similarity to G4 Families 17, 48 and 79.

The PDB structures 22AG, 2KF8, 5LQG, and 5YEY represent telomeric quadruplex DNA forming a range of conformations with antiparallel topology based on varying physiological conditions. These telomeric G4 sequences are determined to have the highest likelihood of matching Family 22. They have a similar loop size to structure 2KM3 (27), which has a variant of CTAGGG repeat instead of TTAGGG repeats. The 2KM3 structure forms a chair type G quadruplex in K+ solution and is most similar to Family 33. Based on the sequence characteristics, these differences in structure which are caused by a one or two bp change can affect the overall prediction of the glycosidic conformation. This in turn can be used to help understand the structure based on the local environmental and interacting conditions.

A 2LXQ G4 structure is found upstream of pilin expression locus in *Neisseria gonorrhoeae*, a human pathogen 5’-G_3_TG_3_TTG_3_TG_3_ sequence is implicated in pilin antigenic variation (87). Known to form an all parallel stranded topology, the sequence was predicted to have the highest likelihood score with Family 40. A highly conserved G4 sequence at NHE III_1_ upstream of promoter 1 been studied and identified to silence transcription of c-MYC (52,88–91) and other short loop G4 sequences that form a similar topology. TAG_3_AG_3_TAG_3_AG_3_T was predicted to be most similar to Family 52 as well as Family 1. Despite following the same 1:2:1 pattern as the 2LXQ structure, the presence of adenosine in place of thymidine as the linker loops is considered as a different family.

Experimental evidence shows that G4s with short loop sequences favor a parallel topology while structures with longer loops tend to form hybrid or antiparallel structures (92). Sequences with thymine compared to adenine as a single length loop have been found to have higher melting point than a single A base (93). Given our clustering scheme, multiple sequences with short loops can show high log-odds for multiple families. In these cases, the Akaike weight can help guide the context and identify multiple families similar to such sequences.

### G4 in enhancers

Potential regulatory roles of G4 families were analyzed by looking at the overlap between G4s and enhancers. The overlapping enhancers were then used as input into the Gene-Enhancer link correlation (http://compbio.mit.edu/epimap/) to determine if any of the overlapping enhancers were correlated with gene expression, and if so, in what cell type. We then performed hierarchical clustering of the intersecting G4s based on the correlations. Two main groups of interest result.

In the first group, 102 G4 sequences are found in 158 genes, belonging to 57 distinct G4 families. GO:BP analysis of this group results in terms associated with immune system processes (e.g. T cell receptor signalling pathway, regulation of leukocyte proliferation, interleukin-10 production and regulation of cytokine production involved in immune response) or signalling cascades (e.g. positive regulation of ERK1 and ERK2 cascade, calcium ion transmembrane import into cytosol, and Fc receptor signalling pathway) (Supplemental Figure 5, Supplemental Table 14).

The second group had ubiquitous high correlation with all cell types in the dataset (Supplemental Figure 7). We identified 234 genes in this group with 107 distinct G4s belonging to 55 distinct families and found enrichment of terms relating to immune responses (e.g. defense response to virus, cytokine-mediated signalling pathway and regulation of defense response), regulated cell death (e.g. apoptotic signalling pathway, extrinsic apoptotic signalling pathway via death domain receptors, and positive regulation of programmed cell death), lipid biosynthesis (e.g. regulation of lipid biosynthetic process and response to fatty acid), and migration (e.g. positive regulation of protein localization and positive regulation of mononuclear cell migration) (Supplemental Figure 6, Supplemental Table 15).

Based on the enriched terms from two groups, it appears as though the G quadruplex functions across multiple pathways in different cell types. It is possible that tissue specific conditions control the actual G4 formation at specific sites, leading to tissue specific functional regulation. The results of the enhancer-gene correlation related to the presence of G4 sequences in enhancer regions are more likely to affect genes in thymus, T cell and lymphoblastoid cells.

## DISCUSSION

Our clustering methodology presented here has allowed for the construction of families of G quadruplexes based on sequence similarity, loop length and composition, and thermodynamic properties. Further analysis of these families uncovers that many of these families have functional enrichments, indicating they are potentially regulated by common mechanisms since they have structural similarities. Comparing our results to the only previously studied family, Pu27, shows a high agreement, with 12 of the 18 Pu27 members belonging to Family 1 (Table 4).

**Table 4.**
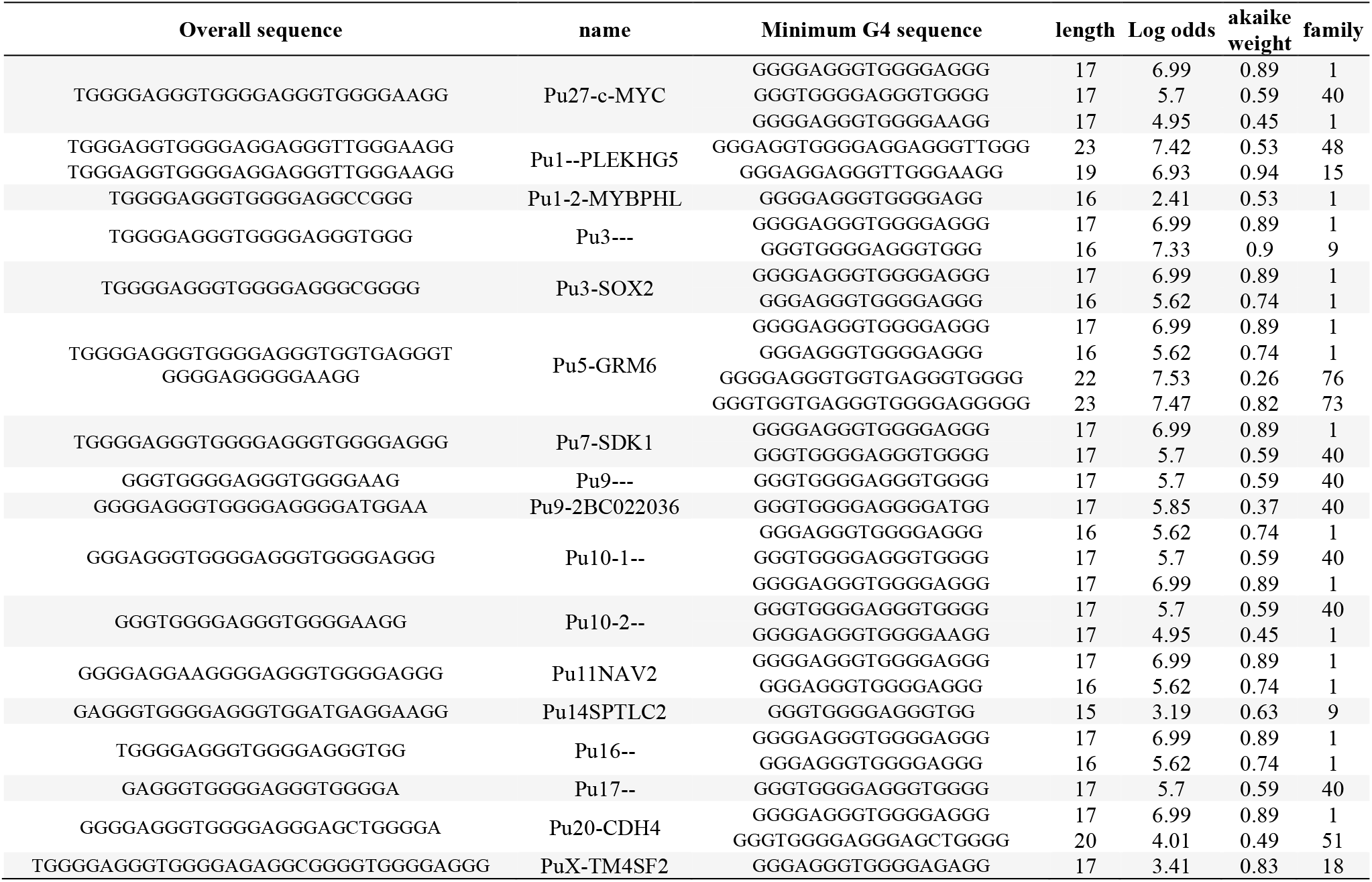
Family prediction for previously identified Pu27 family of G4 sequences.

Multiple transcription factors can bind to the alternative motifs present in G quadruplex regions (57) in response to environmental conditions in response to stimuli. These conditions trigger the folding and unfolding of G4 structures. We identify Family 40 as an alternate conformation in these sequences as multiple tetrads allow the alternate guanine bonds for stable structure. Nucleoside diphosphate kinase (NM23-H2) (94,95) has been previously identified to unfold Pu27 causing the increase of c-MYC transcription while nucleolin (96) has been identified to stabilize the G4 structure. The mechanism of TF binding and control of expression of the expression of c-MYC gene is poorly understood and is beyond the scope of prediction through this model. However, this process sheds light upon the collection of multiple conformation of structures in equilibrium which can alter the change in binding grooves for transcription factors and further downstream process. Failing to take the dynamic nature of Pu27 and other G quadruplex sequences in the genome into account could limit the effectiveness of any therapeutic compounds designed to target it.

Several G4 ligands are currently being considered for their therapeutic value. For instance, CX-5461 is utilized for treatment of BRCA1/2 deficient tumors through topoisomerase II inhibition (97,98), Melanoma cell lines have been treated with G4 ligand RHPS4 that targets the MYC gene (99) among others. G4 ligands such as APTO-253 (100), TMPyP4 (101), telomestatin (102) have been tested for their effect on leukemia. Despite showing promising results and inhibition of cell growth, telomerase shortening and senescence was observed with some of the G4 ligands in different leukemia cells (103). With the information of G4 formation and binding of specific ligands to multiple G4 structures, identification of G4 clusters can provide additional information about DNA damage occurring or novel binding motifs of specific G4 ligands.

G4 structures contribute to genomic instability and the proliferative nature of different cancers. The context and location of individual G4 can serve as a roadblock for many oncogenes, but the presence of G4 in the vicinity of a tumor suppressor gene can have the opposite effect. To understand the intended consequence of these targets for all the G4 ligands, it is important to characterize the thousands of G4 structure present in the genome and classify these structures based on their structure, function, or localization.

This study identifies related families of G quadruplex sequences within the human genome and presents them as clusters described by both an MSA and HMM. The approach described here can easily be applied to other model organisms where G4s are known to play regulatory roles. Many of these clusters were functionally annotated, allowing for a more complete understanding of these structures as well as identification of multiple targets for testing of G4 ligands. As more information on experimentally validated G4 regions becomes available, refinement of clustering methodologies will yield more informative G4 families.

## AVAILABILITY

All code and resulting data is available in the GitHub repository (https://github.com/UofLBioinformatics/G4-Cluster)

## ACCESSION NUMBERS

Experimentally validated G4 sequences were obtained from a publicly available dataset which is available in GEO under accession number GSE63874.

## SUPPLEMENTARY DATA

Supplementary Data are available online.

## ACKNOWLEDGEMENT

We wish to thank members of the Kentucky IDeA Networks of Biomedical Research Excellence (KY INBRE) Bioinformatics Core and the Rouchka and Park labs for their valuable feedback.

## FUNDING

This work was supported by the National Institutes of Health [P20GM103436]. The contents of this work are solely the responsibility of the authors and does not reflect the official views of the National Institutes of Health. Funding for open access charge: National Institutes of Health, P20GM103436.

## CONFLICT OF INTEREST

The authors declare no conflicts of interest.

## Notes

https://github.com/UofLBioinformatics/G4-Cluster

### Competing Interest Statement

The authors have declared no competing interest.

