## Supplemental Figures for "Structural and functional classification of G-quadruplex families within the human genome"

### SUPPLEMENTAL FIGURE 1

#### Supplemental Figure 1. Top 25 GO:BP enrichments for Family 4.

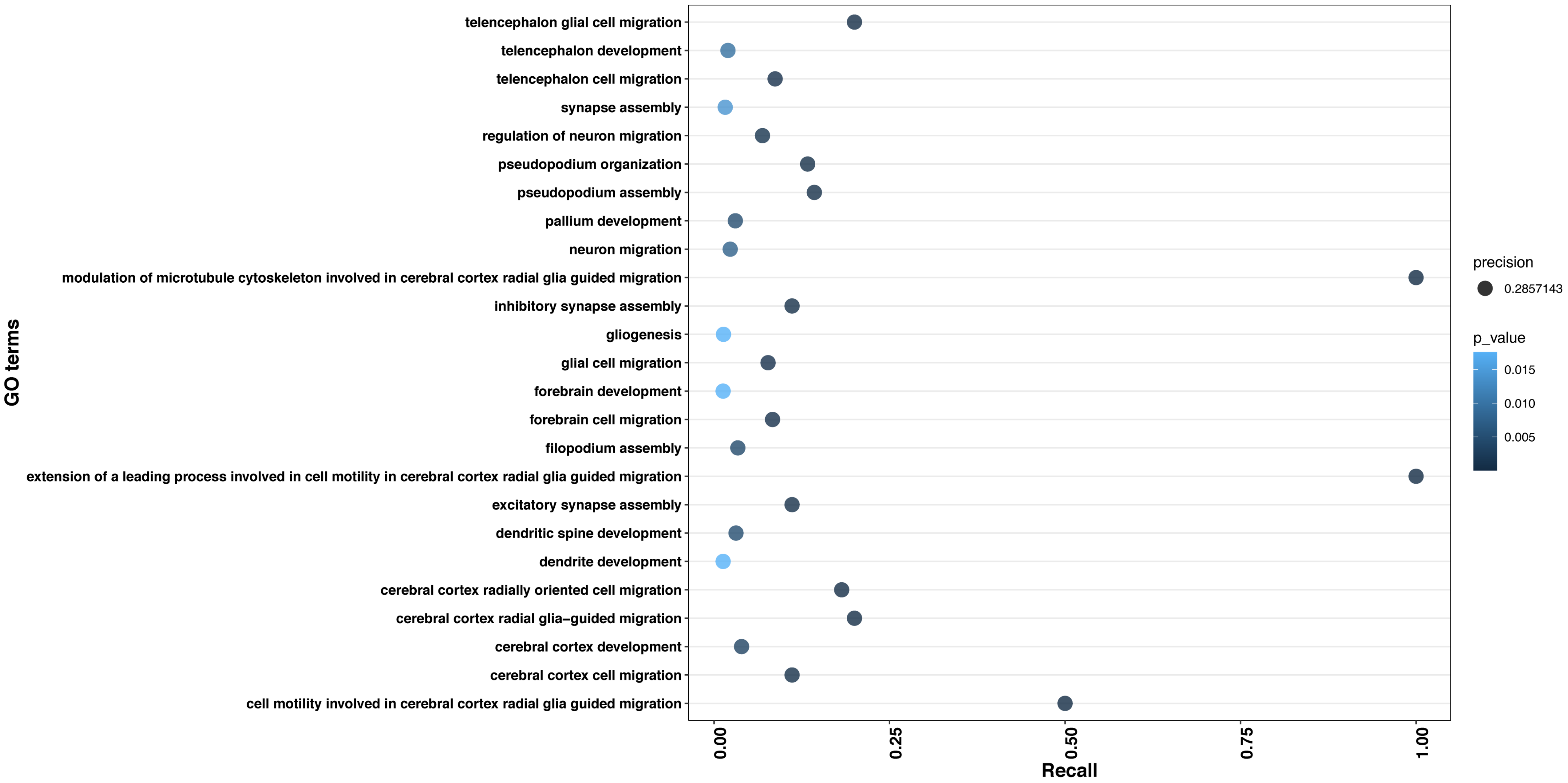

### SUPPLEMENTAL FIGURE 2

#### Supplemental Figure 2. Top 25 GO:BP enrichments for Family 32.

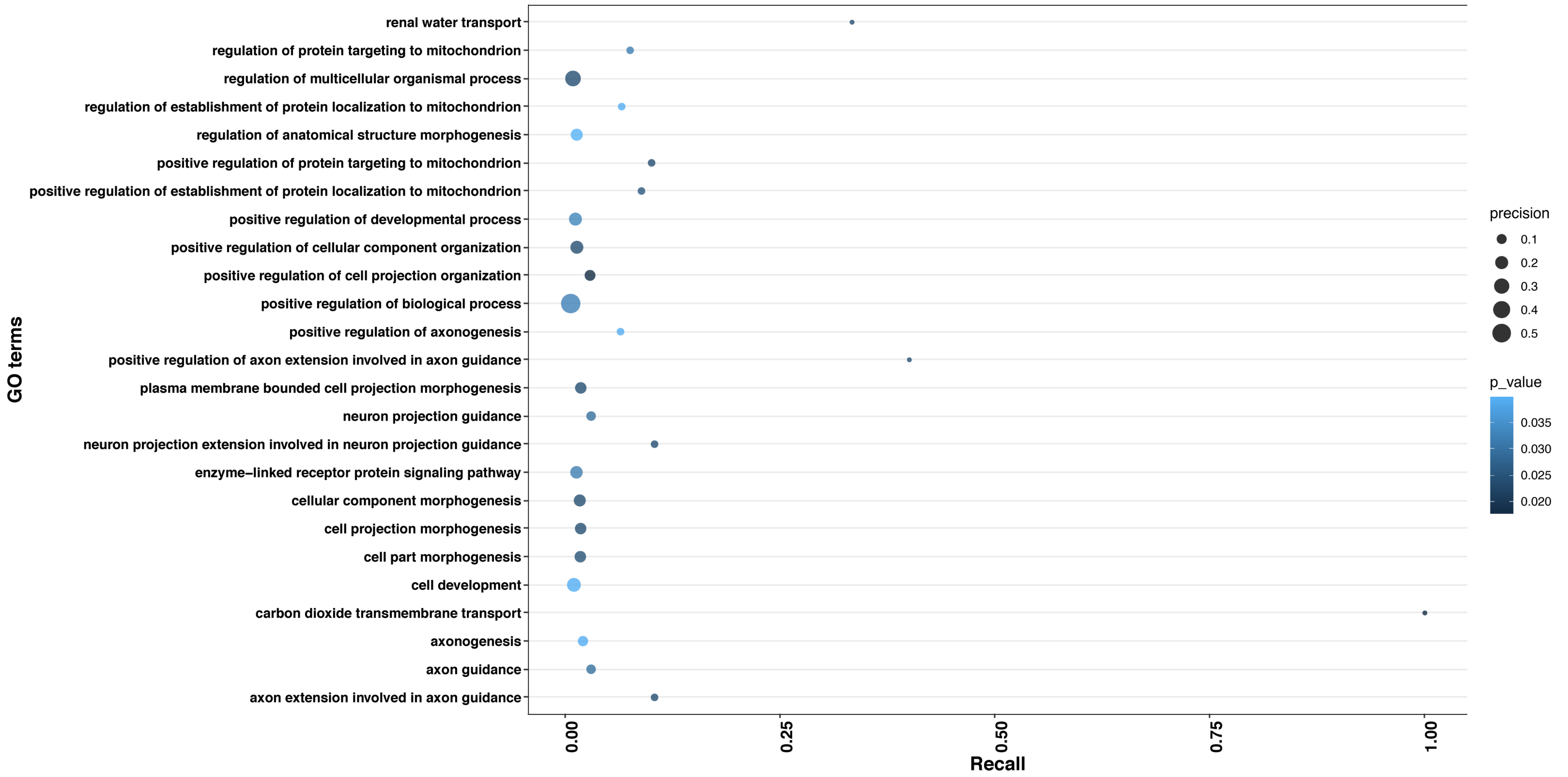

### SUPPLEMENTAL FIGURE 3

**Supplemental Figure 3. Top 25 GO:BP enrichments for Family 74.**

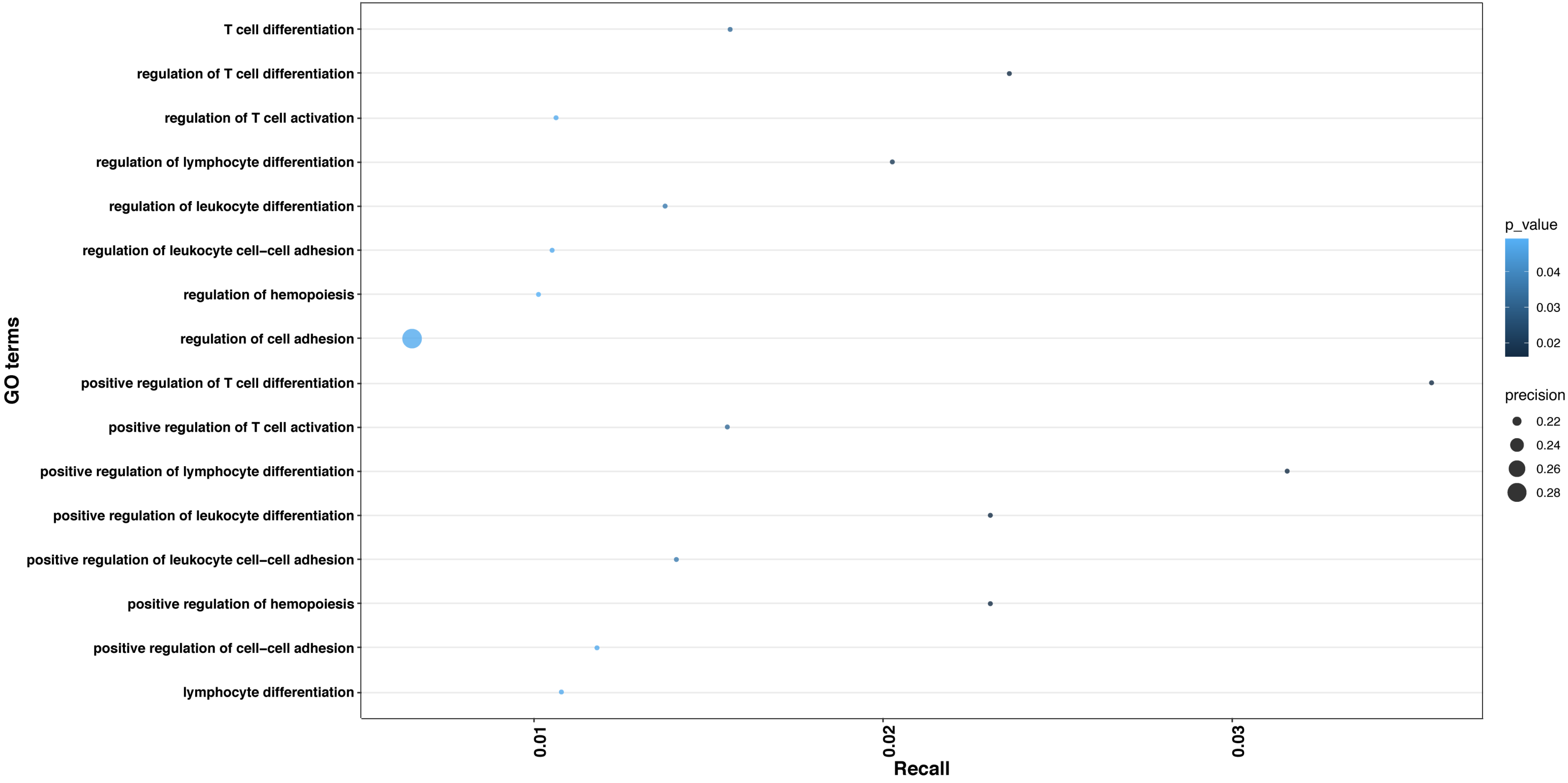

### SUPPLEMENTAL FIGURE 4

Supplemental Figure 4. Top 25 GO:BP enrichments for Family 80.

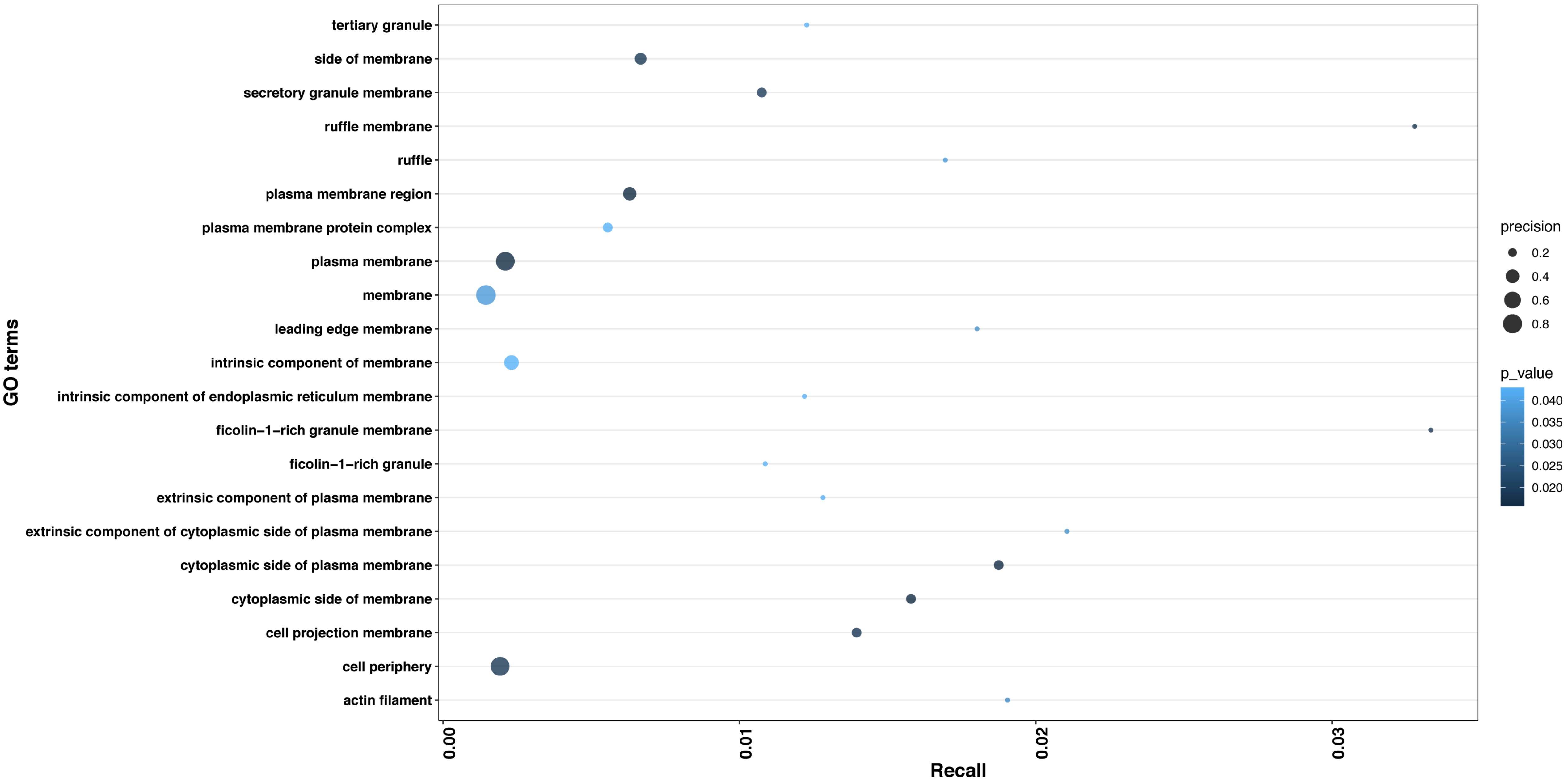

### SUPPLEMENTAL FIGURE 5

**Supplemental Figure 5. Top 25 GO:BP enrichments for experimentally validated G4s overlapping enhancers, group 1.**

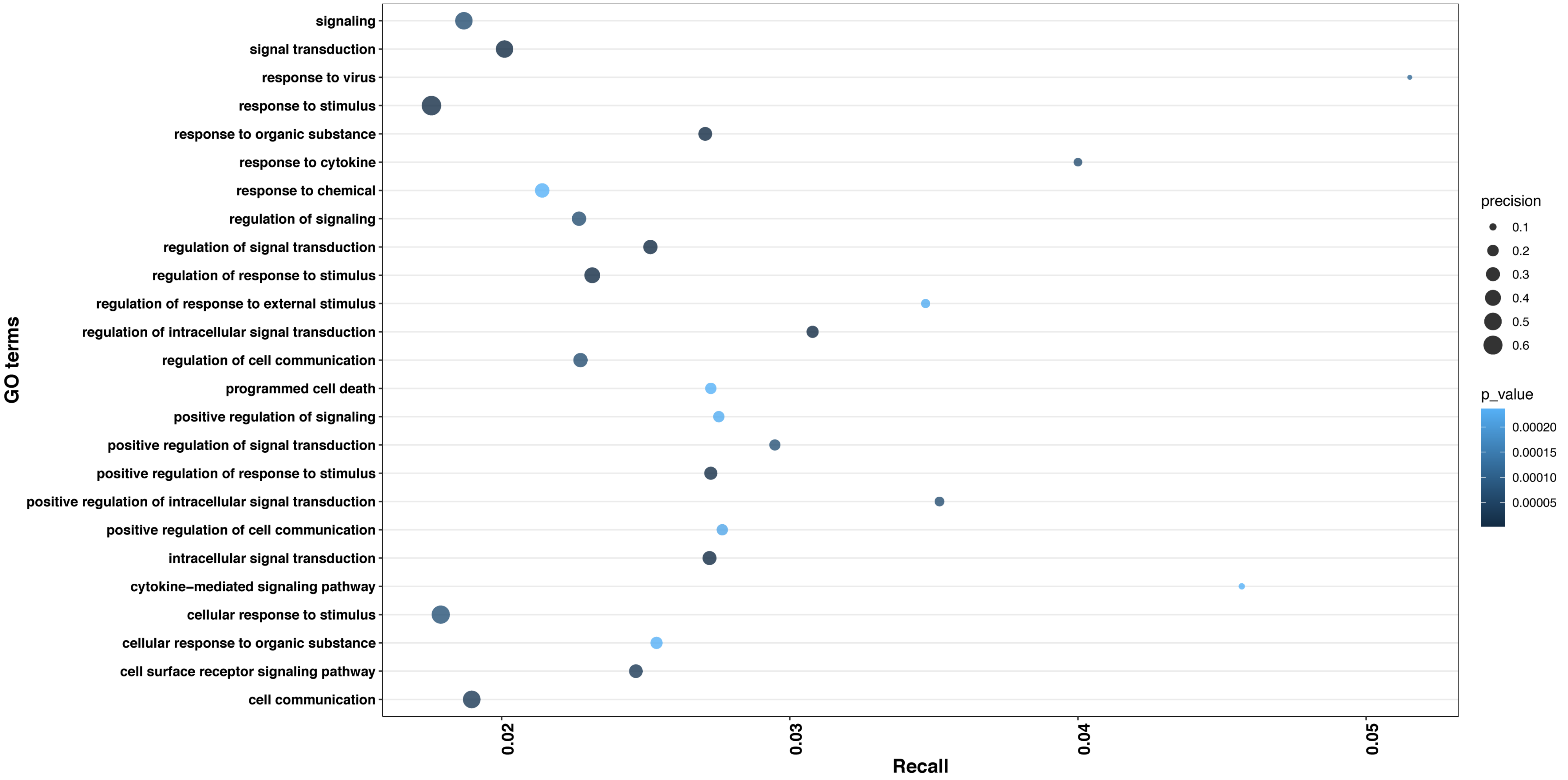

### SUPPLEMENTAL FIGURE 6

**Supplemental Figure 6. Top 25 GO:BP enrichments for experimentally validated G4s overlapping enhancers, group 2.**

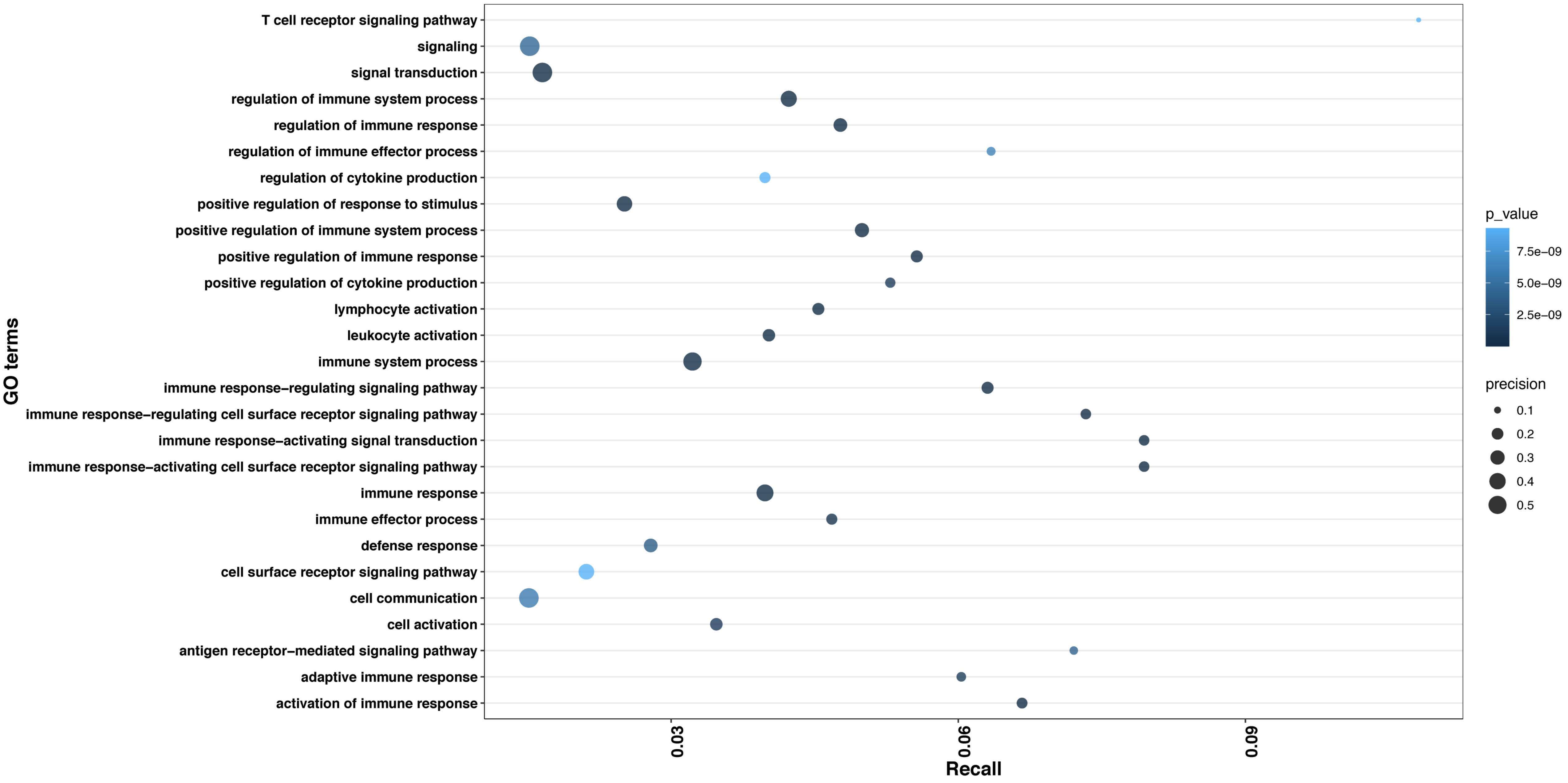

### SUPPLEMENTAL FIGURE 7

**Supplemental Figure 7. Correlation of selected enhancers consisting of pG4 with gene expression in multiple cell types utilizing the epimap correlation group-link data.**

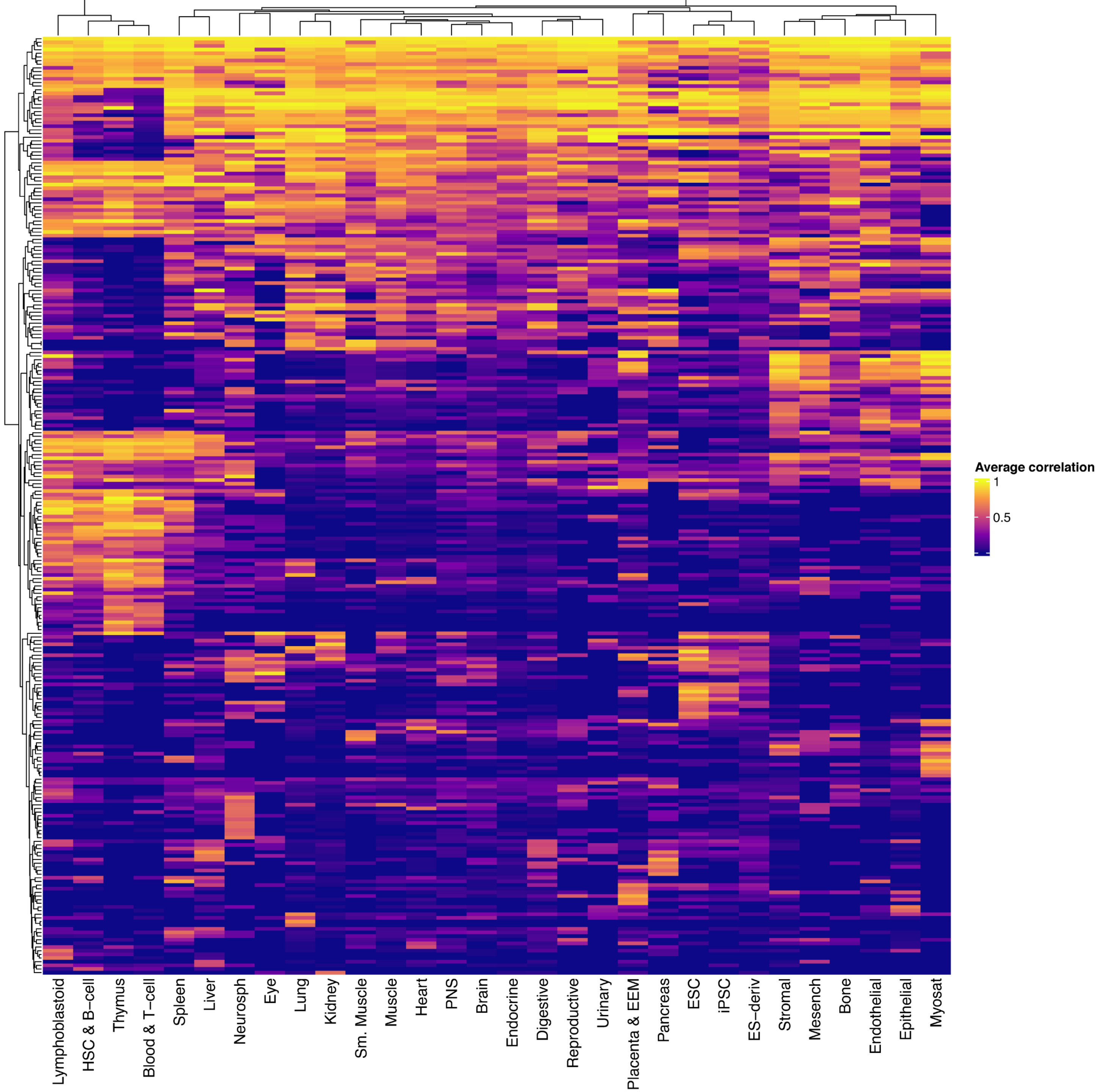
