## Supplemental Tables for "Structural and functional classification of G-quadruplex families within the human genome"

**Supplemental Table 1. Summary of Family 23.**

| Sequence | Location | Experimental Evidence | Gene ID | Anntoation |
|---|---|---|---|---|
| GGGTGGCGGGTGGGGGAGGG | chr10:123316717-123316737 | absent | 9184 | Intergenic |
| GGGTGAGGGTGCGGGTGAGGG | chr10:124610344-124610364 | present | 64077 | Promoter |
| GGGTGAGGGTGCGGGTGAGGG | chr10:124610377-124610397 | present | 64077 | Promoter |
| GGGTGAGGGTGCGGGTGAGGG | chr10:124610509-124610529 | present | 64077 | Promoter |
| GGGTGAGGGTGCGGGTGAGGG | chr10:124610542-124610562 | present | 64077 | Promoter |
| GGGTGAGGGTGCGGGTGAGGG | chr10:124610647-124610667 | present | 64077 | Promoter |
| GGGTGAGGGTGCGGGTGAGGG | chr10:124610752-124610772 | absent | 64077 | Promoter |
| GGGTGAGGGTGCGGGTGAGGG | chr10:124610785-124610805 | absent | 64077 | Promoter |
| GGGTGAGGGTGCGGGTGAGGG | chr10:124610956-124610976 | absent | 64077 | Promoter |
| GGGTGAGGGTGAGGGTGAGGG | chr10:124610410-124610430 | present | 64077 | Promoter |
| GGGTGAGGGTGAGGGTGAGGG | chr10:124610857-124610877 | absent | 64077 | Promoter |
| GGGTGAGGGTGAGGGTGAGGG | chr10:124610890-124610910 | absent | 64077 | Promoter |
| GGGTGGAGGGGTGGGGTGGGG | chr10:132548209-132548229 | present | 3632 | Intron |
| GGGTGGGGGGTGGGGAGAGGG | chr10:15725418-15725438 | present | 8516 | Intergenic |
| GGGTGGGGGGTGGGGAGGGG | chr10:21717394-21717413 | present | 8028 | Intron |
| GGGTGGGGGGTGGGGAGGGG | chr18:11976921-11976940 | absent | 3613 | Promoter |
| GGGTTGGGGGTAGGGTGGGGG | chr10:62544042-62544062 | present | 22891 | Intron |
| GGGGTGTGGGGCAGGGATGGGG | chr10:70678644-70678665 | present | 140766 | Intron |
| GGGTGGGGGCTGGGGAGAGGG | chr10:78293711-78293731 | absent | 414243 | Intron |
| GGGTCGGGGGCGGGGGAGGG | chr11:101129211-101129230 | present | 101054525 | Promoter |
| GGGTGGGAGTGGGATGAGGG | chr11:125159894-125159913 | present | 103695364 | Promoter |
| GGGTTGGGGAGTGGGGTTGGG | chr11:43926065-43926085 | present | 100507300 | Intron |
| GGGTTGGGGGTGGGGTGGGG | chr11:44101102-44101121 | present | 2132 | Promoter |
| GGGATGTGGGAAGGGATGGGG | chr11:69269215-69269235 | present | 26579 | Intergenic |
| GGGGTGGGTGTGGGGTGGGG | chr12:113876018-113876037 | present | 9904 | Intron |
| GGGGTGGGTGTGGGGTGGGG | chr16:46909101-46909120 | present | 84706 | Intron |
| GGGATGGGGGTCGGGTGGGG | chr12:131917175-131917194 | present | 8408 | Promoter |
| GGGGTGGGGGTGGGAGAGGG | chr12:2537890-2537909 | present | 775 | Intron |
| GGGGTGAGGGTAGGGATGGGG | chr12:6640013-6640033 | absent | 84519 | Promoter |
| GGGGTTGGGGAAGGGAGGGGG | chr13:112058816-112058836 | present | 6656 | Intergenic |
| GGGGTGGGGAAGGGATTGGGG | chr13:53043320-53043340 | absent | 10562 | Intron |
| GGGTGGGGGTGGGGGGCAGGG | chr14:100443444-100443464 | present | 79446 | Intron |
| GGGTGGGGGTGGGGCAAGGG | chr14:103532880-103532899 | present | 115708 | Promoter |
| GGGGTGGGTGAAGGGATGGGGG | chr14:105596969-105596990 | present | 102465871 | Intergenic |
| GGGGTGGGTGAAGGGATGGGGG | chr14:105597009-105597030 | present | 102465871 | Intergenic |
| GGGGTGGGTGAAGGGATGGGGG | chr14:105597050-105597071 | present | 102465871 | Intergenic |
| GGGGTGGGTGAAGGGATGGGGG | chr14:105597091-105597112 | present | 102465871 | Intergenic |
| GGGGTGGGTGAAGGGATGGGGG | chr14:105597132-105597153 | absent | 102465871 | Intergenic |
| GGGGTGGGTGAAGGGATGGGGG | chr14:105717880-105717901 | present | 102465871 | Intergenic |
| GGGGTGGGTGAAGGGATGGGGG | chr14:105717921-105717942 | present | 102465871 | Intergenic |
| GGGGTGGGTGAAGGGATGGGGG | chr14:105717962-105717983 | present | 102465871 | Intergenic |
| GGGGTGGGTGAAGGGATGGGGG | chr14:105718002-105718023 | present | 102465871 | Intergenic |
| GGGGTGGGTGAAGGGATGGGGG | chr14:105718043-105718064 | absent | 102465871 | Intergenic |
| GGGGTGGGTGAAGGGATGGGGG | chr14:105718084-105718105 | present | 102465871 | Intergenic |
| GGGGTGGGTGAAGGGATGGGGG | chr14:105718125-105718146 | present | 102465871 | Intergenic |
| GGGGTGGGTGAAGGGATGGGGG | chr14:105718125-105718146 | present | 102465871 | Intergenic |
| GGGGTGGGTGAAGGGATGGGGG | chr14:105718166-105718187 | present | 102465871 | Intergenic |
| GGGGTGGGTGAAGGGATGGGGG | chr14:105718206-105718227 | present | 102465871 | Intergenic |
| GGGGTGGGTGAAGGGATGGGGG | chr14:105718247-105718268 | present | 102465871 | Intergenic |
| GGGGTGGGTGAAGGGATGGGGG | chr14:105718288-105718309 | absent | 102465871 | Intergenic |
| GGGGTGGGTGAAGGGATGGGGG | chr14:105718329-105718350 | absent | 102465871 | Intergenic |
| GGGGTGGGTGAAGGGATGGGGG | chr14:105718370-105718391 | absent | 102465871 | Intergenic |
| GGGGTGGGTGAAGGGATGGGGG | chr14:105718411-105718432 | absent | 102465871 | Intergenic |
| GGGGTGGGTGAAGGGATGGGGG | chr14:105718452-105718473 | absent | 102465871 | Intergenic |
| GGGGTGGGTGAAGGGATGGGGG | chr14:105718493-105718514 | absent | 102465871 | Intergenic |
| GGGGTGGGTGAAGGGATGGGGG | chr14:105718534-105718555 | absent | 102465871 | Intergenic |
| GGGGTGGGTGAAGGGATGGGGG | chr14:105718575-105718596 | absent | 102465871 | Intergenic |
| GGGGTGGGTGAAGGGATGGGGG | chr14:105718616-105718637 | absent | 102465871 | Intergenic |
| GGGGTGGGTGAAGGGATGGGGG | chr14:105718657-105718678 | absent | 102465871 | Intergenic |
| GGGGTGGGTGAAGGGATGGGGG | chr14:105718698-105718719 | absent | 102465871 | Intergenic |
| GGGGTGGGTGAAGGGATGGGGG | chr14:105718739-105718760 | absent | 102465871 | Intergenic |
| GGGGTGGGTGAAGGGATGGGGG | chr14:105718780-105718801 | present | 102465871 | Intergenic |
| GGGGTGGGTGAAGGGATGGGGG | chr14:105718821-105718842 | present | 102465871 | Intergenic |
| GGGGTGGGTGAAGGGATGGGGG | chr14:105718862-105718883 | present | 102465871 | Intergenic |
| GGGGTGGGGGTGGGGTGGGGG | chr14:19037660-19037680 | present | 100508046 | Intron |
| GGGGTGGGGGTGGGGTGGGGG | chr22:15760289-15760309 | present | 106146148 | Intron |

**Supplemental Table 1. (Continued).**

| Sequence | Location | Experimental Evidence | Gene ID | Anntoation |
|---|---|---|---|---|
| GGGTCAGGGGTGGGGTGGGG | chr14:50864482-50864501 | present | 145447 | Intron |
| GGGTTGGGGGCGGGGGTGGG | chr14:65413539-65413558 | present | 2530 | Promoter |
| GGGGTGGGGGTGGGGCGGGG | chr14:68908184-68908203 | present | 87 | Intron |
| GGGGTGAGGGGAAGGGATGGG | chr14:72421211-72421231 | absent | 9628 | Promoter |
| GGGTTGGGGGGGTGGGTGGGG | chr15:65356691-65356711 | present | 9543 | Promoter |
| GGGTGGGGCTGGGGTGTGGG | chr15:88391894-88391913 | absent | 26589 | Intergenic |
| GGGTTTGGGGTGGGGGTAGGG | chr15:89109459-89109479 | absent | 11057 | Intron |
| GGGATTGGGGGTGGGGGAGGG | chr16:27805003-27805023 | present | 23247 | Intron |
| GGGGTGGGGGCCGGGATGGGG | chr16:51096031-51096051 | present | 6299 | Intergenic |
| GGGTGGGGGTGGGCTGAGGG | chr16:67172198-67172217 | present | 8996 | Promoter |
| GGGAGGGGGGTGGGGGGAGGG | chr16:7709746-7709765 | present | 54715 | 3' UTR |
| GGGGTGGGGGCAGGGTGGGG | chr16:79383220-79383239 | present | 51741 | Intergenic |
| GGGTGGGGGTGGGAGTCAGGG | chr16:86705437-86705457 | present | 101928614 | Intergenic |
| GGGAGTGGGGGTGGGGTGGGG | chr17:12982890-12982910 | present | 60528 | Exon |
| GGGAGTGGGGGTGGGGTGGGG | chr9:76754262-76754282 | present | 50652 | Intron |
| GGGTTAGGGGTGGGGTGGGGG | chr17:18127574-18127594 | present | 51168 | Intron |
| GGGGTTGGGGGAGGGAGGGG | chr17:1880069-1880088 | present | 6117 | Promoter |
| GGGCTGGGGGTGGGGAAGGG | chr17:50867047-50867066 | present | 400604 | Promoter |
| GGGGTGGGGAGTGGGGTGGGG | chr17:65457424-65457444 | present | 105827617 | Promoter |
| GGGGTGGGGGGAAGGGAGGGGG | chr17:82020660-82020681 | present | 201254 | Promoter |
| GGGTTGGGGGAGGGATTGGG | chr18:24852934-24852953 | absent | 105372028 | Intron |
| GGGGGTGGGGGTGGGGTGGGG | chr18:26503844-26503864 | present | 284252 | Intron |
| GGGGTTGGGGGTGGGGGGGG | chr18:62610383-62610402 | present | 54877 | Intergenic |
| GGGGTGGCGGGTGGGGTGGGG | chr18:77413747-77413767 | present | 2587 | Intergenic |
| GGGTTGCGGGGTAGGGGAGGG | chr19:35851196-35851216 | absent | 4868 | Promoter |
| GGGGTGGGGGAAGGGAGGGG | chr19:408850-408869 | present | 126567 | Promoter |
| GGGTGGGGGTGGAGGGAGGG | chr19:5117318-5117337 | present | 23030 | Promoter |
| GGGTTGGGGGTGGGCTGGGG | chr1:10670083-10670102 | present | 54897 | Intron |
| GGGTGGGGCTGGGAGTGAGGG | chr1:1075343-1075363 | present | 401934 | Promoter |
| GGGTTGGGACTGGGGTGGGG | chr1:157198319-157198338 | present | 2117 | Intergenic |
| GGGGTGGGGATGGGATGGGG | chr1:202304270-202304289 | present | 59352 | Intron |
| GGGGTGGGGGAAGGGGGTGGGG | chr1:211260741-211260762 | present | 55758 | Promoter |
| GGGTGGAGGGTGGGGTGGGG | chr1:21865234-21865253 | present | 3339 | Intron |
| GGGTGGAGGGTGGGGTGGGG | chr1:30271549-30271568 | absent | 101929406 | Intergenic |
| GGGTGGGGATGGGAGTGAGGG | chr1:23911503-23911523 | present | 1269 | Promoter |
| GGGGTGGGCGGTGGGGTGGGG | chr1:24584671-24584691 | present | 400746 | Promoter |
| GGGTTGGGGGTGGGGTGGGGGG | chr1:36566733-36566754 | present | 1441 | Intergenic |
| GGGTGGGGGTGGGGTGAGGG | chr1:36576371-36576390 | present | 1441 | Intergenic |
| GGGCAGGGGTGGGGTGAGGG | chr1:44506457-44506476 | present | 100847089 | Intron |
| GGGGTGGGCGAAGGGAGTGGGG | chr1:53947258-53947279 | present | 115353 | Promoter |
| GGGGGGGAGTGGGGTGAGGG | chr1:87522994-87523013 | absent | 100505768 | Intergenic |
| GGGGCTGGGGGTGGGGGAGGG | chr20:2820918-2820938 | present | 100288797 | Promoter |
| GGGTTGCGGGGTGGGGTGGGGG | chr20:29750401-29750422 | absent | 245929 | Intergenic |
| GGGTTGCGGGGTGGGGTGGGGG | chr20:30497208-30497229 | absent | 245929 | Intergenic |
| GGGTTGGGGGAGGGGGTGGG | chr20:32980815-32980834 | present | 140732 | Downstream |
| GGGGTGGAGGGTGGGGTGAGGG | chr20:34705516-34705537 | present | 58476 | Promoter |
| GGGTGGGGATGGGGGGAGGG | chr20:43934072-43934091 | present | 84969 | Intron |
| GGGTTGGGGGGTGGGGTGGG | chr20:46247013-46247032 | present | 64405 | Promoter |
| GGGTTGGGGGTAGGGGGTGGG | chr21:32537338-32537358 | present | 59271 | Intergenic |
| GGGTTGCGGGGTGGGGTGGG | chr22:11262057-11262076 | absent | 81061 | Intergenic |
| GGGACGGGGGTGGGGTGGGG | chr22:18855039-18855058 | present | 8214 | Intergenic |
| GGGGTGGGGTTGGGGTGGGG | chr22:19999303-19999322 | present | 421 | Intron |
| GGGGAGGGGGGAGGGATGGGG | chr22:21451131-21451151 | absent | 23119 | 3' UTR |
| GGGTGGGGATGGGGTGAGGG | chr22:48336509-48336528 | present | 100422916 | Intergenic |
| GGGTTGGGGGTGGGGGAGGG | chr2:10264610-10264629 | present | 3241 | Intergenic |
| GGGTTGGGGGTGGGATGGGG | chr2:134334522-134334541 | absent | 4249 | Intron |
| GGGTGGGGATGAGGGTGAGGG | chr2:136030238-136030258 | present | 101928243 | Intergenic |
| GGGTGGGGATGGGGAGAGGG | chr2:186884280-186884299 | present | 151112 | Intergenic |
| GGGGATGGGGGAGGGATGGGG | chr2:235026100-235026120 | present | 23677 | Intron |
| GGGTTGGGGTTGGGGATGGGG | chr2:237474553-237474573 | present | 79083 | Intergenic |
| GGGTGGGGGGAGGGGGAGGG | chr2:71452928-71452947 | present | 8291 | Promoter |
| GGGTTGGGAGGTGGGGGAGGGG | chr2:80306685-80306706 | absent | 1496 | Promoter |
| GGGTGTGGGTGAGGGTGAGGG | chr3:105648950-105648970 | absent | 214 | Intergenic |
| GGGTGGGGGTGGGGAGAAGGG | chr3:14867703-14867723 | absent | 152273 | Intron |
| GGGGTAGGGGTAAGGGATGGGG | chr3:170871296-170871317 | absent | 200916 | Promoter |
| GGGTCCGGGGTGGGGTGGGG | chr3:183908290-183908309 | present | 100616127 | Intergenic |

**Supplemental Table 1. (Continued).**

| Sequence | Location | Experimental Evidence | Gene ID | Anntoation |
|---|---|---|---|---|
| GGGTTGGGGGAGGGATGGGG | chr3:186926653-186926672 | present | 6480 | Promoter |
| GGGCTGGGGGTTGGGTGGGG | chr3:50367106-50367125 | present | 11068 | Promoter |
| GGGTATTGGGGTGGGGTGGGG | chr3:73674103-73674123 | present | 23024 | Intergenic |
| GGGTGGTGGGTGGGGTGGGG | chr3:9937507-9937526 | present | 78987 | Promoter |
| GGGTGGGGTGGGGTGAGGGG | chr4:13540437-13540456 | present | 579 | Promoter |
| GGGTTGGGGGTGGGGCAGGG | chr4:1780111-1780130 | absent | 2261 | Intergenic |
| GGGTTGGGGGTGGGGCAGGG | chr9:99297662-99297681 | present | 100996569 | Intergenic |
| GGGGTGGGGGTAGGGAGGGG | chr4:185101976-185101995 | present | 291 | Intron |
| GGGCTGGGGCTGGGGTGGGG | chr4:3820535-3820554 | present | 152 | Intergenic |
| GGGTCGGGGGTGGGGCGAGGG | chr4:94451863-94451883 | absent | 10611 | Promoter |
| GGGGAGGGGGAGGGATGGGG | chr5:140141772-140141791 | present | 101929719 | Intergenic |
| GGGTTGGGGGCAGGGTTGGGG | chr5:64122940-64122960 | absent | 285671 | Intergenic |
| GGGGTGGGGGAAGGGAAGGG | chr6:35743073-35743092 | present | 221481 | Intron |
| GGGTTGGGGGTTGGGGGTGGGG | chr6:43679847-43679868 | present | 55168 | Intron |
| GGGATGGGGGTGGGGGAGGG | chr6:57178779-57178798 | present | 101927211 | Promoter |
| GGGGTGGGGGAGGGACGGGG | chr7:100472520-100472539 | present | 81628 | Promoter |
| GGGCAGGGGGTGGGGGAGGG | chr7:128832311-128832330 | absent | 2318 | Promoter |
| GGGTTGGGGAGAGGGATGGG | chr7:149751402-149751421 | absent | 84626 | Intergenic |
| GGGTTGGGGGAGTGGGAGGG | chr7:26621451-26621470 | present | 285941 | Intergenic |
| GGGTTTGGGGAGGGAAGGGG | chr7:4482507-4482526 | absent | 221937 | Intergenic |
| GGGTGGGGGCGGGGGGAGGG | chr8:101761018-101761037 | present | 83988 | Intron |
| GGGGTGGGGGCGGGGGAGGG | chr8:113439774-113439793 | present | 114788 | Promoter |
| GGGTTTGGGGTGGGGTGGGGG | chr8:130519615-130519635 | present | 50807 | Intergenic |
| GGGTGTGGGGGTTGGGGGAGGG | chr8:142753544-142753565 | absent | 137797 | Promoter |
| GGGCCGGGGGTGGGGGAGGG | chr8:25689097-25689116 | absent | 64641 | Intergenic |
| GGGTTGGGGCTTGGGGAGGG | chr8:26653400-26653419 | present | 1808 | Intron |
| GGGTTGGGGGTGGGGTGGGGG | chr9:114098608-114098628 | present | 113220 | Promoter |
| GGGTTGTGGGTGGGGATGGGG | chr9:116931301-116931321 | present | 22954 | Intron |
| GGGATTGGGGATGGGGTGGGG | chr9:134806401-134806421 | present | 1289 | Promoter |
| GGGGTTGGGGGTGGGGAGGG | chr9:91299523-91299542 | present | 549 | Intron |
| GGGTGGGCGCGGGGTGAGGG | chr9:98745018-98745037 | absent | 203286 | Intron |
| GGGTTAGGGGGAGGGGTGGGG | chrX:120755247-120755267 | present | 643311 | Intergenic |

**Supplemental Table 2. Summary of Family 79 G4 sequences.**

| Sequence | Location | Experimental Evidence | Gene ID | Annotation |
|---|---|---|---|---|
| GGGAGGGGAGGGGAGGGG | chr1:11692947-11692964 | present | 374946 | Promoter |
| GGGAGGGGAGGGGAGGGG | chr1:32936592-32936609 | present | 127544 | 3' UTR |
| GGGAGGGGAGGGGAGGGG | chr10:12887513-12887530 | present | 83643 | Intergenic |
| GGGAGGGGAGGGGAGGGG | chr11:47595913-47595930 | present | 114900 | Promoter |
| GGGAGGGGAGGGGAGGGG | chr13:99003017-99003034 | present | 23348 | Intron |
| GGGAGGGGAGGGGAGGGG | chr16:54930945-54930962 | present | 10265 | Promoter |
| GGGAGGGGAGGGGAGGGG | chr4:151016295-151016312 | present | 987 | Promoter |
| GGGCGGGGCGCGGGCGGGG | chr10:133262048-133262066 | present | 101 | Promoter |
| GGGCGGGGCGCGGGCGGGG | chr10:133262092-133262110 | present | 101 | Promoter |
| GGGGCGGGGAGGGGCGGGG | chr10:14604436-14604454 | absent | 83641 | Promoter |
| GGGGCGGGGCGGGGCGGG | chr10:17348902-17348919 | absent | 338596 | Intron |
| GGGGCGGGGCGGGGCGGG | chr11:533399-533416 | present | 3265 | Promoter |
| GGGGCGGGGCGGGGCGGG | chr14:89701705-89701722 | absent | 1112 | Intron |
| GGGCGGGAGGGGCGGGG | chr10:3172964-3172980 | present | 10531 | Promoter |
| GGGGAGGGGCGGGGGGGG | chr11:115164500-115164517 | present | 23705 | Intergenic |
| GGGGCGGGGGGGGGCGGG | chr11:134401965-134401982 | present | 27087 | Intron |
| GGGAGGGGCAGGGGCAGGG | chr11:2571906-2571924 | absent | 3784 | Intron |
| GGGAGGGGCAGGGGCAGGG | chr5:149676067-149676085 | present | 389337 | Intergenic |
| GGGCGGGGCAGGGGCGGG | chr11:2902105-2902122 | present | 5002 | Promoter |
| GGGCGGGGAAGGGTGGGG | chr11:2902676-2902693 | present | 5002 | Promoter |
| GGGGGGGAGGGGCGGGG | chr11:64342922-64342938 | present | 283234 | Promoter |
| GGGAGGGGCCGGGGCGGGG | chr11:6473947-6473965 | present | 10612 | Promoter |
| GGGCGGGGCGGGGCGGGG | chr11:65558635-65558652 | absent | 4054 | Promoter |
| GGGCGGGGCGGGGCGGGG | chr21:45555638-45555655 | present | 6573 | Intergenic |
| GGGCGGGGCGGGGCGGGG | chr9:124777075-124777092 | absent | 169611 | Promoter |
| GGGAGGGGCGGGGCGGG | chr11:7020387-7020403 | present | 7761 | Promoter |
| GGGAGGGGCGGGGCGGG | chr16:371393-371409 | present | 10573 | Promoter |
| GGGAGGGGCGGGGCGGG | chr19:17747734-17747750 | present | 23149 | Promoter |
| GGGAGGGGCGGGGCGGG | chr19:44847716-44847732 | present | 5819 | Promoter |
| GGGAGGGGCGGGGCGGG | chr9:70258881-70258897 | absent | 100507299 | Promoter |
| GGGGCAGGGGCAGGGCGGG | chr11:79081753-79081771 | present | 26011 | Intron |
| GGGGAAGGGGGGGCGGGG | chr12:111597150-111597167 | present | 6311 | Promoter |
| GGGCGGGGCGGGGCGGG | chr12:48350921-48350937 | absent | 121274 | Promoter |
| GGGGCGGGGCGGGGGGG | chr14:105441449-105441465 | present | 9112 | Promoter |
| GGGGCGGGGCGGGGGGG | chr5:70126581-70126597 | absent | 6606 | Intergenic |
| GGGGCGGGGCGGGGGGG | chr5:71126892-71126908 | absent | 728340 | Intergenic |
| GGGGCGGGGCGGGGGGG | chr7:99770423-99770439 | present | 1576 | Intron |
| GGGGCGGGGCGGGGGGG | chrX:132751250-132751266 | present | 100874102 | Intron |
| GGGCGGGGCGGGGGCGGG | chr14:22871353-22871370 | present | 26020 | Promoter |
| GGGCGGGGCGGGGGCGGG | chr22:37519650-37519667 | absent | 29775 | Promoter |
| GGGGCGGGGCGGGCGCGGG | chr14:55052093-55052111 | present | 93487 | Promoter |
| GGGCGGGGCGGGGCGCGGG | chr14:99645483-99645501 | absent | 84439 | Promoter |
| GGGGCGGGGCGGGGCGGGG | chr1:3718211-3718229 | absent | 7161 | Intron |
| GGGGCGGGGCGGGGCGGGG | chr15:101724648-101724666 | present | 123283 | Promoter |
| GGGGCGGGGCGGGGCGGGG | chr17:62808389-62808407 | absent | 162333 | Promoter |
| GGGGCGGGGCGGGGCGGGG | chr2:19348311-19348329 | present | 100616307 | Promoter |
| GGGGCGGGGCGGGGCGGGG | chr22:15761338-15761356 | absent | 106146148 | Promoter |
| GGGGCGGGGCGGGGCGGGG | chr22:42500547-42500565 | present | 94009 | Promoter |
| GGGCGGGGAGGGGGCGGG | chr15:39366098-39366115 | present | 400360 | Intron |
| GGGAGGGGACGGGCGGGG | chr15:69298887-69298904 | absent | 54852 | Promoter |
| GGGTGAGGGGGCGGGCGGG | chr15:88257743-88257761 | absent | 4916 | Promoter |
| GGGAGGGTGAGGGGAGGGG | chr15:92439783-92439801 | present | 8128 | Intron |
| GGGCAGGGGAGGGGCGGG | chr16:21511745-21511762 | present | 100500917 | Intergenic |
| GGGCAGGGGAGGGGCGGG | chr16:87860829-87860846 | present | 8140 | Promoter |
| GGGTGAGGGGCGGGCGGG | chr16:22008068-22008085 | present | 730094 | Promoter |
| GGGGAGGGGGCGGGTCGGG | chr16:66604580-66604598 | present | 123920 | Promoter |
| GGGCGGGGAGGGCGCGGG | chr16:706088-706105 | absent | 146330 | Promoter |
| GGGCGGGGAGGGCGCGGG | chr16:67530169-67530186 | present | 79567 | Promoter |
| GGGAGCGGGAGGGGCGGGG | chr16:8847828-8847846 | present | 5373 | 3' UTR |
| GGGCGGGACGGGGCGGGG | chr16:88686619-88686636 | present | 333929 | Promoter |
| GGGCGGGGTAGGGCGGGG | chr16:9091577-9091594 | present | 29035 | Promoter |
| GGGGAGGGGGCGGGGGGG | chr17:3635824-3635841 | present | 23729 | Promoter |
| GGGGAGGGGGCGGGGGGG | chr17:65055061-65055078 | present | 10672 | Promoter |
| GGGGAGGGGCGCGGGCGGG | chr17:43755320-43755338 | absent | 50964 | Promoter |
| GGGCGGGGGGGGCGGG | chr17:518280-518295 | present | 55275 | Promoter |
| GGGCGGGGACGGGGCGGGG | chr17:64130213-64130231 | present | 2081 | Promoter |

**Supplemental Table 2. (Continued).**

| Sequence | Location | Experimental Evidence | Gene ID | Annotation |
|---|---|---|---|---|
| GGGGCGGGGTGGGGCGGG | chr17:74380229-74380246 | present | 350383 | Intergenic |
| GGGTGAGGGCGGGGCTGGG | chr17:75740069-75740087 | absent | 3691 | Promoter |
| GGGGAAGGGGGCGGGGGGG | chr17:81067374-81067392 | present | 10458 | Promoter |
| GGGCCGGGCGGGGCGGG | chr17:81716621-81716637 | present | 1468 | Promoter |
| GGGGGGGCAGGGGCGGGGG | chr17:9825847-9825865 | present | 9340 | Promoter |
| GGGCAGGGGCAGGGCGGGG | chr18:37517402-37517420 | present | 56853 | Intron |
| GGGGCGGGGCGGGCCGGG | chr19:1418556-1418573 | present | 26528 | Promoter |
| GGGCGGGGAGGGGCGGG | chr19:1940858-1940874 | absent | 1455 | Promoter |
| GGGCGGGGCAGGGCGGGG | chr19:3557619-3557636 | absent | 126321 | Promoter |
| GGGAGGGTGAGGGGCGGGG | chr19:45497095-45497113 | present | 6253 | Promoter |
| GGGCGGGGAGGGGCGGGG | chr1:180632001-180632018 | present | 9213 | Promoter |
| GGGCGGGGAGGGGCGGGG | chr19:49388163-49388180 | present | 147872 | Promoter |
| GGGGGGGAAGGGCGGGG | chr19:49556137-49556153 | present | 51070 | Promoter |
| GGGCGGGGCCGGGGCGGG | chr19:52690611-52690628 | present | 55769 | Promoter |
| GGGGCGGGCGGGGCGGG | chr19:676544-676560 | absent | 10272 | Promoter |
| GGGGAGGGGGGCGGGGGGG | chr1:10856271-10856289 | present | 54897 | Intergenic |
| GGGGAGGGGCGGGGCAGGG | chr1:110161522-110161540 | present | 388662 | Intron |
| GGGGCGGGGCGGGGTGGG | chr1:156677373-156677390 | present | 10763 | Promoter |
| GGGGCGGGGCGGGGTGGG | chr3:141402686-141402703 | present | 253461 | Promoter |
| GGGGGGGCGGGGCGGGG | chr1:209825787-209825803 | present | 27042 | Promoter |
| GGGCGGGAAGGGCGGGG | chr1:228276035-228276051 | present | 84033 | Promoter |
| GGGCGGGGGAGGGGCGGGG | chr1:33182128-33182146 | absent | 55223 | Promoter |
| GGGGTGGGGGGGGCGGGG | chr1:42731825-42731842 | present | 149461 | Downstream |
| GGGGAGGGCAGGGCGTGGG | chr21:40155906-40155924 | absent | 100616148 | Intron |
| GGGGCGGGGCCGGGGCGGG | chr21:42975112-42975130 | present | 5316 | Promoter |
| GGGCGGGGAGGGGAGGGG | chr22:27554681-27554698 | present | 4330 | Intergenic |
| GGGGGGGGCGGGGCGGGG | chr22:33528130-33528147 | present | 9215 | Intron |
| GGGGCGGGGCGGGGCAGGG | chr22:44891552-44891570 | present | 23779 | Intron |
| GGGGAGGGCTGGGCTGGGG | chr22:44916501-44916519 | present | 112885 | 5' UTR |
| GGGAGGGGCAGGGCGGGG | chr2:11130389-11130406 | present | 285150 | Promoter |
| GGGAGGGGAGGGGCGGG | chr2:131005683-131005699 | present | 50649 | Intron |
| GGGACGGGGCGGGGCGGG | chr2:197785363-197785380 | present | 66037 | Promoter |
| GGGGTGGGGCGGGGCGGGG | chr2:231683502-231683520 | present | 5757 | Intergenic |
| GGGGAGGGGGCGGGGCGGG | chr2:44778533-44778551 | present | 79823 | Intergenic |
| GGGGAGGGCGGGTGCTGGG | chr3:127140774-127140792 | present | 285311 | Intergenic |
| GGGAGGGGAGGGGCGGGG | chr3:13018647-13018664 | present | 9922 | Intron |
| GGGAGGGGAGGGGCAGGG | chr4:1766503-1766520 | absent | 10460 | Intergenic |
| GGGCGGGGCGGGCGGGG | chr4:2418706-2418722 | absent | 57732 | Promoter |
| GGGGAGGGGCAGGGCTGGG | chr4:3779401-3779419 | present | 152 | Intergenic |
| GGGCGGGAAGGGGCGGGG | chr4:41360750-41360767 | present | 22998 | Promoter |
| GGGAGGGGCAGGGGCGGGG | chr4:6782547-6782565 | absent | 9778 | Promoter |
| GGGCGGGGAGGGGGCGGGG | chr5:179795444-179795462 | present | 4056 | Promoter |
| GGGGAGGGGCAGGGCGGGG | chr5:2011684-2011702 | present | 101929081 | Intergenic |
| GGGATGGGGAGGGGCGGGG | chr5:72299335-72299353 | present | 23107 | Intron |
| GGGAGGGGAAGGGGCGGGG | chr6:34223494-34223512 | present | 3159 | Intergenic |
| GGGGAGGGGAGGGCGCGGG | chr7:148884873-148884891 | absent | 2146 | Promoter |
| GGGGAGGGGGGGGCGGGG | chr7:151520014-151520031 | present | 6009 | Promoter |
| GGGCGGGGAGGGCCGGGG | chr7:1669902-1669919 | present | 392617 | Intergenic |
| GGGCAGGGCGGGGCGGG | chr7:2096544-2096560 | present | 8379 | Intron |
| GGGTAGGGGCGGGGCGGGG | chr7:75410724-75410742 | absent | 378108 | 3' UTR |
| GGGCGGGGAAGGGCGGGG | chr7:7566848-7566865 | absent | 54468 | Promoter |
| GGGCGGGGCGGGGCCGGG | chr7:99375518-99375535 | absent | 10095 | Promoter |
| GGGGAGGGGTCGGGGGGG | chr8:133453251-133453268 | present | 6482 | Downstream |
| GGGAGCGGGGCGGGGCGGG | chr8:26383646-26383664 | present | 665 | Promoter |
| GGGAAGGGGCAGGGCGGGG | chr9:1046289-1046307 | present | 102800446 | Promoter |
| GGGGGGGCGGGGCGGG | chr9:113012640-113012655 | present | 169834 | Promoter |
| GGGGCGGGGCGGGGCCGGG | chr9:124853507-124853525 | absent | 401551 | Promoter |
| GGGCGGGGAGGGCCGGG | chr9:133557493-133557509 | present | 9719 | Intron |
| GGGGCGGGGTCGGGGCGGG | chr9:5629030-5629048 | present | 57589 | Promoter |
| GGGGCGGGCCGGGGCGGG | chr9:91423955-91423972 | present | 4783 | Promoter |
| GGGGCGGGCGCGGGGCGGG | chrX:153411372-153411390 | present | 139735 | Intergenic |
| GGGCGGGGCGGGGCGAGGG | chrX:154546963-154546981 | present | 2539 | Promoter |
| GGGCGGGCCGGGGCGGGG | chrX:155458643-155458660 | absent | 8263 | Promoter |
| GGGGAGGGGCGGGGCGGGG | chrX:76427616-76427634 | present | 57692 | Promoter |

**Supplemental Table 3. Sequence repeats capable of forming multiple G4 structures.**

| Location | Gene ID | Gene Symbol | Sequence |
|---|---|---|---|
| chr4:1318076-1318118 | 10296 | MAEA | GGGGGAAGCCGGGCACACGGGGCCAGGGAGGCGGGTGGGTGGG |
| chr4:2756374-2756472 | 79155 | TNIP2 | GGGCGGGGCTGCGCGGGGAAGGGCGGGGCTGCGCGGGGGGCGGGGCTGCGTGGGGGAGGGCGGGGCCGGGCTGTTGT |
| | | | GGGTGGGCGGGGCGCACCGGGG |
| chr4:49572959-49573006 | | | GGGGGGTGGGGAGGGTTGGGGGGATTAAGGGGTGGGGAGGGTTGGGGG |
| chr5:1295088-1295155 | 7015 | TERT | GGGGAGGGGCTGGGAGGGCCCGGAGGGGGCTGGGCCGGGGACCCGGGAGGGGTCGGGACGGGGCGGGG |
| chr5:2204989-2205021 | | | GGGAAGTGGGGGTGGGGCTGGGGCTTGGGCGGG |
| chr7:101017720-101018062 | 102724094 | LOC102724094 | GGGGGAAGGGAAGGGGTCCCAGGGGGAGGGAGGGAGTCCCAGGGGGAAGGGAAGGGGTCCCAGGGGGAGGGAGGGA |
| | 140453 | MUC17 | GTCCCAGGGGGAAGGGAAGGGGTCCCAGGGGGAGGGAGGGAATCCCAGGGGGAAGGGAAGGGGTCCCAGGGGGAGG |
| | 10071 | MUC12 | GAGGGAGTCCCAGGGGGAAGGGAAGGGGTCCCAGGGGGAGGGAGGGAGTCCCAGGGGGAAAGGAAGGGGTCCCAGG |
| | | | GGGAGGAAGGGAGTCCCAGGGGGAAGGGAAAGGGAAGGGGTCCCAGGGGGAAGGGAGGGAGTCCCAGGGGGAGAAG |
| | | | GGAGTCCCAGGGGGAGAAGGGAGTCCCGGGGGAAGAGGG |
| chr7:129978853-129978883 | | RP11-306G20.1 | GGGGGTGGGGGTGGGTGTGGGAGGGCAGGGG |
| chr7:149018792-149018849 | 9601 | PDIA4 | GGGGTGGGGGTGGGGGGGTAGGAGGGGGGAGTAGTGGGGTGGGCAGGGTGGGTTGGGG |
| chr7:26864641-26864689 | 8935 | SKAP2 | GGGCCGGGGCGGGGAGATGGGTGGGAAGGGACACGAAGGGCCTGAGGGG |
| chr7:44224942-44224975 | 816 | CAMK2B | GGGAGGGGCTGGGCAGGGCTGGGAAAGGGGTGGG |
| chr7:74218035-74218071 | 7462 | LAT2 | GGGGGCTGGGGGTGGGCAGGGCCTGAGGGGAGAGGGG |
| chr8:142348488-142348542 | 203062 | TSNARE1 | GGGGCAGCCTGGGGCGGGAGCGGGGGCCAGGGGAGGGTGGGCATGGGGTGCCGGG |
| chr8:142348556-142348610 | 203062 | TSNARE1 | GGGGCAGCCTGGGGCGGGAGCGGGGGCCAGGGGAGGGTGGGCATGGGGTGCCGGG |
| chr8:144465691-144465773 | 90990 | KIFC2 | GGGGAGAAGGGCCGGGGCGGGGCTGCGAGGGGCGGGGTCTGGGCGGGGCTGCGAGGGGCGGGGGGTCTGGGCGGGG |
| | 50626 | CYHR1 | CTGAGGG |
| chr9:32956186-32956230 | | | GGGAGGGTCCGGGAAGGGTCCCTGGGTGGGGGAGGGGGAAAGGGG |
| chr9:39145609-39145636 | 79937 | CNTNAP3 | GGGGGCTGGGGGCATGGGGAGGGTAGGG |
| chr9:5840502-5840539 | | | GGGGGCCGGGGAGGGGCAGTGGGCATGGGTAGGGAGGG |
| chr9:93210282-93210324 | 65268 | WNK2 | GGGGTGAGGGATGGGCAGGGTGGGCAGGGATGGGGGACTGGGG |
| chrX:107610224-107610250 | | | GGGGGGCAGGGGCAGGGAAGGGGAGGG |
| chrX:12711746-12711782 | 9758 | FRMPD4 | GGGGACTTGGGGCGGGGGGCAGGGTTGGGGGGAAGGG |
| chr1:11971187-11971260 | 5351 | PLOD1 | GGGGCAGGGGGATGGGGTGGGAGGGGTAGGGTGGAGTGGGGGGCTTGGGTGGAAGGGCCAGGGGTGGGTGGGGG |
| chr1:38005556-38005593 | 2275 | FHL3 | GGGCTGGGGGGCCGGGGGCGGGGTCCGGGCGGGGCGGG |
| | 51118 | UTP11 | |
| chr10:131738939-131738977 | | | GGGGTGGGTGGGGAGGGAGCATGGGGTGGGCAGGGTGGG |
| chr10:44647571-44647613 | | | GGGTGAGGGGGATGGGTGGGGGATGGGCAGGGTAGGGCAGGGG |
| chr10:86347049-86347092 | 2894 | GRID1 | GGGTGGGTGGGGCAGGGCAGGAGGGTGGGGCTGGGCAGTTAGGG |
| chr11:119695264-119695328 | 5818 | NECTIN1 | GGGGACTGGTGGGGAGGGTGGGGACCTGGGAGGGGTGGGAGAGGGAGATGGGAATATGGGCAGGG |
| chr11:63562894-63562932 | 54979 | HRASLS2 | GGGAGGGATTAGCTGGGGAGGGAGGGTCCAGGGAAAGGG |
| chr12:123034471-123034509 | 57605 | PITPNM2 | GGGTCGTGGGGCAAGGGTGGGTCATGGGGTGAGGGTGGG |
| chr12:47939348-47939389 | 7421 | VDR | GGGTGGGGCTTGGGGGAGGTGGGTCCTGGGGTGGGGATGGGG |
| chr12:53499343-53499404 | 6895 | TARBP2 | GGGGGGGTGGGGGGCAGGGATGGGTCTGGGTCCTGGGATCCGGGCGTGGAGGGAGGGTCGGG |
| | 7786 | MAP3K12 | |
| chr14:23386616-23386678 | 4624 | MYH6 | GGGAGGCCTGGGAAGGGGTGGGGCGAGGGCGGGCAGACAGGGCACAGGGCAGGGTTGAGAGGG |
| chr16:3957697-3957730 | 115 | ADCY9 | GGGATGGGGGTCCTGGGAGGGCAGGGCTAGGGGG |
| chr17:50604073-50604120 | 8913 | CACNA1G | GGGGGTGGGGAGCAGGGTCAAGGGACAAGGGAGGGTCTGGGCTGGGGG |
| chr17:82698358-82698455 | 10966 | RAB40B | GGGTGAGCGCGGGCGGAGGGCGTGCCGGGGTGCGGGCGCGGGGCCGGGGAGGGGCGCGGGGCTGGGGAGGGGGTGC |
| | | | GGGTGGGGGTCCGGGTCCGGGG |

**Supplemental Table 3. (Continued).**

| Location | Gene ID | Gene Symbol | Sequence |
|---|---|---|---|
| chr18:79395460-79395704 | 4772 | NFATC1 | GGGGGGGCGCACGGGGAGGGGGGGGCGCACGGGGAGGGGGGGCGCCCGGGGAGGGGGGCGCCCGGGGAGGGGGGGCGCACGGGGAGGGGGGCGCACGGGGAGGGGGGGCGCACGGGGAGGGGGGCGCACGGGGAGGGGCGCACGGGGAGGGGATGGGGGCGTAGGGGCGGGAACGGGGAATCCGGGGGCCGGGCAGGGGGGCGCTGGGGCTGGCGGGGGAAACGGGGGCGAACGGGCCAGACGGG |
| chr18:79617457-79617512 | | | GGGGGGCGGGGTCCTGGGGGGGCAGGGTCCCGGGGAGGGCGGGGTCCCGGGGAGGG |
| chr19:6373235-6373294 | 84266 | ALKBH7 | GGGGTTGTGGGGCCAGGGGGGTCGGGCGCAGGGATGGGGCGGGGCCACGCTGGGGCGGGG |
| chr19:9851039-9851068 | | | GGGGGTGAGGGGGTGGGTGGGAGGTGAGGG |
| chr20:1225680-1225711 | 642636 | RAD21L1 | GGGACCGGGGGCAGGGGGCGGGGAAGGGCGGG |
| chr21:40006234-40006276 | | | GGGCCTGGGGAGAGGGAGGGCCTGGGGAGAGGGAGGGACTGGG |
| chr21:45287489-45287569 | 23275<br>642852 | POFUT2<br>LOC642852 | GGGGCAGGGGCCAGGGGGATGGGATGGAGCGGGGTCAGGGGGCAGGGGTCAGGGGAATGGGATGGGGTCAGGGTTCTGGGG |
| chr22:43280423-43280510 | 101927447<br>80274 | Z99756.1<br>LOC101927447<br>SCUBE1 | GGGTGAGGGGAAGGGACGGGGATGGGTGAGGGGAAGGGACGGGAGGATGGGTGAGGGGAAGGGACAGGGGGATGGGTGAGGGGAAGGG |
| chr22:49037526-49037614 | | | GGGAGCGGGAGGGGCCAGGGGGTGGGACGGGGCGGGGAGAGGGAGAAGAGGGGTTCTGGGTGGGAAGGATTGGGGAGCGGGAGGAGGGG |
| chr3:129606851-129606910 | 23129 | PLXND1 | GGGCGGCCAGGGGCAGGCGGGGGTCCCGGGGGCGGGCGGGGCCGGGGCGGGGAGTGAGGG |
| chr4:88699286-88699364 | 8916<br>266812 | HERC3<br>NAP1L5 | GGGCAGGGTGGGGAAGAAGAGGGTGGGGCTGCGTGGGTGGGTGGGGAGGAAGAGGGTGGGGCTGGGTGGGTGGGTGGGG |
| chr5:524556-524624 | 6550 | SLC9A3 | GGGCTCCGGGGGAGGGTGGGCACCAGGGAGCGCGGGGTGGGCGTGCGCGGGCGGGGCGGGCGTGCCGGG |
| chr6:34486149-34486233 | 29993 | PACSIN1 | GGGGGTGAGGGGTGGAGGGACAGGGGGCCTGGGAACCCAGGGAGAGGGAGGCAGGGCCTAGGGGTGGGGTGAGGTGGGTTTGGGG |
| chr6:44222239-44222286 | 2030 | SLC29A1 | GGGCTGGCGGGGATGTGGGGGATGGGGGTGGGGTGGGGGAGGGTTGGG |
| chr7:157138531-157138583 | 9690 | UBE3C | GGGCCGGGATGGGGTGCAGGGCAGGGTGCGGGGTGCAGGGTGGGGTGCAGGGG |
| chr8:142348624-142348668 | 203062 | TSNARE1 | GGGGCGGGAGCGGGGGCCAGGGGAGGGTGGGCATGGGGTGCCGGG |
| chr8:142464060-142464098 | 575 | ADGRB1 | GGGGGCAGGGAGGCGGGGCAAGGGTGGGATGGGAGAGGG |
| chr8:26383526-26383612 | 665 | BNIP3L | GGGCGGGGCGGGGCGGGGGCGGGCCTGGGGGGCGGGGAGGCCGGGTGGGCGGAGCGGGCCGCGGAGGGGACGTGGGCCGGGATGGGG |
| chr9:132589622-132589662 | 56751 | BARHL1 | GGGGGGCACTGGGCTGGGGCGCCAGGGAGGGCCGGGCAGGG |
| chr9:136800509-136800550 | 84960<br>55684 | RP11-216L13.19<br>CCDC183<br>RABL6<br>RP11-216L13.18 | GGGAAGGGCGGGGGTCAGGGGCTGGGATCTGGGAGGGGCGGG |
| chrX:120157742-120157823 | 727940 | RHOXF2B | GGGGTGGGGGGAGTAGGGCGGGGGAGGGAGTAGGGCGGGGGGGCGTAGGGTGGAGGGGGGGAGTAGGGCGGGGGGGCCGGGG |
| chr1:2640997-2641101 | 100287898 | TTC34 | GGGGCAGGGAAGCGGGGTGTGGGGAGGGCAGGGAAGGGGGGTGTGGGGAGGGCTGGGAAGGGAGGTATGGGGAGGGCTGGGAAGGGAGGTATGGGGAGGGCTGGG |
| chr1:29727790-29727836 | | | GGGTGGGCTTGGGGAGGGGTGGAGGGAAGGGTGGGCTGAGGGAGGGG |
| chr14:103109168-103109254 | 91828 | EXOC3L4 | GGGAGGGAGACACGGGGACAGGGTGGAGAGGGAGGGAGGGAGGGAGACATGGGGACAGGGTGGCGAGGGAGAGAGGGAGACCGGGGG |
| chr14:103109565-103109722 | 91828 | EXOC3L4 | GGGAGACCCAGGGACAGGGTGGAGAGGGAGGGAGGGAGGGAGACATGGGGGCAGGGTGGAGAGGGAGGGAGGAGGGAGACATGGGGGCAGGGTGGAGAGGGAGGGAGGAGGGAGACATGGGGGCAGGGTGGAGAGGGAGGGAGGAGGGAGACATGGGG |
| chr16:47887333-47887374 | 101927132<br>100507534 | RP11-523L20.2<br>LINC02133<br>LINC02192 | GGGTCACAGGGTCAGGGAGGGGGCTGCGGGTGGGAGGCAGGG |

**Supplemental Table 3. (Continued).**

| Location | Gene ID | Gene Symbol | Sequence |
|---|---|---|---|
| chr19:38390491-38390536 | 399473<br>199720 | SPRED3<br>GGN | GGGGCATGCGGGGAGGGTAGGGACCTGGGGAGGGAGGGGAGAGGGG |
| chr2:129877629-129877660 | | | GGGGAGAGGGGCGGGGGCGGGGCCGGGCCGGG |
| chr21:43075944-43076011 | 875 | CBS | GGGGTGGGGAAGGGGTGGGGGGGAGGGGCCCGGGCTGGGTGGGGTGGAGGAGGGGCTGGGGGGGCGGG |
| chr22:44939685-44939728 | 112885 | PHF21B | GGGGATGGGTGGGAGCAGGGCTAGGGAGGGGCGAAGGGATAGGG |
| chr3:50204812-50204840 | 10991 | SLC38A3 | GGGGTGGGGTGGGGGCGAGGGTGGGAGGG |
| chr8:10852146-10852235 | | | GGGAGGGGAAGGGGAGGGCAGGGAAGAGAAGGGTCAGCACGGGGAAGAGAAGGGGAGCGCGGGGAAGGGAAGGGTCAGCGCGGGGAAGGG |
| chr9:127809879-127809969 | 2356 | FPGS | GGGGCGGGATCTTGGGGAAGGGCGGGGCGGGGTCGTGGGGAAGGGCGGGGGCGGGCCCATGGGGAGGGGCGGGGTCGTGGGCGGGGACGGG |
| chr9:135757071-135757140 | 57582 | KCNT1 | GGGGTATCAGCGGGGGCGATGGAGGGTGGGGGTGGGGCCCAGCAGGGGAGGGGCAGGGTGGGGAAGAGGG |
| chrX:154065081-154065110 | 4204 | MECP2 | GGGGTGGGTGGGGTGGGGGCCGGGGAAGGG |
| chr5:181098279-181098403 | | | GGGGGAAGGGACTGGAGGGGAGGGGAGGGGAGGGGACGGGTGGGGAGGGGTGGGGCGGGGAGGGAAAGGGTGGGGAGCAGAGGGGTGGGGAGGGGAGGGGTGGGGAGAGGTAGGGAGGGGAGGGG |
| chr7:76282546-76282601 | 222183 | SRRM3 | GGGGGCTGGGGGCGGGGAGGGTTCCCTGGGGCGGGGCTTAGGGCGGAGGGGCGGGG |

**Supplemental Table 4: Summary of Family 2 G4 sequences.**

| Sequence | Location | Experimental Evidence | Gene ID | Anntation |
|---|---|---|---|---|
| GGGGAGGGCCTGGGACAGGG | chr11:1648838-1648857 | absent | 387742 | Intergenic |
| GGGAGGGCCTTGGGACAGGG | chr1:226482764-226482783 | absent | 375057 | Intergenic |
| GGGAGGGGCCTGGGACAGGG | chr9:135803176-135803195 | present | 57582 | Intergenic |
| GGGGGAATGGGCTGGGACAGGG | chr1:7715171-7715192 | absent | 23261 | Intron |
| GGGAAGGGGGCTGGGAAAGGG | chr17:74059333-74059353 | absent | 6169 | Intron |
| GGGCTGGGCATGGGACAGGG | chr14:74084190-74084209 | present | 4329 | Promoter |
| GGGGAGTGGGCTGGGACAGGG | chr1:120652107-120652127 | absent | 101954277 | Intergenic |
| GGGGAGTGGGCTGGGACAGGG | chr1:149272171-149272191 | absent | 400818 | Intergenic |

**Supplemental Table 5. Summary of Family 3 G4 sequences.**

| Sequence | Location | Experimental Evidence | Gene ID | Annotation |
|---|---|---|---|---|
| GGGAGGGGGCTGCAGGGAGCTGGG | chr19:41700278-41700301 | present | 1087 | Intron |
| GGGAGGGGGCTGCAGGGAGCTGGG | chr19:41700314-41700337 | present | 1087 | Intron |
| GGGAGGGGGCTGCAGGGAGCTGGG | chr19:41700350-41700373 | present | 1087 | Intron |
| GGGAGGGGGCTGCAGGGAGCTGGG | chr19:41700386-41700409 | present | 1087 | Intron |
| GGGAGGGGGCTGCAGGGATGGGGG | chr12:124531250-124531273 | present | 9612 | Promoter |
| GGGAGGGGGCTGCAGGGATGGGGG | chr3:53193909-53193932 | absent | 5580 | Intergenic |
| GGGAGGGGCTGCAGGGATGGGGG | chr22:43426947-43426969 | absent | 758 | Intron |
| GGGAGGGGGAGGCAGGGTTGGGG | chr1:206041760-206041782 | absent | 440712 | Intron |
| GGGAGGGTGCTCCTGGGATGGGG | chr17:1312485-1312507 | present | 286753 | Intergenic |
| GGGAGGGGGCTTCTGGGGTGGGG | chr3:13852428-13852450 | present | 7476 | Intron |

**Supplemental Table 6. Summary of Family 4 G4 sequences.**

| Sequence | Location | Experimental Evidence | Gene ID | Annotation |
|---|---|---|---|---|
| GGGCCTGGGAGGGAAGGAGAGGG | chr4:3513681-3513703 | absent | 4043 | Intron |
| GGGCTAGGGTCGGGAGTAGAGGG | chr2:88972574-88972596 | absent | 100616399 | Intron |
| GGGGCTGTGGAGGGAGGGAGAGGG | chr15:41893960-41893983 | absent | 51332 | Promoter |
| GGGCTGGGGCGGGAAGGAGAGGG | chr1:121185193-121185215 | absent | 653464 | Promoter |
| GGGCTGGGGCGGGAAGGAGAGGG | chr1:143972835-143972857 | absent | 554282 | Promoter |
| GGGGCTGGGGCGGGAAGGAGAGGG | chr1:206203718-206203741 | present | 729533 | Promoter |
| GGGCAGGGCGAGGGATGGAGAGGG | chr17:39144386-39144409 | absent | 57125 | Intron |
| GGGCATGGGCGGGGTGGAGAGGG | chr3:143313988-143314010 | absent | 100885796 | Intron |
| GGGGTGGGGAGGGAATGTGAGGG | chr10:70886991-70887013 | absent | 5092 | Promoter |

**Supplemental Table 7. Summary of Family 32 G4 sequences.**

| Sequence | Location | Experimental Evidence | Gene ID | Annotation |
|---|---|---|---|---|
| GGGAAGGGGAAGGGACAGGG | chr1:1136393-1136412 | present | 254099 | Promoter |
| GGGGTGGGGTGGGGAGAGGG | chr1:161197724-161197743 | present | 4720 | Promoter |
| GGGCTGGGGTTGGGGCTGGG | chr1:201275493-201275512 | present | 5317 | Intergenic |
| GGGCTGGGGCTGGGGCAGGG | chr1:229251426-229251445 | absent | 5867 | 3' UTR |
| GGGGTGGGGTGGGGGATGGG | chr1:3110907-3110926 | present | 63976 | Intron |
| GGGCTGTGGGCGGGGCTAGGG | chr1:37735516-37735536 | present | 284656 | Promoter |
| GGGCTGGGAGAGGGCCTGGG | chr1:54138165-54138184 | absent | 200008 | Intron |
| GGGATGGGCATGGGGGAGGG | chr10:124644752-124644771 | present | 64077 | Intron |
| GGGGTGGGGGTGGGGTTGGG | chr10:130945693-130945712 | present | 100422867 | Intergenic |
| GGGATGGGCTGGGGGCTGGG | chr10:27715853-27715872 | present | 283078 | Intron |
| GGGGTGGGGTGGGGGCAGGG | chr10:69897023-69897042 | present | 1305 | Intron |
| GGGATGGGGCATGGGGAGGG | chr10:79375641-79375660 | present | 10105 | Intergenic |
| GGGCAGGGGGTGGGGCAGGG | chr11:404765-404784 | present | 11187 | Promoter |
| GGGCAGGGGGAGGGGGAGGG | chr11:64934344-64934363 | absent | 170589 | Promoter |
| GGGGTGAGGGTGGGGGTGGG | chr12:129184860-129184879 | present | 101927735 | Intron |
| GGGGTGGGGGATGGGGAGGG | chr12:51905879-51905898 | present | 94 | Promoter |
| GGGGTGGGGGAGGGGCAGGG | chr13:111296247-111296266 | present | 8874 | Intron |
| GGGCAGGGCTGGGTGGAGGG | chr13:31009892-31009911 | absent | 122046 | Intergenic |
| GGGCAGGGGTTGGGTGAGGG | chr14:24147068-24147087 | present | 5721 | Promoter |
| GGGTTGGGGGCGGGGGTGGG | chr14:65413539-65413558 | present | 2530 | Promoter |
| GGGGTGTGGGTGGGGCAGGG | chr14:95332851-95332870 | present | 101929080 | Promoter |
| GGGGAGGGGTGGGGACCGGG | chr14:96219408-96219427 | present | 623 | Intron |
| GGGATGGGGAGGGGACAGGG | chr15:77583296-77583315 | absent | 84894 | Intron |
| GGGCTGGGGAGGGGACAGGG | chr15:85258124-85258143 | absent | 11214 | Intergenic |
| GGGGAGGGCTGGGACCAGGG | chr15:90884045-90884064 | present | 2242 | Promoter |
| GGGCTGGGACAGGGCCAGGG | chr16:32301164-32301183 | absent | 729264 | Intergenic |
| GGGCAGGGCTGGGTCCAGGG | chr16:32302937-32302956 | absent | 729264 | Intergenic |
| GGGCTGGGACAGGGCCAGGG | chr16:33507434-33507453 | absent | 24150 | Intergenic |
| GGGCAGGGCTGGGTCCAGGG | chr16:33509201-33509220 | absent | 24150 | Intergenic |
| GGGGAGGGCATGGGGCAGGG | chr16:46623763-46623782 | present | 79801 | Promoter |
| GGGGTGGGGTTGGGGGAGGG | chr16:54970795-54970814 | present | 10265 | Intergenic |
| GGGCAGGGCTGGGAGAAGGG | chr17:10232070-10232089 | absent | 8522 | Intergenic |
| GGGAAGGGGAGGGGTCTGGG | chr17:58116701-58116720 | absent | 140735 | Intergenic |
| GGGCTGGGCCTGGGCCTGGG | chr17:61402103-61402122 | absent | 6909 | Promoter |
| GGGCGGGGCTGGGTCCTGGG | chr17:76079074-76079093 | present | 353174 | Promoter |
| GGGGTGGGGGTGGGGATGGG | chr17:77413847-77413866 | present | 10801 | Intron |
| GGGCTGGGGGAGGGGGTGGG | chr17:82435883-82435902 | present | 284004 | Promoter |
| GGGGTGGGGTGGGTGGAGGG | chr17:8743246-8743265 | absent | 146849 | Promoter |
| GGGCAGGGGCCTGGGGAGGG | chr18:10168387-10168406 | present | 9218 | Intergenic |
| GGGGGGGGGGTGGGGGTGGG | chr18:22548091-22548110 | present | 64693 | Intergenic |
| GGGCCCAGGGTGGGGCAGGG | chr18:37415961-37415980 | present | 56853 | Intron |
| GGGCCGGGGCTCTGGGCGGG | chr18:62596252-62596271 | absent | 54877 | Intergenic |
| GGGGTGGGATGGGGGCTGGG | chr19:17101093-17101112 | present | 4650 | Promoter |
| GGGCAGGTGGGTGGGCAGGG | chr19:42351769-42351788 | absent | 102465875 | Promoter |
| GGGAGGGGCGAGGGCCAGGG | chr19:51421895-51421914 | present | 89790 | Promoter |
| GGGTTGGGGGTGGGGGAGGG | chr2:10264610-10264629 | present | 3241 | Intergenic |
| GGGAAGGGGGTGGGAGAGGG | chr2:205734541-205734560 | absent | 8828 | Intron |
| GGGCAGGGACATGGGGTGGG | chr2:233976450-233976469 | present | 79054 | Promoter |
| GGGGTGAGGGTGGGGGTGGG | chr20:32612791-32612810 | present | 149950 | Intergenic |
| GGGTTGGGGGAGGGGGTGGG | chr20:32980815-32980834 | present | 140732 | Downstream |
| GGGGTGGGACAGTGGGAGGG | chr21:40286094-40286113 | present | 1826 | Intron |
| GGGTGGGGCTAGGGCCAGGG | chr22:21613160-21613179 | absent | 150223 | Intron |
| GGGGTGGGAGTGAGGGTGGG | chr3:11222231-11222250 | present | 3269 | Promoter |
| GGGCTGGGGCTGGGCCAGGG | chr3:125190497-125190516 | absent | 84561 | Promoter |
| GGGCTGGGGCAGGGGCCGGG | chr3:127823053-127823072 | absent | 11343 | Promoter |
| GGGGAGGGCATGGGGCAGGG | chr3:129088461-129088480 | absent | 2815 | 3' UTR |
| GGGCTGGGGGAGAGGGTGGG | chr3:129348117-129348136 | present | 339942 | Intergenic |
| GGGGGGGGGTGGGGGCAGGG | chr3:131466690-131466709 | present | 11222 | Promoter |
| GGGAGGGGCTGGGGCCTGGG | chr3:13628839-13628858 | absent | 2199 | Promoter |
| GGGCAGGGCTCGGGACAGGG | chr3:42732393-42732412 | absent | 100874114 | Promoter |
| GGGTAGGGAAAGGGAAAGGG | chr3:45979832-45979851 | absent | 79443 | Intron |
| GGGGTGGGGTAGGGGGAGGG | chr3:49489904-49489923 | present | 1605 | Promoter |
| GGGGTGGGGTGGGGGATGGG | chr3:49642027-49642046 | present | 8927 | Promoter |
| GGGGTGGGTGCGGGGCAGGG | chr4:102826891-102826910 | present | 7323 | Promoter |
| GGGGTGGGGAGGGGGCTGGG | chr4:141929124-141929143 | present | 3600 | Intergenic |
| GGGCAGGGGAATTGGGTGGG | chr4:152304991-152305010 | absent | 55294 | Intergenic |

**Supplemental Table 7. (Continued).**

| Sequence | Location | Experimental Evidence | Gene ID | Annotation |
|---|---|---|---|---|
| GGGCAGGGGTGGTGGGTGGG | chr5:113656482-113656501 | absent | 64848 | Intergenic |
| GGGGTGGGGCTTGGGGAGGG | chr5:134151255-134151274 | present | 6932 | 3' UTR |
| GGGGTGTGGGCGGGGCAGGG | chr5:151772043-151772062 | present | 10146 | Promoter |
| GGGCTGGGGCTAGGGGCGGG | chr5:168410001-168410020 | present | 23286 | Promoter |
| GGGCAGGGGCAGGGTGAGGG | chr6:157921891-157921910 | absent | 51429 | Intron |
| GGGCTGGGGCAGGGGGAGGG | chr6:166558569-166558588 | absent | 6196 | Intron |
| GGGCAGGGGGTGGGGGAGGG | chr7:128832311-128832330 | absent | 2318 | Promoter |
| GGGAGGGGCTGGGGGCTGGG | chr7:30915933-30915952 | absent | 358 | Promoter |
| GGGAGGGGCCGGGAGCTGGG | chr7:74454399-74454418 | absent | 9569 | Promoter |
| GGGGTGGGGGCGGGGGAGGG | chr8:113439774-113439793 | present | 114788 | Promoter |
| GGGCAGGGGTGGGGGGAGGG | chr8:127255311-127255330 | present | 100507056 | Intergenic |
| GGGAGGGGTTGGGGGCTGGG | chr8:144496661-144496680 | present | 84988 | Promoter |
| GGGCTGTGGGCGGGCCAGGG | chr8:144502891-144502910 | absent | 2875 | Promoter |
| GGGGTGGGGGAGGGGTTGGG | chr8:25517201-25517220 | present | 157313 | Intergenic |
| GGGGTGGGAGTAGGGGAGGG | chr8:38451245-38451264 | present | 2260 | Intron |
| GGGGTTGGGGTGGGGGAGGG | chr8:54466113-54466132 | present | 64321 | Intergenic |
| GGGCTGGGGTTGGGGAGGGG | chr8:99974271-99974290 | present | 26166 | Promoter |
| GGGATGGGCTTGGGCCTGGG | chr9:113536716-113536735 | absent | 5998 | Promoter |
| GGGCAGAGGGTGGGGCAGGG | chr9:121763670-121763689 | absent | 153090 | Promoter |
| GGGGTTGGGTGGGGGCTGGG | chr9:124506593-124506612 | present | 2516 | Promoter |
| GGGCAGGGGACGGGGGTGGG | chr9:27312142-27312161 | absent | 54586 | Intergenic |
| GGGCTGGGGTCGGGGTGGGG | chr9:84670271-84670290 | present | 4915 | Promoter |
| GGGATGGGGAGGGAACAGGG | chrX:18891734-18891753 | present | 100132163 | Promoter |
| GGGCTGGGGCAGGGATAGGG | chrX:19081377-19081396 | absent | 10149 | Promoter |

**Supplemental Table 8. Summary of Family 75 G4 sequences.**

| Sequence | Location | Experimental Evidence | Gene ID | Annotation |
|---|---|---|---|---|
| GGGGGAGGGAGGGCCTGGG | chr11:19817544-19817562 | present | 89797 | Intron |
| GGGGGTGGGAGGGGCAGGG | chr11:65967601-65967619 | present | 9092 | Intron |
| GGGTGGAGGGAGGGCTGGG | chr12:130663161-130663179 | absent | 23504 | Intron |
| GGGGGTGGGGGGGCCTGGG | chr12:56096956-56096974 | present | 2065 | Promoter |
| GGGGGTGGGAGGGCAGGG | chr16:78000971-78000988 | present | 10143 | Intergenic |
| GGGGTGGGAGGGGCATGGG | chr17:41888505-41888523 | present | 47 | Intron |
| GGGGGTGGGAGGGCATGGG | chr19:52643765-52643783 | present | 55769 | Intron |
| GGGGGTGGGAGGGCATGGG | chr19:52700738-52700756 | present | 55769 | Downstream |
| GGGGGTGGGAGGGCATGGG | chr19:52762022-52762040 | present | 162966 | Downstream |
| GGGGGTGGGAGGGCATGGG | chr19:52818986-52819004 | present | 7576 | Promoter |
| GGGGGTGGGAGGGCATGGG | chr19:52855275-52855293 | present | 7576 | Promoter |
| GGGGGTGGGAGGGCATGGG | chr19:52938557-52938575 | present | 388559 | Intron |
| GGGGGTGGGAGGGCACGGG | chr19:53129043-53129061 | present | 55786 | Promoter |
| GGGTGGTGGGAGGGATGGG | chr1:18332823-18332841 | absent | 84966 | Intron |
| GGGGGTGGGAGGGCCTGGG | chr20:63293908-63293926 | present | 57642 | Promoter |
| GGGGGTGGGTAGGGCCGGG | chr2:219060082-219060100 | present | 3549 | Promoter |
| GGGGAGAGGGAGGGCCGGG | chr4:40244011-40244029 | absent | 399 | 3' UTR |
| GGGGGAGGGAGGGCTTGGG | chr8:124855332-124855350 | present | 157381 | Intron |

**Supplemental Table 9. Summary of Family 80 G4 sequences.**

| Sequence | Location | Experimental Evidence | Gene ID | Distance To TSS | annot |
|---|---|---|---|---|---|
| GGGGCGGGCTGGGGGCGGGG | chr11:67630369-67630388 | present | 254552 | -439 | Promoter |
| GGGTCGGGGCCGGGGGAGGG | chr11:968778-968797 | present | 161 | -16587 | Intron |
| GGGGCGGGCCTCGGGGGCGGGG | chr14:93184925-93184946 | present | 64112 | 0 | Promoter |
| GGGTGCGGGGCCGGGGGGAGGGG | chr17:39927095-39927117 | absent | 94103 | 112 | Promoter |
| GGGGCTGGGGGCGGGGGCGGG | chr17:79836273-79836293 | present | 8535 | 1588 | Promoter |
| GGGGCGGGGCCGGGGGCGGG | chr19:18097748-18097767 | present | 23031 | -26 | Promoter |
| GGGGCGGGTCGTGGGCGGGG | chr19:2096722-2096741 | present | 126308 | -49 | Promoter |
| GGGGCTGGGTCGGGGGCGGGG | chr19:55081615-55081635 | present | 54869 | -57 | Promoter |
| GGGGCGGGCCCAGGGGCGGG | chr1:16700286-16700305 | present | 100500876 | 19031 | Intergenic |
| GGGGCGGGCCGGGGGGAGGGG | chr1:185411921-185411941 | present | 100288079 | -76882 | Intergenic |
| GGGGCGGGCCGGGGGCGGGG | chr20:5001518-5001537 | present | 9962 | -19 | Promoter |
| GGGGGCGGGCTCGGGGGCGGGG | chr21:5022922-5022943 | present | 23308 | 389 | Promoter |
| GGGGCGGGGCACGGGGGAGGG | chr4:1011532-1011552 | present | 53834 | -270 | Promoter |
| GGGGCGGGACCGGGGAGAGGGG | chr6:11043967-11043988 | present | 100506409 | 207 | Promoter |
| GGGGGGGGTAGTGGGCGGGG | chr7:156169292-156169311 | present | 389602 | 206660 | Intergenic |
| GGGGCGGGCCGTGGGCCGGG | chr7:158829641-158829660 | absent | 57488 | -13 | Promoter |
| GGGGCAGGCCGGGCGGGAGGGG | chr7:20798511-20798532 | absent | 221833 | -11625 | Intergenic |
| GGGGCGGGGGCCGCGGGCGGG | chr7:6374902-6374922 | absent | 5879 | 339 | Promoter |
| GGGGCCGGGGCCGGGGCCGGG | chr8:22049068-22049088 | absent | 2039 | -61 | Promoter |
| GGGGCGGGCTCGGGGGCGGGG | chr8:66962618-66962638 | present | 100129654 | -28 | Promoter |
| GGGGCCGGGCCGAGGGGCGGG | chr9:132241390-132241410 | present | 84628 | -31 | Promoter |

**Supplemental Table 10. Enriched GO:BP categories for Family 4.**

| GO Term | GO Term Name | P Value | ADJ P Value |
|---|---|---|---|
| GO:0021815 | modulation of microtubule cytoskeleton involved in cerebral cortex radial glia guided migration | 2.13E-05 | 2.13E-05 |
| GO:0021816 | extension of a leading process involved in cell motility in cerebral cortex radial glia guided migration | 2.13E-05 | 2.13E-05 |
| GO:0021814 | cell motility involved in cerebral cortex radial glia guided migration | 8.51E-05 | 8.51E-05 |
| GO:0022030 | telencephalon glial cell migration | 0.000382 | 0.000382 |
| GO:0021801 | cerebral cortex radial glia-guided migration | 0.000382 | 0.000382 |
| GO:0021799 | cerebral cortex radially oriented cell migration | 0.000389 | 0.000389 |
| GO:0031269 | pseudopodium assembly | 0.000552 | 0.000552 |
| GO:0031268 | pseudopodium organization | 0.000557 | 0.000557 |
| GO:0021795 | cerebral cortex cell migration | 0.00059 | 0.00059 |
| GO:1904861 | excitatory synapse assembly | 0.00059 | 0.00059 |
| GO:1904862 | inhibitory synapse assembly | 0.00059 | 0.00059 |
| GO:0022029 | telencephalon cell migration | 0.000893 | 0.000893 |
| GO:0021885 | forebrain cell migration | 0.000899 | 0.000899 |
| GO:0008347 | glial cell migration | 0.000983 | 0.000983 |
| GO:2001222 | regulation of neuron migration | 0.001145 | 0.001145 |
| GO:0021987 | cerebral cortex development | 0.003356 | 0.003356 |
| GO:0046847 | filopodium assembly | 0.004232 | 0.004232 |
| GO:0060996 | dendritic spine development | 0.004705 | 0.004705 |
| GO:0021543 | pallium development | 0.00474 | 0.00474 |
| GO:0001764 | neuron migration | 0.007448 | 0.007448 |
| GO:0021537 | telencephalon development | 0.009569 | 0.009569 |
| GO:0007416 | synapse assembly | 0.0142 | 0.0142 |
| GO:0030900 | forebrain development | 0.017572 | 0.017572 |
| GO:0016358 | dendrite development | 0.017572 | 0.017572 |
| GO:0042063 | gliogenesis | 0.017572 | 0.017572 |
| GO:0030336 | negative regulation of cell migration | 0.031913 | 0.031913 |
| GO:2000146 | negative regulation of cell motility | 0.034153 | 0.034153 |
| GO:0050808 | synapse organization | 0.035988 | 0.035988 |
| GO:0040013 | negative regulation of locomotion | 0.035988 | 0.035988 |
| GO:0034329 | cell junction assembly | 0.046932 | 0.046932 |

**Supplemental Table 11. Enriched GO:BP categories for Family 32.**

| GO ID | GO Name | P Value | ADJ P Value |
|---|---|---|---|
| GO:0031346 | positive regulation of cell projection organization | 0.017646 | 0.017646 |
| GO:0035378 | carbon dioxide transmembrane transport | 0.017646 | 0.017646 |
| GO:0032989 | cellular component morphogenesis | 0.023881 | 0.023881 |
| GO:0048842 | positive regulation of axon extension involved in axon guidance | 0.023881 | 0.023881 |
| GO:0048858 | cell projection morphogenesis | 0.023881 | 0.023881 |
| GO:0003097 | renal water transport | 0.023881 | 0.023881 |
| GO:0051130 | positive regulation of cellular component organization | 0.023881 | 0.023881 |
| GO:0051239 | regulation of multicellular organismal process | 0.023881 | 0.023881 |
| GO:0048846 | axon extension involved in axon guidance | 0.023881 | 0.023881 |
| GO:0120039 | plasma membrane bounded cell projection morphogenesis | 0.023881 | 0.023881 |
| GO:1902284 | neuron projection extension involved in neuron projection guidance | 0.023881 | 0.023881 |
| GO:1903955 | positive regulation of protein targeting to mitochondrion | 0.023881 | 0.023881 |
| GO:0032990 | cell part morphogenesis | 0.024503 | 0.024503 |
| GO:1903749 | positive regulation of establishment of protein localization to mitochondrion | 0.025365 | 0.025365 |
| GO:0097485 | neuron projection guidance | 0.029584 | 0.029584 |
| GO:0007411 | axon guidance | 0.029584 | 0.029584 |
| GO:1903214 | regulation of protein targeting to mitochondrion | 0.032099 | 0.032099 |
| GO:0007167 | enzyme-linked receptor protein signaling pathway | 0.032099 | 0.032099 |
| GO:0048518 | positive regulation of biological process | 0.032578 | 0.032578 |
| GO:0051094 | positive regulation of developmental process | 0.033021 | 0.033021 |
| GO:0050772 | positive regulation of axonogenesis | 0.03945 | 0.03945 |
| GO:0048468 | cell development | 0.03945 | 0.03945 |
| GO:1903747 | regulation of establishment of protein localization to mitochondrion | 0.03945 | 0.03945 |
| GO:0007409 | axonogenesis | 0.03945 | 0.03945 |
| GO:0022603 | regulation of anatomical structure morphogenesis | 0.039943 | 0.039943 |
| GO:0048812 | neuron projection morphogenesis | 0.045216 | 0.045216 |
| GO:0000902 | cell morphogenesis | 0.045216 | 0.045216 |
| GO:0008361 | regulation of cell size | 0.045216 | 0.045216 |
| GO:0051347 | positive regulation of transferase activity | 0.045887 | 0.045887 |
| GO:0061564 | axon development | 0.045887 | 0.045887 |
| GO:0090066 | regulation of anatomical structure size | 0.046587 | 0.046587 |
| GO:0120035 | regulation of plasma membrane bounded cell projection organization | 0.046587 | 0.046587 |

**Supplemental Table 12. Enriched GO:BP categories for Family 75.**

| GO ID | GO Name | P Value | ADJ P Value |
|---|---|---|---|
| GO:0045582 | positive regulation of T cell differentiation | 0.015898 | 0.015898 |
| GO:0045621 | positive regulation of lymphocyte differentiation | 0.015898 | 0.015898 |
| GO:1903708 | positive regulation of hemopoiesis | 0.016146 | 0.016146 |
| GO:1902107 | positive regulation of leukocyte differentiation | 0.016146 | 0.016146 |
| GO:0045580 | regulation of T cell differentiation | 0.016146 | 0.016146 |
| GO:0045619 | regulation of lymphocyte differentiation | 0.019728 | 0.019728 |
| GO:0050870 | positive regulation of T cell activation | 0.03222 | 0.03222 |
| GO:0030217 | T cell differentiation | 0.03222 | 0.03222 |
| GO:1903039 | positive regulation of leukocyte cell-cell adhesion | 0.03674 | 0.03674 |
| GO:1902105 | regulation of leukocyte differentiation | 0.03674 | 0.03674 |
| GO:0022409 | positive regulation of cell-cell adhesion | 0.047638 | 0.047638 |
| GO:1903037 | regulation of leukocyte cell-cell adhesion | 0.04765 | 0.04765 |
| GO:0030155 | regulation of cell adhesion | 0.04765 | 0.04765 |
| GO:0030098 | lymphocyte differentiation | 0.04765 | 0.04765 |
| GO:0050863 | regulation of T cell activation | 0.04765 | 0.04765 |
| GO:1903706 | regulation of hemopoiesis | 0.049277 | 0.049277 |

**Supplemental Table 13. Enriched GO:BP categories for Family 80.**

| GO ID | GO Name | P Value | ADJ P Value |
|---|---|---|---|
| GO:0098562 | cytoplasmic side of membrane | 1.57E-02 | 1.57E-02 |
| GO:0005886 | plasma membrane | 1.57E-02 | 1.57E-02 |
| GO:0009898 | cytoplasmic side of plasma membrane | 1.57E-02 | 1.57E-02 |
| GO:0098590 | plasma membrane region | 1.57E-02 | 1.57E-02 |
| GO:0031253 | cell projection membrane | 1.81E-02 | 1.81E-02 |
| GO:0101003 | ficolin-1-rich granule membrane | 1.86E-02 | 1.86E-02 |
| GO:0098552 | side of membrane | 1.86E-02 | 1.86E-02 |
| GO:0071944 | cell periphery | 1.86E-02 | 1.86E-02 |
| GO:0032587 | ruffle membrane | 1.86E-02 | 1.86E-02 |
| GO:0030667 | secretory granule membrane | 1.93E-02 | 1.93E-02 |
| GO:0031256 | leading edge membrane | 3.64E-02 | 3.64E-02 |
| GO:0005884 | actin filament | 3.64E-02 | 3.64E-02 |
| GO:0031234 | extrinsic component of cytoplasmic side of plasma membrane | 3.64E-02 | 3.64E-02 |
| GO:0001726 | ruffle | 3.85E-02 | 3.85E-02 |
| GO:0016020 | membrane | 3.85E-02 | 3.85E-02 |
| GO:0019897 | extrinsic component of plasma membrane | 4.29E-02 | 4.29E-02 |
| GO:0031224 | intrinsic component of membrane | 4.29E-02 | 4.29E-02 |
| GO:0031227 | intrinsic component of endoplasmic reticulum membrane | 4.29E-02 | 4.29E-02 |
| GO:0098797 | plasma membrane protein complex | 4.29E-02 | 4.29E-02 |
| GO:0101002 | ficolin-1-rich granule | 4.29E-02 | 4.29E-02 |
| GO:0070820 | tertiary granule | 4.29E-02 | 4.29E-02 |

**Supplemental Table 14. Enriched GO:BP categories for experimentally validated G4s overlapping enhancers, group 1.**

| GO ID | GO Name | P Value | ADJ P Value |
|---|---|---|---|
| GO:0048583 | regulation of response to stimulus | 2.28E-06 | 2.28E-06 |
| GO:0035556 | intracellular signal transduction | 2.28E-06 | 2.28E-06 |
| GO:0010033 | response to organic substance | 2.36E-06 | 2.36E-06 |
| GO:0007165 | signal transduction | 3.86E-06 | 3.86E-06 |
| GO:1902531 | regulation of intracellular signal transduction | 3.86E-06 | 3.86E-06 |
| GO:0009966 | regulation of signal transduction | 3.86E-06 | 3.86E-06 |
| GO:0050896 | response to stimulus | 6.92E-06 | 6.92E-06 |
| GO:0048584 | positive regulation of response to stimulus | 6.92E-06 | 6.92E-06 |
| GO:0007166 | cell surface receptor signaling pathway | 2.89E-05 | 2.89E-05 |
| GO:0007154 | cell communication | 2.89E-05 | 2.89E-05 |
| GO:0010646 | regulation of cell communication | 6.71E-05 | 6.71E-05 |
| GO:1902533 | positive regulation of intracellular signal transduction | 6.71E-05 | 6.71E-05 |
| GO:0023051 | regulation of signaling | 6.71E-05 | 6.71E-05 |
| GO:0023052 | signaling | 6.71E-05 | 6.71E-05 |
| GO:0034097 | response to cytokine | 6.96E-05 | 6.96E-05 |
| GO:0051716 | cellular response to stimulus | 7.39E-05 | 7.39E-05 |
| GO:0009967 | positive regulation of signal transduction | 7.42E-05 | 7.42E-05 |
| GO:0009615 | response to virus | 0.000118423 | 0.0001184 |
| GO:0010647 | positive regulation of cell communication | 0.000218782 | 0.0002188 |
| GO:0032101 | regulation of response to external stimulus | 0.000228634 | 0.0002286 |
| GO:0023056 | positive regulation of signaling | 0.000228634 | 0.0002286 |
| GO:0071310 | cellular response to organic substance | 0.000234694 | 0.0002347 |
| GO:0019221 | cytokine-mediated signaling pathway | 0.000236278 | 0.0002363 |
| GO:0042221 | response to chemical | 0.000236278 | 0.0002363 |
| GO:0012501 | programmed cell death | 0.000236278 | 0.0002363 |
| GO:0006915 | apoptotic process | 0.000236919 | 0.0002369 |
| GO:0009605 | response to external stimulus | 0.000237548 | 0.0002375 |
| GO:0044419 | biological process involved in interspecies interaction between organisms | 0.000270631 | 0.0002706 |
| GO:0097190 | apoptotic signaling pathway | 0.000378032 | 0.000378 |
| GO:0070887 | cellular response to chemical stimulus | 0.000398413 | 0.0003984 |
| GO:0051607 | defense response to virus | 0.000398413 | 0.0003984 |
| GO:0140546 | defense response to symbiont | 0.000398413 | 0.0003984 |
| GO:0007249 | I-kappaB kinase/NF-kappaB signaling | 0.000398413 | 0.0003984 |
| GO:1904747 | positive regulation of apoptotic process involved in development | 0.000693039 | 0.000693 |
| GO:1902339 | positive regulation of apoptotic process involved in morphogenesis | 0.000693039 | 0.000693 |
| GO:0009753 | response to jasmonic acid | 0.000693039 | 0.000693 |
| GO:0071395 | cellular response to jasmonic acid stimulus | 0.000693039 | 0.000693 |
| GO:0031347 | regulation of defense response | 0.000693039 | 0.000693 |
| GO:0071345 | cellular response to cytokine stimulus | 0.000694499 | 0.0006945 |
| GO:0048518 | positive regulation of biological process | 0.001032031 | 0.001032 |
| GO:0051055 | negative regulation of lipid biosynthetic process | 0.001298251 | 0.0012983 |
| GO:0008219 | cell death | 0.00131134 | 0.0013113 |
| GO:0080134 | regulation of response to stress | 0.00131134 | 0.0013113 |
| GO:0070542 | response to fatty acid | 0.001354706 | 0.0013547 |
| GO:1901798 | positive regulation of signal transduction by p53 class mediator | 0.001354706 | 0.0013547 |
| GO:0016032 | viral process | 0.001418243 | 0.0014182 |
| GO:0002376 | immune system process | 0.001550915 | 0.0015509 |
| GO:0043122 | regulation of I-kappaB kinase/NF-kappaB signaling | 0.001991596 | 0.0019916 |
| GO:0043207 | response to external biotic stimulus | 0.002342802 | 0.0023428 |
| GO:0051707 | response to other organism | 0.002342802 | 0.0023428 |
| GO:0038061 | NIK/NF-kappaB signaling | 0.002342802 | 0.0023428 |
| GO:0043067 | regulation of programmed cell death | 0.003172007 | 0.003172 |
| GO:1903829 | positive regulation of protein localization | 0.003172007 | 0.003172 |
| GO:0008630 | intrinsic apoptotic signaling pathway in response to DNA damage | 0.003495663 | 0.0034957 |
| GO:1901222 | regulation of NIK/NF-kappaB signaling | 0.003495663 | 0.0034957 |
| GO:0045071 | negative regulation of viral genome replication | 0.003495663 | 0.0034957 |
| GO:0033209 | tumor necrosis factor-mediated signaling pathway | 0.003654884 | 0.0036549 |
| GO:0071398 | cellular response to fatty acid | 0.003654884 | 0.0036549 |
| GO:1902337 | regulation of apoptotic process involved in morphogenesis | 0.003677392 | 0.0036774 |
| GO:0009607 | response to biotic stimulus | 0.003677392 | 0.0036774 |
| GO:1904748 | regulation of apoptotic process involved in development | 0.003677392 | 0.0036774 |
| GO:0045833 | negative regulation of lipid metabolic process | 0.003695766 | 0.0036958 |
| GO:0016601 | Rac protein signal transduction | 0.004136778 | 0.0041368 |
| GO:0072331 | signal transduction by p53 class mediator | 0.004424419 | 0.0044244 |
| GO:0042981 | regulation of apoptotic process | 0.004424419 | 0.0044244 |
| GO:0006952 | defense response | 0.004489547 | 0.0044895 |

**Supplemental Table 14. (Continued).**

| GO ID | GO Name | P Value | ADJ P Value |
|---|---|---|---|
| GO:0048522 | positive regulation of cellular process | 0.004536959 | 0.004537 |
| GO:0071677 | positive regulation of mononuclear cell migration | 0.004728666 | 0.0047287 |
| GO:0050789 | regulation of biological process | 0.005144355 | 0.0051444 |
| GO:0048525 | negative regulation of viral process | 0.005144355 | 0.0051444 |
| GO:0032502 | developmental process | 0.005423495 | 0.0054235 |
| GO:0050793 | regulation of developmental process | 0.005888178 | 0.0058882 |
| GO:0008625 | extrinsic apoptotic signaling pathway via death domain receptors | 0.00633763 | 0.0063376 |
| GO:1902644 | tertiary alcohol metabolic process | 0.006593339 | 0.0065933 |
| GO:0030647 | aminoglycoside antibiotic metabolic process | 0.006614403 | 0.0066144 |
| GO:0070383 | DNA cytosine deamination | 0.006614403 | 0.0066144 |
| GO:0044597 | daunorubicin metabolic process | 0.006614403 | 0.0066144 |
| GO:0097193 | intrinsic apoptotic signaling pathway | 0.006614403 | 0.0066144 |
| GO:0060561 | apoptotic process involved in morphogenesis | 0.006614403 | 0.0066144 |
| GO:0070278 | extracellular matrix constituent secretion | 0.006614403 | 0.0066144 |
| GO:0032103 | positive regulation of response to external stimulus | 0.006708023 | 0.006708 |
| GO:0071356 | cellular response to tumor necrosis factor | 0.00737156 | 0.0073716 |
| GO:0043065 | positive regulation of apoptotic process | 0.007883042 | 0.007883 |
| GO:0046890 | regulation of lipid biosynthetic process | 0.007883042 | 0.007883 |
| GO:0031327 | negative regulation of cellular biosynthetic process | 0.008116453 | 0.0081165 |
| GO:0030638 | polyketide metabolic process | 0.008227763 | 0.0082278 |
| GO:0010648 | negative regulation of cell communication | 0.008227763 | 0.0082278 |
| GO:0044598 | doxorubicin metabolic process | 0.008227763 | 0.0082278 |
| GO:0016554 | cytidine to uridine editing | 0.008227763 | 0.0082278 |
| GO:0006950 | response to stress | 0.008227763 | 0.0082278 |
| GO:0032880 | regulation of protein localization | 0.008227763 | 0.0082278 |
| GO:0048856 | anatomical structure development | 0.008355755 | 0.0083558 |
| GO:0097191 | extrinsic apoptotic signaling pathway | 0.008544215 | 0.0085442 |
| GO:0044249 | cellular biosynthetic process | 0.008552028 | 0.008552 |
| GO:0023057 | negative regulation of signaling | 0.008552028 | 0.008552 |
| GO:0006954 | inflammatory response | 0.009069224 | 0.0090692 |
| GO:0009890 | negative regulation of biosynthetic process | 0.009624718 | 0.0096247 |
| GO:0043068 | positive regulation of programmed cell death | 0.009624718 | 0.0096247 |
| GO:0002831 | regulation of response to biotic stimulus | 0.00989629 | 0.0098963 |
| GO:0048523 | negative regulation of cellular process | 0.010235836 | 0.0102358 |
| GO:0009893 | positive regulation of metabolic process | 0.010235836 | 0.0102358 |
| GO:0030865 | cortical cytoskeleton organization | 0.010319334 | 0.0103193 |
| GO:0002682 | regulation of immune system process | 0.010684546 | 0.0106845 |
| GO:0010941 | regulation of cell death | 0.0108774 | 0.0108774 |
| GO:1901576 | organic substance biosynthetic process | 0.010926992 | 0.010927 |
| GO:0051896 | regulation of protein kinase B signaling | 0.010926992 | 0.010927 |
| GO:0034612 | response to tumor necrosis factor | 0.010946178 | 0.0109462 |
| GO:0032102 | negative regulation of response to external stimulus | 0.011229625 | 0.0112296 |
| GO:0002468 | dendritic cell antigen processing and presentation | 0.012370175 | 0.0123702 |
| GO:0019216 | regulation of lipid metabolic process | 0.013075598 | 0.0130756 |
| GO:0048585 | negative regulation of response to stimulus | 0.013075598 | 0.0130756 |
| GO:0045892 | negative regulation of DNA-templated transcription | 0.013571874 | 0.0135719 |
| GO:0002484 | antigen processing and presentation of endogenous peptide antigen via MHC class I via ER pathway | 0.01392641 | 0.0139264 |
| GO:0002486 | antigen processing and presentation of endogenous peptide antigen via MHC class I via ER pathway, TAP-independent | 0.01392641 | 0.0139264 |
| GO:0071798 | response to prostaglandin D | 0.01392641 | 0.0139264 |
| GO:1902679 | negative regulation of RNA biosynthetic process | 0.01392641 | 0.0139264 |
| GO:1903507 | negative regulation of nucleic acid-templated transcription | 0.01392641 | 0.0139264 |
| GO:0048869 | cellular developmental process | 0.01392641 | 0.0139264 |
| GO:0071799 | cellular response to prostaglandin D stimulus | 0.01392641 | 0.0139264 |
| GO:0045006 | DNA deamination | 0.01401026 | 0.0140103 |
| GO:2000010 | positive regulation of protein localization to cell surface | 0.01401026 | 0.0140103 |
| GO:1900025 | negative regulation of substrate adhesion-dependent cell spreading | 0.01401026 | 0.0140103 |
| GO:0042448 | progesterone metabolic process | 0.01401026 | 0.0140103 |
| GO:0010771 | negative regulation of cell morphogenesis involved in differentiation | 0.01401026 | 0.0140103 |
| GO:0032879 | regulation of localization | 0.014084941 | 0.0140849 |
| GO:0009058 | biosynthetic process | 0.014084941 | 0.0140849 |
| GO:0045069 | regulation of viral genome replication | 0.014214363 | 0.0142144 |
| GO:1901700 | response to oxygen-containing compound | 0.014264573 | 0.0142646 |
| GO:0008207 | C21-steroid hormone metabolic process | 0.014714282 | 0.0147143 |
| GO:0009649 | entrainment of circadian clock | 0.014714282 | 0.0147143 |

**Supplemental Table 14. (Continued).**

| GO ID | GO Name | P Value | ADJ P Value |
|---|---|---|---|
| GO:0031341 | regulation of cell killing | 0.014714282 | 0.0147143 |
| GO:0045732 | positive regulation of protein catabolic process | 0.014783883 | 0.0147839 |
| GO:0046649 | lymphocyte activation | 0.014783883 | 0.0147839 |
| GO:0019079 | viral genome replication | 0.014854681 | 0.0148547 |
| GO:0050792 | regulation of viral process | 0.015553577 | 0.0155536 |
| GO:0065007 | biological regulation | 0.015880386 | 0.0158804 |
| GO:0002687 | positive regulation of leukocyte migration | 0.016099492 | 0.0160995 |
| GO:0043542 | endothelial cell migration | 0.016558791 | 0.0165588 |
| GO:0048519 | negative regulation of biological process | 0.01755302 | 0.017553 |
| GO:0031349 | positive regulation of defense response | 0.017642697 | 0.0176427 |
| GO:0019058 | viral life cycle | 0.017642697 | 0.0176427 |
| GO:0030154 | cell differentiation | 0.017814671 | 0.0178147 |
| GO:0043491 | protein kinase B signaling | 0.018152988 | 0.018153 |
| GO:0045869 | negative regulation of single stranded viral RNA replication via double stranded DNA intermediate | 0.018723896 | 0.0187239 |
| GO:0034694 | response to prostaglandin | 0.018723896 | 0.0187239 |
| GO:0030036 | actin cytoskeleton organization | 0.019390353 | 0.0193904 |
| GO:0051253 | negative regulation of RNA metabolic process | 0.019530455 | 0.0195305 |
| GO:1904377 | positive regulation of protein localization to cell periphery | 0.019927046 | 0.019927 |
| GO:0001775 | cell activation | 0.019927046 | 0.019927 |
| GO:0050688 | regulation of defense response to virus | 0.019927046 | 0.019927 |
| GO:0019882 | antigen processing and presentation | 0.020806217 | 0.0208062 |
| GO:0031326 | regulation of cellular biosynthetic process | 0.020872326 | 0.0208723 |
| GO:0061044 | negative regulation of vascular wound healing | 0.021001067 | 0.0210011 |
| GO:0071072 | negative regulation of phospholipid biosynthetic process | 0.021001067 | 0.0210011 |
| GO:0003332 | negative regulation of extracellular matrix constituent secretion | 0.021001067 | 0.0210011 |
| GO:0010604 | positive regulation of macromolecule metabolic process | 0.021001067 | 0.0210011 |
| GO:0010942 | positive regulation of cell death | 0.021646385 | 0.0216464 |
| GO:1900744 | regulation of p38MAPK cascade | 0.024190946 | 0.0241909 |
| GO:0060341 | regulation of cellular localization | 0.024190946 | 0.0241909 |
| GO:0016137 | glycoside metabolic process | 0.024636699 | 0.0246367 |
| GO:0006955 | immune response | 0.024636699 | 0.0246367 |
| GO:0051173 | positive regulation of nitrogen compound metabolic process | 0.024636699 | 0.0246367 |
| GO:0051701 | biological process involved in interaction with host | 0.025116865 | 0.0251169 |
| GO:0050691 | regulation of defense response to virus by host | 0.025813005 | 0.025813 |
| GO:0030029 | actin filament-based process | 0.025944795 | 0.0259448 |
| GO:0009889 | regulation of biosynthetic process | 0.025944795 | 0.0259448 |
| GO:0010558 | negative regulation of macromolecule biosynthetic process | 0.025944795 | 0.0259448 |
| GO:0051897 | positive regulation of protein kinase B signaling | 0.025963166 | 0.0259632 |
| GO:0045595 | regulation of cell differentiation | 0.025963166 | 0.0259632 |
| GO:0009968 | negative regulation of signal transduction | 0.026066467 | 0.0260665 |
| GO:1903900 | regulation of viral life cycle | 0.026153803 | 0.0261538 |
| GO:0030155 | regulation of cell adhesion | 0.027008361 | 0.0270084 |
| GO:0044403 | biological process involved in symbiotic interaction | 0.027008361 | 0.0270084 |
| GO:0043652 | engulfment of apoptotic cell | 0.027008361 | 0.0270084 |
| GO:0016553 | base conversion or substitution editing | 0.027008361 | 0.0270084 |
| GO:0033036 | macromolecule localization | 0.027008361 | 0.0270084 |
| GO:0002685 | regulation of leukocyte migration | 0.027008361 | 0.0270084 |
| GO:0002684 | positive regulation of immune system process | 0.027008361 | 0.0270084 |
| GO:0002483 | antigen processing and presentation of endogenous peptide antigen | 0.027008361 | 0.0270084 |
| GO:0033993 | response to lipid | 0.027008361 | 0.0270084 |
| GO:0043123 | positive regulation of I-kappaB kinase/NF-kappaB signaling | 0.027008361 | 0.0270084 |
| GO:0050794 | regulation of cellular process | 0.027008361 | 0.0270084 |
| GO:0009719 | response to endogenous stimulus | 0.027008361 | 0.0270084 |
| GO:0045091 | regulation of single stranded viral RNA replication via double stranded DNA intermediate | 0.027008361 | 0.0270084 |
| GO:0071222 | cellular response to lipopolysaccharide | 0.027268538 | 0.0272685 |
| GO:0019222 | regulation of metabolic process | 0.027268538 | 0.0272685 |
| GO:0043277 | apoptotic cell clearance | 0.027268538 | 0.0272685 |
| GO:0051247 | positive regulation of protein metabolic process | 0.027268538 | 0.0272685 |
| GO:0008610 | lipid biosynthetic process | 0.027268538 | 0.0272685 |
| GO:0071675 | regulation of mononuclear cell migration | 0.028066202 | 0.0280662 |
| GO:0051093 | negative regulation of developmental process | 0.028066202 | 0.0280662 |
| GO:1903726 | negative regulation of phospholipid metabolic process | 0.028066202 | 0.0280662 |
| GO:1904238 | pericyte cell differentiation | 0.028066202 | 0.0280662 |
| GO:0001910 | regulation of leukocyte mediated cytotoxicity | 0.028579395 | 0.0285794 |
| GO:0019218 | regulation of steroid metabolic process | 0.028579395 | 0.0285794 |

**Supplemental Table 14. (Continued).**

| GO ID | GO Name | P Value | ADJ P Value |
|---|---|---|---|
| GO:0045934 | negative regulation of nucleobase-containing compound metabolic process | 0.028579395 | 0.0285794 |
| GO:0039692 | single stranded viral RNA replication via double stranded DNA intermediate | 0.028579395 | 0.0285794 |
| GO:0031324 | negative regulation of cellular metabolic process | 0.029203388 | 0.0292034 |
| GO:0002697 | regulation of immune effector process | 0.030651865 | 0.0306519 |
| GO:0045862 | positive regulation of proteolysis | 0.030651865 | 0.0306519 |
| GO:0051234 | establishment of localization | 0.031842009 | 0.031842 |
| GO:0019748 | secondary metabolic process | 0.033101255 | 0.0331013 |
| GO:2000059 | negative regulation of ubiquitin-dependent protein catabolic process | 0.033101255 | 0.0331013 |
| GO:2000630 | positive regulation of miRNA metabolic process | 0.033101255 | 0.0331013 |
| GO:0002819 | regulation of adaptive immune response | 0.033137999 | 0.033138 |
| GO:0045321 | leukocyte activation | 0.033186568 | 0.0331866 |
| GO:1904375 | regulation of protein localization to cell periphery | 0.033464279 | 0.0334643 |
| GO:0071496 | cellular response to external stimulus | 0.033909758 | 0.0339098 |
| GO:0045444 | fat cell differentiation | 0.033909758 | 0.0339098 |
| GO:1905475 | regulation of protein localization to membrane | 0.034848048 | 0.034848 |
| GO:0071219 | cellular response to molecule of bacterial origin | 0.034848048 | 0.034848 |
| GO:1902742 | apoptotic process involved in development | 0.0351558 | 0.0351558 |
| GO:1901701 | cellular response to oxygen-containing compound | 0.0351558 | 0.0351558 |
| GO:0098542 | defense response to other organism | 0.0351558 | 0.0351558 |
| GO:2001141 | regulation of RNA biosynthetic process | 0.035397391 | 0.0353974 |
| GO:0030178 | negative regulation of Wnt signaling pathway | 0.035397391 | 0.0353974 |
| GO:0048260 | positive regulation of receptor-mediated endocytosis | 0.036870748 | 0.0368707 |
| GO:1905897 | regulation of response to endoplasmic reticulum stress | 0.037418355 | 0.0374184 |
| GO:0006629 | lipid metabolic process | 0.037771303 | 0.0377713 |
| GO:0031328 | positive regulation of cellular biosynthetic process | 0.038812722 | 0.0388127 |
| GO:0038066 | p38MAPK cascade | 0.038928765 | 0.0389288 |
| GO:0045807 | positive regulation of endocytosis | 0.038928765 | 0.0389288 |
| GO:0051179 | localization | 0.038928765 | 0.0389288 |
| GO:0048513 | animal organ development | 0.039697904 | 0.0396979 |
| GO:0062014 | negative regulation of small molecule metabolic process | 0.040705221 | 0.0407052 |
| GO:0009059 | macromolecule biosynthetic process | 0.040837389 | 0.0408374 |
| GO:0097435 | supramolecular fiber organization | 0.041494668 | 0.0414947 |
| GO:0001667 | ameboidal-type cell migration | 0.041494668 | 0.0414947 |
| GO:0001892 | embryonic placenta development | 0.042976289 | 0.0429763 |
| GO:0022604 | regulation of cell morphogenesis | 0.043574098 | 0.0435741 |
| GO:0030199 | collagen fibril organization | 0.043788574 | 0.0437886 |
| GO:0001912 | positive regulation of leukocyte mediated cytotoxicity | 0.043788574 | 0.0437886 |
| GO:0030162 | regulation of proteolysis | 0.043806228 | 0.0438062 |
| GO:0050729 | positive regulation of inflammatory response | 0.04466043 | 0.0446604 |
| GO:0051785 | positive regulation of nuclear division | 0.046435533 | 0.0464355 |
| GO:0009891 | positive regulation of biosynthetic process | 0.046435533 | 0.0464355 |
| GO:0043433 | negative regulation of DNA-binding transcription factor activity | 0.046827685 | 0.0468277 |
| GO:0031652 | positive regulation of heat generation | 0.046840867 | 0.0468409 |
| GO:0019883 | antigen processing and presentation of endogenous antigen | 0.046840867 | 0.0468409 |
| GO:0010876 | lipid localization | 0.047285215 | 0.0472852 |
| GO:0007586 | digestion | 0.047408175 | 0.0474082 |
| GO:0016477 | cell migration | 0.04744546 | 0.0474455 |
| GO:0051090 | regulation of DNA-binding transcription factor activity | 0.048491772 | 0.0484918 |
| GO:0051246 | regulation of protein metabolic process | 0.048491772 | 0.0484918 |
| GO:0090090 | negative regulation of canonical Wnt signaling pathway | 0.048491772 | 0.0484918 |
| GO:1903506 | regulation of nucleic acid-templated transcription | 0.049386871 | 0.0493869 |

**Supplemental Table 15. Enriched GO:BP categories for experimentally validated G4s overlapping enhancers, group 2.**

| GO ID | GO Name | P Value | ADJ P Value |
|---|---|---|---|
| GO:0002376 | immune system process | 4.15E-23 | 4.15E-23 |
| GO:0006955 | immune response | 5.99E-23 | 5.99E-23 |
| GO:0002682 | regulation of immune system process | 2.97E-21 | 2.97E-21 |
| GO:0002684 | positive regulation of immune system process | 7.09E-18 | 7.09E-18 |
| GO:0050776 | regulation of immune response | 1.08E-15 | 1.08E-15 |
| GO:0002764 | immune response-regulating signaling pathway | 8.25E-15 | 8.25E-15 |
| GO:0050778 | positive regulation of immune response | 1.77E-13 | 1.77E-13 |
| GO:0002429 | immune response-activating cell surface receptor signaling pathway | 1.83E-13 | 1.83E-13 |
| GO:0002757 | immune response-activating signal transduction | 1.83E-13 | 1.83E-13 |
| GO:0002768 | immune response-regulating cell surface receptor signaling pathway | 8.77E-13 | 8.77E-13 |
| GO:0002253 | activation of immune response | 1.39E-12 | 1.39E-12 |
| GO:0046649 | lymphocyte activation | 1.75E-11 | 1.75E-11 |
| GO:0045321 | leukocyte activation | 4.23E-11 | 4.23E-11 |
| GO:0048584 | positive regulation of response to stimulus | 6.13E-11 | 6.13E-11 |
| GO:0007165 | signal transduction | 6.13E-11 | 6.13E-11 |
| GO:0002252 | immune effector process | 5.25E-10 | 5.25E-10 |
| GO:0001819 | positive regulation of cytokine production | 1.28E-09 | 1.28E-09 |
| GO:0001775 | cell activation | 1.28E-09 | 1.28E-09 |
| GO:0002250 | adaptive immune response | 1.30E-09 | 1.30E-09 |
| GO:0006952 | defense response | 3.61E-09 | 3.61E-09 |
| GO:0050851 | antigen receptor-mediated signaling pathway | 4.06E-09 | 4.06E-09 |
| GO:0023052 | signaling | 4.37E-09 | 4.37E-09 |
| GO:0007154 | cell communication | 5.81E-09 | 5.81E-09 |
| GO:0002697 | regulation of immune effector process | 6.28E-09 | 6.28E-09 |
| GO:0050852 | T cell receptor signaling pathway | 9.32E-09 | 9.32E-09 |
| GO:0007166 | cell surface receptor signaling pathway | 9.32E-09 | 9.32E-09 |
| GO:0001817 | regulation of cytokine production | 9.32E-09 | 9.32E-09 |
| GO:0001816 | cytokine production | 1.03E-08 | 1.03E-08 |
| GO:0002700 | regulation of production of molecular mediator of immune response | 2.40E-08 | 2.40E-08 |
| GO:0048583 | regulation of response to stimulus | 2.48E-08 | 2.48E-08 |
| GO:0042110 | T cell activation | 2.73E-08 | 2.73E-08 |
| GO:0032103 | positive regulation of response to external stimulus | 7.15E-08 | 7.15E-08 |
| GO:0050896 | response to stimulus | 1.27E-07 | 1.27E-07 |
| GO:0002440 | production of molecular mediator of immune response | 2.12E-07 | 2.12E-07 |
| GO:0051716 | cellular response to stimulus | 2.31E-07 | 2.31E-07 |
| GO:0002702 | positive regulation of production of molecular mediator of immune response | 2.55E-07 | 2.55E-07 |
| GO:0002699 | positive regulation of immune effector process | 3.32E-07 | 3.32E-07 |
| GO:1903131 | mononuclear cell differentiation | 4.15E-07 | 4.15E-07 |
| GO:0032101 | regulation of response to external stimulus | 4.74E-07 | 4.74E-07 |
| GO:0030098 | lymphocyte differentiation | 4.95E-07 | 4.95E-07 |
| GO:0031347 | regulation of defense response | 7.94E-07 | 7.94E-07 |
| GO:0046631 | alpha-beta T cell activation | 1.63E-06 | 1.63E-06 |
| GO:0002521 | leukocyte differentiation | 1.65E-06 | 1.65E-06 |
| GO:0070663 | regulation of leukocyte proliferation | 2.73E-06 | 2.73E-06 |
| GO:0006954 | inflammatory response | 2.82E-06 | 2.82E-06 |
| GO:0031663 | lipopolysaccharide-mediated signaling pathway | 3.23E-06 | 3.23E-06 |
| GO:0046629 | gamma-delta T cell activation | 3.28E-06 | 3.28E-06 |
| GO:0031349 | positive regulation of defense response | 3.75E-06 | 3.75E-06 |
| GO:0010628 | positive regulation of gene expression | 4.15E-06 | 4.15E-06 |
| GO:1903037 | regulation of leukocyte cell-cell adhesion | 4.15E-06 | 4.15E-06 |
| GO:0043207 | response to external biotic stimulus | 4.62E-06 | 4.62E-06 |
| GO:0051707 | response to other organism | 4.62E-06 | 4.62E-06 |
| GO:0051240 | positive regulation of multicellular organismal process | 4.87E-06 | 4.87E-06 |
| GO:0098542 | defense response to other organism | 5.73E-06 | 5.73E-06 |
| GO:0002831 | regulation of response to biotic stimulus | 5.76E-06 | 5.76E-06 |
| GO:0050670 | regulation of lymphocyte proliferation | 5.76E-06 | 5.76E-06 |
| GO:0019221 | cytokine-mediated signaling pathway | 5.76E-06 | 5.76E-06 |
| GO:0032944 | regulation of mononuclear cell proliferation | 6.38E-06 | 6.38E-06 |
| GO:1903039 | positive regulation of leukocyte cell-cell adhesion | 7.12E-06 | 7.12E-06 |
| GO:0002718 | regulation of cytokine production involved in immune response | 7.48E-06 | 7.48E-06 |
| GO:0002367 | cytokine production involved in immune response | 7.48E-06 | 7.48E-06 |
| GO:0009607 | response to biotic stimulus | 7.59E-06 | 7.59E-06 |
| GO:0032755 | positive regulation of interleukin-6 production | 8.88E-06 | 8.88E-06 |
| GO:0032735 | positive regulation of interleukin-12 production | 1.12E-05 | 1.12E-05 |
| GO:0045785 | positive regulation of cell adhesion | 1.26E-05 | 1.26E-05 |
| GO:0007159 | leukocyte cell-cell adhesion | 1.40E-05 | 1.40E-05 |

**Supplemental Table 15. (Continued).**

| GO ID | GO Name | P Value | ADJ P Value |
|---|---|---|---|
| GO:0000165 | MAPK cascade | 1.41E-05 | 1.41E-05 |
| GO:0032675 | regulation of interleukin-6 production | 1.41E-05 | 1.41E-05 |
| GO:0032637 | interleukin-8 production | 1.41E-05 | 1.41E-05 |
| GO:0032635 | interleukin-6 production | 1.41E-05 | 1.41E-05 |
| GO:0032677 | regulation of interleukin-8 production | 1.41E-05 | 1.41E-05 |
| GO:0070661 | leukocyte proliferation | 1.68E-05 | 1.68E-05 |
| GO:0051249 | regulation of lymphocyte activation | 1.91E-05 | 1.91E-05 |
| GO:0002833 | positive regulation of response to biotic stimulus | 2.18E-05 | 2.18E-05 |
| GO:0032757 | positive regulation of interleukin-8 production | 2.52E-05 | 2.52E-05 |
| GO:0044419 | biological process involved in interspecies interaction between organisms | 2.79E-05 | 2.79E-05 |
| GO:0046651 | lymphocyte proliferation | 3.25E-05 | 3.25E-05 |
| GO:0097530 | granulocyte migration | 3.40E-05 | 3.40E-05 |
| GO:0022409 | positive regulation of cell-cell adhesion | 3.49E-05 | 3.49E-05 |
| GO:0002221 | pattern recognition receptor signaling pathway | 3.55E-05 | 3.55E-05 |
| GO:0032943 | mononuclear cell proliferation | 3.56E-05 | 3.56E-05 |
| GO:0070371 | ERK1 and ERK2 cascade | 3.65E-05 | 3.65E-05 |
| GO:0050900 | leukocyte migration | 4.13E-05 | 4.13E-05 |
| GO:0071345 | cellular response to cytokine stimulus | 4.32E-05 | 4.32E-05 |
| GO:0070374 | positive regulation of ERK1 and ERK2 cascade | 4.36E-05 | 4.36E-05 |
| GO:0002520 | immune system development | 4.94E-05 | 4.94E-05 |
| GO:0030097 | hemopoiesis | 5.42E-05 | 5.42E-05 |
| GO:0009605 | response to external stimulus | 6.53E-05 | 6.53E-05 |
| GO:0043410 | positive regulation of MAPK cascade | 6.53E-05 | 6.53E-05 |
| GO:0002443 | leukocyte mediated immunity | 6.61E-05 | 6.61E-05 |
| GO:0035556 | intracellular signal transduction | 6.61E-05 | 6.61E-05 |
| GO:0050865 | regulation of cell activation | 6.67E-05 | 6.67E-05 |
| GO:0048534 | hematopoietic or lymphoid organ development | 7.00E-05 | 7.00E-05 |
| GO:1990266 | neutrophil migration | 7.20E-05 | 7.20E-05 |
| GO:0022407 | regulation of cell-cell adhesion | 7.20E-05 | 7.20E-05 |
| GO:0002220 | innate immune response activating cell surface receptor signaling pathway | 7.20E-05 | 7.20E-05 |
| GO:0032615 | interleukin-12 production | 7.75E-05 | 7.75E-05 |
| GO:0032655 | regulation of interleukin-12 production | 7.75E-05 | 7.75E-05 |
| GO:0050863 | regulation of T cell activation | 8.28E-05 | 8.28E-05 |
| GO:0002758 | innate immune response-activating signal transduction | 8.30E-05 | 8.30E-05 |
| GO:0032760 | positive regulation of tumor necrosis factor production | 8.79E-05 | 8.79E-05 |
| GO:0002694 | regulation of leukocyte activation | 9.15E-05 | 9.15E-05 |
| GO:0050870 | positive regulation of T cell activation | 9.63E-05 | 9.63E-05 |
| GO:0070372 | regulation of ERK1 and ERK2 cascade | 0.000101 | 0.000101363 |
| GO:0002720 | positive regulation of cytokine production involved in immune response | 0.000114 | 0.000114437 |
| GO:1903557 | positive regulation of tumor necrosis factor superfamily cytokine production | 0.000117 | 0.000116819 |
| GO:0050764 | regulation of phagocytosis | 0.000125 | 0.000125399 |
| GO:0070665 | positive regulation of leukocyte proliferation | 0.00014 | 0.000139526 |
| GO:0043408 | regulation of MAPK cascade | 0.000161 | 0.000161131 |
| GO:0034097 | response to cytokine | 0.000169 | 0.000169097 |
| GO:0002460 | adaptive immune response based on somatic recombination of immune receptors built from immunoglobulin superfamily domains | 0.000176 | 0.000175603 |
| GO:0009966 | regulation of signal transduction | 0.000178 | 0.000177725 |
| GO:0002683 | negative regulation of immune system process | 0.000178 | 0.000178272 |
| GO:0032680 | regulation of tumor necrosis factor production | 0.000188 | 0.000187589 |
| GO:0032640 | tumor necrosis factor production | 0.000188 | 0.000187589 |
| GO:0007249 | I-kappaB kinase/NF-kappaB signaling | 0.000188 | 0.000187589 |
| GO:0042129 | regulation of T cell proliferation | 0.000196 | 0.000195896 |
| GO:1902531 | regulation of intracellular signal transduction | 0.000202 | 0.000201588 |
| GO:0050867 | positive regulation of cell activation | 0.000202 | 0.000201588 |
| GO:0050766 | positive regulation of phagocytosis | 0.000204 | 0.000203772 |
| GO:0002703 | regulation of leukocyte mediated immunity | 0.000204 | 0.000203772 |
| GO:0030155 | regulation of cell adhesion | 0.000206 | 0.00020608 |
| GO:0001818 | negative regulation of cytokine production | 0.000225 | 0.000225163 |
| GO:0051251 | positive regulation of lymphocyte activation | 0.000225 | 0.000225163 |
| GO:0051209 | release of sequestered calcium ion into cytosol | 0.000225 | 0.000225163 |
| GO:0050727 | regulation of inflammatory response | 0.000225 | 0.000225163 |
| GO:1903555 | regulation of tumor necrosis factor superfamily cytokine production | 0.000225 | 0.000225163 |
| GO:0002639 | positive regulation of immunoglobulin production | 0.000225 | 0.000225163 |
| GO:0071706 | tumor necrosis factor superfamily cytokine production | 0.000225 | 0.000225163 |
| GO:0051283 | negative regulation of sequestering of calcium ion | 0.000234 | 0.000233805 |
| GO:0051282 | regulation of sequestering of calcium ion | 0.000249 | 0.000248588 |

**Supplemental Table 15. (Continued).**

| GO ID | GO Name | P Value | ADJ P Value |
|---|---|---|---|
| GO:0051235 | maintenance of location | 0.000289 | 0.000289196 |
| GO:0051651 | maintenance of location in cell | 0.000293 | 0.000293015 |
| GO:0051208 | sequestering of calcium ion | 0.000297 | 0.00029746 |
| GO:0042098 | T cell proliferation | 0.000327 | 0.000326578 |
| GO:0045059 | positive thymic T cell selection | 0.000327 | 0.000326578 |
| GO:0002675 | positive regulation of acute inflammatory response | 0.000331 | 0.000330722 |
| GO:0043122 | regulation of I-kappaB kinase/NF-kappaB signaling | 0.000348 | 0.000347721 |
| GO:0097529 | myeloid leukocyte migration | 0.000414 | 0.000414075 |
| GO:0002224 | toll-like receptor signaling pathway | 0.000415 | 0.000415086 |
| GO:0050671 | positive regulation of lymphocyte proliferation | 0.000415 | 0.000415086 |
| GO:0032602 | chemokine production | 0.000416 | 0.000416113 |
| GO:0032642 | regulation of chemokine production | 0.000416 | 0.000416113 |
| GO:0032946 | positive regulation of mononuclear cell proliferation | 0.000432 | 0.000432255 |
| GO:0097553 | calcium ion transmembrane import into cytosol | 0.000435 | 0.000435396 |
| GO:0048518 | positive regulation of biological process | 0.000435 | 0.000435396 |
| GO:0030217 | T cell differentiation | 0.000462 | 0.000462365 |
| GO:0043405 | regulation of MAP kinase activity | 0.000495 | 0.000495183 |
| GO:0050794 | regulation of cellular process | 0.0005 | 0.000499699 |
| GO:0002637 | regulation of immunoglobulin production | 0.000513 | 0.00051273 |
| GO:0002696 | positive regulation of leukocyte activation | 0.000546 | 0.000546307 |
| GO:0009617 | response to bacterium | 0.000588 | 0.000588485 |
| GO:0032613 | interleukin-10 production | 0.000615 | 0.000614909 |
| GO:0032653 | regulation of interleukin-10 production | 0.000615 | 0.000614909 |
| GO:0010646 | regulation of cell communication | 0.000615 | 0.000614909 |
| GO:0043368 | positive T cell selection | 0.000617 | 0.000617145 |
| GO:0031664 | regulation of lipopolysaccharide-mediated signaling pathway | 0.000617 | 0.000617145 |
| GO:0023051 | regulation of signaling | 0.000634 | 0.000633952 |
| GO:0042102 | positive regulation of T cell proliferation | 0.000641 | 0.000640656 |
| GO:0050729 | positive regulation of inflammatory response | 0.000648 | 0.000648482 |
| GO:0032663 | regulation of interleukin-2 production | 0.000648 | 0.000648482 |
| GO:0070383 | DNA cytosine deamination | 0.000648 | 0.000648482 |
| GO:0032623 | interleukin-2 production | 0.000648 | 0.000648482 |
| GO:0071310 | cellular response to organic substance | 0.000729 | 0.00072903 |
| GO:0038093 | Fc receptor signaling pathway | 0.000782 | 0.000782428 |
| GO:0002449 | lymphocyte mediated immunity | 0.000791 | 0.000790591 |
| GO:0006909 | phagocytosis | 0.000838 | 0.000838027 |
| GO:0045061 | thymic T cell selection | 0.000885 | 0.00088504 |
| GO:0016554 | cytidine to uridine editing | 0.000885 | 0.00088504 |
| GO:0071216 | cellular response to biotic stimulus | 0.000891 | 0.000891177 |
| GO:0045089 | positive regulation of innate immune response | 0.000898 | 0.000898357 |
| GO:0002879 | positive regulation of acute inflammatory response to non-antigenic stimulus | 0.000908 | 0.000907555 |
| GO:0002426 | immunoglobulin production in mucosal tissue | 0.000908 | 0.000907555 |
| GO:2000557 | regulation of immunoglobulin production in mucosal tissue | 0.000908 | 0.000907555 |
| GO:0045087 | innate immune response | 0.000908 | 0.000907555 |
| GO:2000558 | positive regulation of immunoglobulin production in mucosal tissue | 0.000908 | 0.000907555 |
| GO:0002525 | acute inflammatory response to non-antigenic stimulus | 0.000908 | 0.000907555 |
| GO:0033993 | response to lipid | 0.000908 | 0.000907555 |
| GO:0002877 | regulation of acute inflammatory response to non-antigenic stimulus | 0.000908 | 0.000907555 |
| GO:0071674 | mononuclear cell migration | 0.001082 | 0.001081991 |
| GO:0009615 | response to virus | 0.001111 | 0.001111069 |
| GO:0032722 | positive regulation of chemokine production | 0.001127 | 0.00112736 |
| GO:0002685 | regulation of leukocyte migration | 0.001202 | 0.001201974 |
| GO:0140546 | defense response to symbiont | 0.001215 | 0.001215436 |
| GO:0051607 | defense response to virus | 0.001215 | 0.001215436 |
| GO:0071396 | cellular response to lipid | 0.001236 | 0.001236267 |
| GO:0006950 | response to stress | 0.001259 | 0.001259325 |
| GO:0038094 | Fc-gamma receptor signaling pathway | 0.001263 | 0.001262965 |
| GO:0030593 | neutrophil chemotaxis | 0.001263 | 0.001262965 |
| GO:0045058 | T cell selection | 0.001263 | 0.001262965 |
| GO:0032609 | interferon-gamma production | 0.001263 | 0.001262965 |
| GO:0032649 | regulation of interferon-gamma production | 0.001263 | 0.001262965 |
| GO:0002218 | activation of innate immune response | 0.001366 | 0.001366144 |
| GO:0002532 | production of molecular mediator involved in inflammatory response | 0.001366 | 0.001366144 |
| GO:0080134 | regulation of response to stress | 0.001407 | 0.001406934 |
| GO:0045123 | cellular extravasation | 0.001468 | 0.001467829 |
| GO:0071222 | cellular response to lipopolysaccharide | 0.001546 | 0.001546387 |

**Supplemental Table 15. (Continued).**

| GO ID | GO Name | P Value | ADJ P Value |
|---|---|---|---|
| GO:1902533 | positive regulation of intracellular signal transduction | 0.001547 | 0.001547171 |
| GO:0071677 | positive regulation of mononuclear cell migration | 0.001551 | 0.001551313 |
| GO:0032733 | positive regulation of interleukin-10 production | 0.001551 | 0.001551313 |
| GO:0002526 | acute inflammatory response | 0.001551 | 0.001551313 |
| GO:0070588 | calcium ion transmembrane transport | 0.001602 | 0.001602101 |
| GO:0045006 | DNA deamination | 0.001725 | 0.001724872 |
| GO:0002673 | regulation of acute inflammatory response | 0.001728 | 0.001728104 |
| GO:0051239 | regulation of multicellular organismal process | 0.001746 | 0.001746288 |
| GO:0050789 | regulation of biological process | 0.001863 | 0.001862778 |
| GO:0002819 | regulation of adaptive immune response | 0.002011 | 0.002010754 |
| GO:0002705 | positive regulation of leukocyte mediated immunity | 0.002074 | 0.002074331 |
| GO:0071219 | cellular response to molecule of bacterial origin | 0.00217 | 0.002170104 |
| GO:0009967 | positive regulation of signal transduction | 0.002227 | 0.002226784 |
| GO:0002238 | response to molecule of fungal origin | 0.002259 | 0.00225885 |
| GO:0072676 | lymphocyte migration | 0.002259 | 0.00225885 |
| GO:0030183 | B cell differentiation | 0.002259 | 0.00225885 |
| GO:0071226 | cellular response to molecule of fungal origin | 0.002259 | 0.00225885 |
| GO:0002237 | response to molecule of bacterial origin | 0.002327 | 0.002327333 |
| GO:0045869 | negative regulation of single stranded viral RNA replication via double stranded DNA intermediate | 0.002534 | 0.002533743 |
| GO:0045088 | regulation of innate immune response | 0.002534 | 0.002533743 |
| GO:0002385 | mucosal immune response | 0.002534 | 0.002533743 |
| GO:0033077 | T cell differentiation in thymus | 0.002534 | 0.002533743 |
| GO:0045859 | regulation of protein kinase activity | 0.002698 | 0.002698387 |
| GO:0006816 | calcium ion transport | 0.002721 | 0.002720968 |
| GO:0031295 | T cell costimulation | 0.002776 | 0.002775759 |
| GO:0002366 | leukocyte activation involved in immune response | 0.002848 | 0.002847835 |
| GO:0002251 | organ or tissue specific immune response | 0.003046 | 0.003046143 |
| GO:0071621 | granulocyte chemotaxis | 0.003053 | 0.003053229 |
| GO:0002377 | immunoglobulin production | 0.003062 | 0.003062258 |
| GO:0010033 | response to organic substance | 0.003271 | 0.003271259 |
| GO:0002263 | cell activation involved in immune response | 0.003282 | 0.003282432 |
| GO:0031294 | lymphocyte costimulation | 0.00329 | 0.00328994 |
| GO:0007252 | I-kappaB phosphorylation | 0.003534 | 0.00353432 |
| GO:0098609 | cell-cell adhesion | 0.003796 | 0.003796477 |
| GO:0060326 | cell chemotaxis | 0.003911 | 0.003910682 |
| GO:0002437 | inflammatory response to antigenic stimulus | 0.003911 | 0.003910682 |
| GO:0043549 | regulation of kinase activity | 0.003975 | 0.003974909 |
| GO:0016553 | base conversion or substitution editing | 0.004052 | 0.004051782 |
| GO:0045091 | regulation of single stranded viral RNA replication via double stranded DNA intermediate | 0.004052 | 0.004051782 |
| GO:0007186 | G protein-coupled receptor signaling pathway | 0.004052 | 0.004051782 |
| GO:0002710 | negative regulation of T cell mediated immunity | 0.004052 | 0.004051782 |
| GO:0033630 | positive regulation of cell adhesion mediated by integrin | 0.004052 | 0.004051782 |
| GO:0071398 | cellular response to fatty acid | 0.004052 | 0.004051782 |
| GO:0001954 | positive regulation of cell-matrix adhesion | 0.004143 | 0.004142522 |
| GO:0002534 | cytokine production involved in inflammatory response | 0.004491 | 0.004490953 |
| GO:1900015 | regulation of cytokine production involved in inflammatory response | 0.004491 | 0.004490953 |
| GO:0039692 | single stranded viral RNA replication via double stranded DNA intermediate | 0.00464 | 0.004640311 |
| GO:0002698 | negative regulation of immune effector process | 0.00464 | 0.004640311 |
| GO:0050853 | B cell receptor signaling pathway | 0.00464 | 0.004640311 |
| GO:0002719 | negative regulation of cytokine production involved in immune response | 0.00464 | 0.004640311 |
| GO:1901222 | regulation of NIK/NF-kappaB signaling | 0.00486 | 0.00486036 |
| GO:0014065 | phosphatidylinositol 3-kinase signaling | 0.005023 | 0.005022521 |
| GO:0032651 | regulation of interleukin-1 beta production | 0.005085 | 0.005085136 |
| GO:0032611 | interleukin-1 beta production | 0.005085 | 0.005085136 |
| GO:0019722 | calcium-mediated signaling | 0.005259 | 0.005258622 |
| GO:0006935 | chemotaxis | 0.00539 | 0.005390484 |
| GO:0042330 | taxis | 0.00539 | 0.005390484 |
| GO:0070887 | cellular response to chemical stimulus | 0.005912 | 0.005911985 |
| GO:0030888 | regulation of B cell proliferation | 0.005959 | 0.005958639 |
| GO:0002706 | regulation of lymphocyte mediated immunity | 0.006009 | 0.006008889 |
| GO:0045577 | regulation of B cell differentiation | 0.006017 | 0.00601741 |
| GO:2000523 | regulation of T cell costimulation | 0.006155 | 0.006154972 |
| GO:0032496 | response to lipopolysaccharide | 0.006435 | 0.006435286 |
| GO:0010811 | positive regulation of cell-substrate adhesion | 0.006435 | 0.006435286 |

**Supplemental Table 15. (Continued).**

| GO ID | GO Name | P Value | ADJ P Value |
|---|---|---|---|
| GO:0010647 | positive regulation of cell communication | 0.006466 | 0.006465847 |
| GO:0032703 | negative regulation of interleukin-2 production | 0.006743 | 0.006743232 |
| GO:0002891 | positive regulation of immunoglobulin mediated immune response | 0.006743 | 0.006743232 |
| GO:0002714 | positive regulation of B cell mediated immunity | 0.006743 | 0.006743232 |
| GO:0023056 | positive regulation of signaling | 0.006762 | 0.006761765 |
| GO:0050777 | negative regulation of immune response | 0.0068 | 0.006800008 |
| GO:0048525 | negative regulation of viral process | 0.006959 | 0.006959196 |
| GO:1903169 | regulation of calcium ion transmembrane transport | 0.007022 | 0.007021788 |
| GO:0050854 | regulation of antigen receptor-mediated signaling pathway | 0.007121 | 0.007120548 |
| GO:0032102 | negative regulation of response to external stimulus | 0.007489 | 0.007488582 |
| GO:0006801 | superoxide metabolic process | 0.007609 | 0.007609318 |
| GO:0051924 | regulation of calcium ion transport | 0.007691 | 0.007691448 |
| GO:0014066 | regulation of phosphatidylinositol 3-kinase signaling | 0.007925 | 0.00792478 |
| GO:0032652 | regulation of interleukin-1 production | 0.008266 | 0.008265842 |
| GO:0032612 | interleukin-1 production | 0.008266 | 0.008265842 |
| GO:0061099 | negative regulation of protein tyrosine kinase activity | 0.008378 | 0.008378341 |
| GO:0050790 | regulation of catalytic activity | 0.008464 | 0.008464202 |
| GO:0002692 | negative regulation of cellular extravasation | 0.008464 | 0.008464202 |
| GO:1903721 | positive regulation of I-kappaB phosphorylation | 0.008464 | 0.008464202 |
| GO:0033634 | positive regulation of cell-cell adhesion mediated by integrin | 0.008464 | 0.008464202 |
| GO:0065007 | biological regulation | 0.00872 | 0.008720037 |
| GO:0016064 | immunoglobulin mediated immune response | 0.00872 | 0.008720037 |
| GO:0002832 | negative regulation of response to biotic stimulus | 0.008851 | 0.008851095 |
| GO:0045071 | negative regulation of viral genome replication | 0.008978 | 0.008978332 |
| GO:0002701 | negative regulation of production of molecular mediator of immune response | 0.009167 | 0.009166584 |
| GO:0019724 | B cell mediated immunity | 0.009947 | 0.009947069 |
| GO:0038096 | Fc-gamma receptor signaling pathway involved in phagocytosis | 0.010124 | 0.010123844 |
| GO:0032743 | positive regulation of interleukin-2 production | 0.010124 | 0.010123844 |
| GO:0050672 | negative regulation of lymphocyte proliferation | 0.010124 | 0.010123844 |
| GO:0002433 | immune response-regulating cell surface receptor signaling pathway involved in phagocytosis | 0.010124 | 0.010123844 |
| GO:0032945 | negative regulation of mononuclear cell proliferation | 0.010662 | 0.010661701 |
| GO:0072678 | T cell migration | 0.010662 | 0.010661701 |
| GO:0031666 | positive regulation of lipopolysaccharide-mediated signaling pathway | 0.010859 | 0.010859184 |
| GO:0035701 | hematopoietic stem cell migration | 0.010859 | 0.010859184 |
| GO:0002752 | cell surface pattern recognition receptor signaling pathway | 0.010859 | 0.010859184 |
| GO:0045619 | regulation of lymphocyte differentiation | 0.010859 | 0.010859184 |
| GO:0002732 | positive regulation of dendritic cell cytokine production | 0.010859 | 0.010859184 |
| GO:2000272 | negative regulation of signaling receptor activity | 0.010859 | 0.010859184 |
| GO:0010529 | negative regulation of transposition | 0.010859 | 0.010859184 |
| GO:0010528 | regulation of transposition | 0.010859 | 0.010859184 |
| GO:0070542 | response to fatty acid | 0.010859 | 0.010859184 |
| GO:1903719 | regulation of I-kappaB phosphorylation | 0.010859 | 0.010859184 |
| GO:0030595 | leukocyte chemotaxis | 0.010859 | 0.010859184 |
| GO:0008284 | positive regulation of cell population proliferation | 0.010859 | 0.010859184 |
| GO:0071675 | regulation of mononuclear cell migration | 0.011266 | 0.011265926 |
| GO:0038061 | NIK/NF-kappaB signaling | 0.011266 | 0.011265926 |
| GO:0042113 | B cell activation | 0.011266 | 0.011265926 |
| GO:0039694 | viral RNA genome replication | 0.011833 | 0.011832838 |
| GO:0050901 | leukocyte tethering or rolling | 0.011833 | 0.011832838 |
| GO:0048015 | phosphatidylinositol-mediated signaling | 0.012113 | 0.012112633 |
| GO:0043254 | regulation of protein-containing complex assembly | 0.012578 | 0.012577554 |
| GO:0048017 | inositol lipid-mediated signaling | 0.012864 | 0.012863661 |
| GO:2000406 | positive regulation of T cell migration | 0.012912 | 0.012911804 |
| GO:0080111 | DNA demethylation | 0.012912 | 0.012911804 |
| GO:0070664 | negative regulation of leukocyte proliferation | 0.013452 | 0.013451563 |
| GO:0042221 | response to chemical | 0.013978 | 0.013977709 |
| GO:0007204 | positive regulation of cytosolic calcium ion concentration | 0.013978 | 0.013977709 |
| GO:0051279 | regulation of release of sequestered calcium ion into cytosol | 0.013993 | 0.013993161 |
| GO:0150077 | regulation of neuroinflammatory response | 0.013993 | 0.013993161 |
| GO:0032196 | transposition | 0.013993 | 0.013993161 |
| GO:0032689 | negative regulation of interferon-gamma production | 0.013993 | 0.013993161 |
| GO:0016477 | cell migration | 0.014138 | 0.014137848 |
| GO:0072507 | divalent inorganic cation homeostasis | 0.014506 | 0.014505758 |
| GO:0030890 | positive regulation of B cell proliferation | 0.015167 | 0.015166606 |
| GO:0043507 | positive regulation of JUN kinase activity | 0.015167 | 0.015166606 |

**Supplemental Table 15. (Continued).**

| GO ID | GO Name | P Value | ADJ P Value |
|---|---|---|---|
| GO:0061097 | regulation of protein tyrosine kinase activity | 0.015436 | 0.015435761 |
| GO:1902105 | regulation of leukocyte differentiation | 0.015478 | 0.015477851 |
| GO:0050864 | regulation of B cell activation | 0.015762 | 0.015762353 |
| GO:0071356 | cellular response to tumor necrosis factor | 0.015762 | 0.015762353 |
| GO:0051338 | regulation of transferase activity | 0.016043 | 0.016043375 |
| GO:0045730 | respiratory burst | 0.01619 | 0.016189681 |
| GO:0002431 | Fc receptor mediated stimulatory signaling pathway | 0.01619 | 0.016189681 |
| GO:0042554 | superoxide anion generation | 0.01619 | 0.016189681 |
| GO:0050850 | positive regulation of calcium-mediated signaling | 0.01619 | 0.016189681 |
| GO:0006812 | cation transport | 0.016473 | 0.016473414 |
| GO:0002725 | negative regulation of T cell cytokine production | 0.016473 | 0.016473414 |
| GO:0002371 | dendritic cell cytokine production | 0.016473 | 0.016473414 |
| GO:0001771 | immunological synapse formation | 0.016473 | 0.016473414 |
| GO:0002730 | regulation of dendritic cell cytokine production | 0.016473 | 0.016473414 |
| GO:2001187 | positive regulation of CD8-positive, alpha-beta T cell activation | 0.016473 | 0.016473414 |
| GO:0002274 | myeloid leukocyte activation | 0.016676 | 0.01667619 |
| GO:0051345 | positive regulation of hydrolase activity | 0.017032 | 0.017031812 |
| GO:0006874 | cellular calcium ion homeostasis | 0.01719 | 0.017190273 |
| GO:0035510 | DNA dealkylation | 0.01719 | 0.017190273 |
| GO:0071900 | regulation of protein serine/threonine kinase activity | 0.018455 | 0.018455118 |
| GO:0032715 | negative regulation of interleukin-6 production | 0.018634 | 0.018634454 |
| GO:0046632 | alpha-beta T cell differentiation | 0.019694 | 0.01969355 |
| GO:0002687 | positive regulation of leukocyte migration | 0.019694 | 0.01969355 |
| GO:1905155 | positive regulation of membrane invagination | 0.019694 | 0.01969355 |
| GO:0050856 | regulation of T cell receptor signaling pathway | 0.019694 | 0.01969355 |
| GO:0034142 | toll-like receptor 4 signaling pathway | 0.019694 | 0.01969355 |
| GO:0033632 | regulation of cell-cell adhesion mediated by integrin | 0.019694 | 0.01969355 |
| GO:0060100 | positive regulation of phagocytosis, engulfment | 0.019694 | 0.01969355 |
| GO:1902622 | regulation of neutrophil migration | 0.019694 | 0.01969355 |
| GO:2000403 | positive regulation of lymphocyte migration | 0.019694 | 0.01969355 |
| GO:0002381 | immunoglobulin production involved in immunoglobulin-mediated immune response | 0.019694 | 0.01969355 |
| GO:0031348 | negative regulation of defense response | 0.020164 | 0.020163552 |
| GO:1903706 | regulation of hemopoiesis | 0.020164 | 0.020163552 |
| GO:0002823 | negative regulation of adaptive immune response based on somatic recombination of immune receptors built from immunoglobulin superfamily domains | 0.021179 | 0.021179088 |
| GO:0010469 | regulation of signaling receptor activity | 0.021888 | 0.02188832 |
| GO:0055074 | calcium ion homeostasis | 0.021888 | 0.02188832 |
| GO:0034612 | response to tumor necrosis factor | 0.022143 | 0.022143164 |
| GO:0048870 | cell motility | 0.022143 | 0.022143164 |
| GO:0030101 | natural killer cell activation | 0.022143 | 0.022143164 |
| GO:0002889 | regulation of immunoglobulin mediated immune response | 0.022536 | 0.022536238 |
| GO:0042100 | B cell proliferation | 0.023001 | 0.023000802 |
| GO:0034154 | toll-like receptor 7 signaling pathway | 0.023074 | 0.023073529 |
| GO:0042116 | macrophage activation | 0.02398 | 0.023979626 |
| GO:0002712 | regulation of B cell mediated immunity | 0.023996 | 0.023995791 |
| GO:0002707 | negative regulation of lymphocyte mediated immunity | 0.023996 | 0.023995791 |
| GO:0050871 | positive regulation of B cell activation | 0.024457 | 0.024457447 |
| GO:2000404 | regulation of T cell migration | 0.025704 | 0.025703611 |
| GO:0002822 | regulation of adaptive immune response based on somatic recombination of immune receptors built from immunoglobulin superfamily domains | 0.026016 | 0.026016257 |
| GO:0018108 | peptidyl-tyrosine phosphorylation | 0.026467 | 0.026467035 |
| GO:0034695 | response to prostaglandin E | 0.02664 | 0.026639653 |
| GO:0060099 | regulation of phagocytosis, engulfment | 0.02664 | 0.026639653 |
| GO:0071346 | cellular response to interferon-gamma | 0.02664 | 0.026639653 |
| GO:1905153 | regulation of membrane invagination | 0.02664 | 0.026639653 |
| GO:0001867 | complement activation, lectin pathway | 0.02664 | 0.026639653 |
| GO:0001779 | natural killer cell differentiation | 0.02664 | 0.026639653 |
| GO:0043277 | apoptotic cell clearance | 0.026973 | 0.026973093 |
| GO:0018212 | peptidyl-tyrosine modification | 0.027082 | 0.027081775 |
| GO:0043269 | regulation of ion transport | 0.027416 | 0.0274162 |
| GO:0002695 | negative regulation of leukocyte activation | 0.027886 | 0.027885725 |
| GO:0007155 | cell adhesion | 0.028279 | 0.028279444 |
| GO:0046634 | regulation of alpha-beta T cell activation | 0.028279 | 0.028279444 |
| GO:0009620 | response to fungus | 0.028279 | 0.028279444 |
| GO:0002820 | negative regulation of adaptive immune response | 0.028279 | 0.028279444 |
| GO:0033628 | regulation of cell adhesion mediated by integrin | 0.028279 | 0.028279444 |

**Supplemental Table 15. (Continued).**

| GO ID | GO Name | P Value | ADJ P Value |
|---|---|---|---|
| GO:0045069 | regulation of viral genome replication | 0.028279 | 0.028279444 |
| GO:0043124 | negative regulation of I-kappaB kinase/NF-kappaB signaling | 0.028279 | 0.028279444 |
| GO:0019932 | second-messenger-mediated signaling | 0.028785 | 0.028784641 |
| GO:1903900 | regulation of viral life cycle | 0.029052 | 0.029052059 |
| GO:0002821 | positive regulation of adaptive immune response | 0.029229 | 0.029228647 |
| GO:0031341 | regulation of cell killing | 0.029229 | 0.029228647 |
| GO:2000010 | positive regulation of protein localization to cell surface | 0.029564 | 0.029563665 |
| GO:1903428 | positive regulation of reactive oxygen species biosynthetic process | 0.029564 | 0.029563665 |
| GO:0002281 | macrophage activation involved in immune response | 0.029564 | 0.029563665 |
| GO:0002438 | acute inflammatory response to antigenic stimulus | 0.029564 | 0.029563665 |
| GO:0150078 | positive regulation of neuroinflammatory response | 0.029564 | 0.029563665 |
| GO:0031665 | negative regulation of lipopolysaccharide-mediated signaling pathway | 0.029564 | 0.029563665 |
| GO:0061756 | leukocyte adhesion to vascular endothelial cell | 0.029564 | 0.029563665 |
| GO:0002708 | positive regulation of lymphocyte mediated immunity | 0.029822 | 0.029821865 |
| GO:0002456 | T cell mediated immunity | 0.029822 | 0.029821865 |
| GO:0051090 | regulation of DNA-binding transcription factor activity | 0.030156 | 0.030156343 |
| GO:0072503 | cellular divalent inorganic cation homeostasis | 0.0309 | 0.030900223 |
| GO:0001932 | regulation of protein phosphorylation | 0.032852 | 0.032852444 |
| GO:0062208 | positive regulation of pattern recognition receptor signaling pathway | 0.032901 | 0.032901349 |
| GO:0043506 | regulation of JUN kinase activity | 0.032901 | 0.032901349 |
| GO:0050730 | regulation of peptidyl-tyrosine phosphorylation | 0.033073 | 0.033073105 |
| GO:0010820 | positive regulation of T cell chemotaxis | 0.033396 | 0.033396178 |
| GO:0032725 | positive regulation of granulocyte macrophage colony-stimulating factor production | 0.033396 | 0.033396178 |
| GO:0002283 | neutrophil activation involved in immune response | 0.033396 | 0.033396178 |
| GO:0051403 | stress-activated MAPK cascade | 0.033557 | 0.033557327 |
| GO:0002704 | negative regulation of leukocyte mediated immunity | 0.034456 | 0.034455812 |
| GO:0050732 | negative regulation of peptidyl-tyrosine phosphorylation | 0.034456 | 0.034455812 |
| GO:0043406 | positive regulation of MAP kinase activity | 0.035444 | 0.035443875 |
| GO:0065009 | regulation of molecular function | 0.036371 | 0.036370684 |
| GO:0042130 | negative regulation of T cell proliferation | 0.036381 | 0.036381229 |
| GO:0031098 | stress-activated protein kinase signaling cascade | 0.036388 | 0.036387884 |
| GO:1901701 | cellular response to oxygen-containing compound | 0.036632 | 0.036631881 |
| GO:0034694 | response to prostaglandin | 0.037076 | 0.037075732 |
| GO:0034116 | positive regulation of heterotypic cell-cell adhesion | 0.037076 | 0.037075732 |
| GO:0010819 | regulation of T cell chemotaxis | 0.037076 | 0.037075732 |
| GO:0034134 | toll-like receptor 2 signaling pathway | 0.037076 | 0.037075732 |
| GO:0033631 | cell-cell adhesion mediated by integrin | 0.037076 | 0.037075732 |
| GO:0040011 | locomotion | 0.037076 | 0.037075732 |
| GO:0062207 | regulation of pattern recognition receptor signaling pathway | 0.038791 | 0.038791444 |
| GO:0002285 | lymphocyte activation involved in immune response | 0.039305 | 0.039305427 |
| GO:0032645 | regulation of granulocyte macrophage colony-stimulating factor production | 0.041185 | 0.041185212 |
| GO:0061154 | endothelial tube morphogenesis | 0.041185 | 0.041185212 |
| GO:0032604 | granulocyte macrophage colony-stimulating factor production | 0.041185 | 0.041185212 |
| GO:0051092 | positive regulation of NF-kappaB transcription factor activity | 0.041185 | 0.041185212 |
| GO:1900221 | regulation of amyloid-beta clearance | 0.041185 | 0.041185212 |
| GO:0098655 | cation transmembrane transport | 0.041185 | 0.041185212 |
| GO:0003159 | morphogenesis of an endothelium | 0.041185 | 0.041185212 |
| GO:0050866 | negative regulation of cell activation | 0.041185 | 0.041185212 |
| GO:0035746 | granzyme A production | 0.042275 | 0.04227481 |
| GO:0002669 | positive regulation of T cell anergy | 0.042275 | 0.04227481 |
| GO:0002325 | natural killer cell differentiation involved in immune response | 0.042275 | 0.04227481 |
| GO:2000334 | positive regulation of blood microparticle formation | 0.042275 | 0.04227481 |
| GO:0045584 | negative regulation of cytotoxic T cell differentiation | 0.042275 | 0.04227481 |
| GO:0150129 | positive regulation of interleukin-33 production | 0.042275 | 0.04227481 |
| GO:0072682 | eosinophil extravasation | 0.042275 | 0.04227481 |
| GO:0035782 | mature natural killer cell chemotaxis | 0.042275 | 0.04227481 |
| GO:2000332 | regulation of blood microparticle formation | 0.042275 | 0.04227481 |
| GO:0097534 | lymphoid lineage cell migration | 0.042275 | 0.04227481 |
| GO:0032826 | regulation of natural killer cell differentiation involved in immune response | 0.042275 | 0.04227481 |
| GO:0002442 | serotonin secretion involved in inflammatory response | 0.042275 | 0.04227481 |
| GO:0097535 | lymphoid lineage cell migration into thymus | 0.042275 | 0.04227481 |
| GO:0042325 | regulation of phosphorylation | 0.042275 | 0.04227481 |
| GO:0090721 | primary adaptive immune response involving T cells and B cells | 0.042275 | 0.04227481 |
| GO:2000526 | positive regulation of glycoprotein biosynthetic process involved in immunological synapse formation | 0.042275 | 0.04227481 |
| GO:0090720 | primary adaptive immune response | 0.042275 | 0.04227481 |

**Supplemental Table 15. (Continued).**

| GO ID | GO Name | P Value | ADJ P Value |
|---|---|---|---|
| GO:0002554 | serotonin secretion by platelet | 0.042275 | 0.04227481 |
| GO:2000513 | positive regulation of granzyme A production | 0.042275 | 0.04227481 |
| GO:2000511 | regulation of granzyme A production | 0.042275 | 0.04227481 |
| GO:2000517 | regulation of T-helper 1 cell activation | 0.042275 | 0.04227481 |
| GO:0042543 | protein N-linked glycosylation via arginine | 0.042275 | 0.04227481 |
| GO:0038123 | toll-like receptor TLR1:TLR2 signaling pathway | 0.042275 | 0.04227481 |
| GO:0002913 | positive regulation of lymphocyte anergy | 0.042275 | 0.04227481 |
| GO:2000420 | negative regulation of eosinophil extravasation | 0.042275 | 0.04227481 |
| GO:2000419 | regulation of eosinophil extravasation | 0.042275 | 0.04227481 |
| GO:0002439 | chronic inflammatory response to antigenic stimulus | 0.042275 | 0.04227481 |
| GO:2000518 | negative regulation of T-helper 1 cell activation | 0.042275 | 0.04227481 |
| GO:0014895 | smooth muscle hypertrophy | 0.042275 | 0.04227481 |
| GO:0072564 | blood microparticle formation | 0.042275 | 0.04227481 |
| GO:0048298 | positive regulation of isotype switching to IgA isotypes | 0.042275 | 0.04227481 |
| GO:0002351 | serotonin production involved in inflammatory response | 0.042275 | 0.04227481 |
| GO:0150127 | regulation of interleukin-33 production | 0.042275 | 0.04227481 |
| GO:0072639 | interleukin-33 production | 0.042275 | 0.04227481 |
| GO:0061048 | negative regulation of branching involved in lung morphogenesis | 0.042275 | 0.04227481 |
| GO:1901700 | response to oxygen-containing compound | 0.042275 | 0.04227481 |
| GO:0010604 | positive regulation of macromolecule metabolic process | 0.042333 | 0.042332873 |
| GO:1900017 | positive regulation of cytokine production involved in inflammatory response | 0.042404 | 0.042404035 |
| GO:0033623 | regulation of integrin activation | 0.042404 | 0.042404035 |
| GO:0034162 | toll-like receptor 9 signaling pathway | 0.042404 | 0.042404035 |
| GO:0046635 | positive regulation of alpha-beta T cell activation | 0.042404 | 0.042404035 |
| GO:0032753 | positive regulation of interleukin-4 production | 0.042404 | 0.042404035 |
| GO:0050792 | regulation of viral process | 0.042598 | 0.042597594 |
| GO:0001952 | regulation of cell-matrix adhesion | 0.044537 | 0.044537257 |
| GO:2001185 | regulation of CD8-positive, alpha-beta T cell activation | 0.046537 | 0.046537297 |
| GO:2000379 | positive regulation of reactive oxygen species metabolic process | 0.046537 | 0.046537297 |
| GO:0150076 | neuroinflammatory response | 0.046537 | 0.046537297 |
| GO:1900225 | regulation of NLRP3 inflammasome complex assembly | 0.046537 | 0.046537297 |
| GO:0046641 | positive regulation of alpha-beta T cell proliferation | 0.046537 | 0.046537297 |
| GO:0043652 | engulfment of apoptotic cell | 0.046537 | 0.046537297 |
| GO:0030050 | vesicle transport along actin filament | 0.046537 | 0.046537297 |
| GO:0034341 | response to interferon-gamma | 0.046932 | 0.046932004 |
| GO:0030001 | metal ion transport | 0.047045 | 0.04704543 |
| GO:0051336 | regulation of hydrolase activity | 0.047944 | 0.047943669 |
| GO:1903038 | negative regulation of leukocyte cell-cell adhesion | 0.048216 | 0.048216447 |
| GO:0031343 | positive regulation of cell killing | 0.048216 | 0.048216447 |
| GO:1904062 | regulation of cation transmembrane transport | 0.048851 | 0.048851314 |
| GO:0042127 | regulation of cell population proliferation | 0.049172 | 0.04917162 |
